# Aquarius RNA helicase Protects Pluripotent Stem Cell Identity

**DOI:** 10.64898/2026.04.24.720612

**Authors:** Maxime Lalonde, Elizabeth Márquez-Gómez, Clare S.K. Lee, Adam Burton, Ioannis Tsirkas, Fernanda Rezende Pabst, Marcel Werner, Atiqa Sajid, Augusto Chaves Murriello, Xanthoula Karypidou, Andreas Ettinger, Tamas Schauer, Tobias Straub, Maria-Elena Torres-Padilla, Antonio Scialdone, Stephan Hamperl

**Affiliations:** Institute of Epigenetics and Stem Cells (IES), Helmholtz Munich, Munich, Germany; Core Facility Bioinformatics, Biomedizinisches Centrum (BMC) LMU Munich, Germany

**Keywords:** Embryonic Stem Cells, Pluripotency, Transcription-Replication Conflicts, Aquarius, R-loop Resolution, Transcription fidelity, Genomic Instability, Cell Fate Plasticity, Single-Cell Microscopy

## Abstract

Pluripotent stem cells must reconcile rapid replication with a highly dynamic transcriptional program, creating an inherent susceptibility to transcription–replication conflicts (TRCs). We demonstrate that embryonic stem cells (ESCs) operate in a “resilient” replication mode, tolerating high genomic traffic through the constitutive upregulation of R-loop and TRC resolution pathways. Through a targeted functional screen, we identify the RNA helicase Aquarius (AQR) as an essential safeguard of this state. AQR depletion downregulates these resolution factors, collapsing this stress-resistant program and driving ESCs into unstable, heterogenous states, marked by increased transcriptional entropy and cell-to-cell noise. Mechanistically, we show that key identity-defining genes are preferentially located within R-loop and TRC prone regions, making them uniquely vulnerable to AQR depletion. Our findings establish AQR as a critical governor of transcriptional fidelity, demonstrating that genomic resilience is fundamental to maintaining pluripotent cell identity.

## Introduction

Embryonic stem cells (ESCs) display a dynamic epigenome which is linked to its pluripotency features. Differentiation of ESCs relies on transcriptional rewiring and involves reorganization of DNA replication timing, 3D genome organization and the epigenome to steer and enforce cell identity transitions^1^. In mouse ESC (mESC) cultures, a small transient subpopulation of totipotent-like cells arise that resemble cells of the two-cell (2C) stage of early mouse embryos. These 2-cell–like cells (2CLCs) exhibit a distinctive transcriptional program and chromatin landscape, characterized by activation of MERVL retrotransposons and 2C-specific genes. Crucially, recent studies have demonstrated that the induction of replication stress (RS) and the subsequent slowing of replication forks can act as a primary driver for the emergence of these totipotent-like cells^2^.

A frequent endogenous obstacle to replication forks that leads to RS are transcribing RNA polymerases^3^. The lack of coordination between transcription and replication can lead to transcription-replication conflicts (TRCs), a hazardous genomic event resulting in fork stalling, DNA breakage, and mutations, making the management of genomic “traffic” a central problem of cell biology^3–5^. R-loops, RNA:DNA hybrid structures with a displaced single-stranded DNA, often form during transcription and can exacerbate these conflicts by creating physical barriers to replication fork progression^3,6^. While R-loops can serve physiological roles in transcription regulation and genome organization, their persistence can hinder replication fork progression and promote genome instability^7^. Although cells possess multiple layers of spatial and temporal regulation to minimize R-loops and TRCs, the proliferative features of mESCs, including an unusually short G1-phase^8^, a high density of replication origins, and an open, accessible chromatin structure^9–12^, fundamentally challenge known safeguarding mechanisms against TRCs. Given that R-loops and TRCs were shown to evict protective nucleosomes and disrupt local gene expression profiles^13^, a fundamental question is how mESCs accommodate this intense ‘genomic traffic’ while maintaining genome integrity and the pluripotency transcription program. More broadly, this raises the question of whether different cell identities employ distinct strategies to coordinate transcription and DNA replication.

In this study, we demonstrate that unperturbed mESCs adopt a unique ‘resilient’ replication strategy characterized by an unconventional spatial overlap between transcription and replication activities. Unlike differentiated cells that strictly segregate these processes, pluripotent cells tolerate high genomic traffic through the constitutive upregulation of specialized resolution pathways. Through a targeted functional screen, we identify the RNA helicase Aquarius (AQR) as an essential safeguard of this resilient state. We show that AQR depletion collapses the coordination between transcription and replication (T-R coordination), triggering R-loop mediated replication stress and an erosion of pluripotent identity. Single-cell transcriptomics reveal that this loss of identity is driven by a surge in transcriptional entropy and cell-to-cell variance, reflecting a systemic failure in regulatory constraint and enhancing the potency of these cells. Mechanistically, we find that key identity-defining pluripotency genes are preferentially enriched within genomic ’hotspots’ prone to R-loop-dependent conflicts, rendering the core transcriptional network intrinsically vulnerable to stochastic interference. Together, our findings establish AQR as a critical governor of transcriptional fidelity and reveal that the active management of genomic traffic is a fundamental requirement for maintaining the stability of the pluripotent state.

## Results

### Pluripotency is characterized by high replication activity and high transcription-replication overlap

To understand how pluripotent cells manage the challenge of combining rapid replication with a highly dynamic transcription program, we first compared global replication and transcription dynamics in mESCs, 2CLCs and mouse embryonic fibroblasts (MEFs). In comparison to fast-cycling, pluripotent mESCs, 2CLCs offer a model of reprogramming to a totipotent-like cell state with an altered transcription program, whereas MEFs represent a more differentiated, transcriptionally stable and slow-cycling multipotent state. To identify 2CLCs within the mESC populations, we used a MERVL::tbGFP mESC reporter line, where turbo-GFP (tbGFP) expression selectively marks the 2C-like state ^14,2^. Using quantitative high-throughput imaging, we simultaneously monitored active DNA synthesis (EdU incorporation) and elongating RNA polymerase II (C-terminal domain (CTD) phosphorylated at Ser2, PIIpS2)^15^ (**Fig. 1A**). Consistent with their short G1 phase, mESCs exhibited a vastly expanded S-phase population (65%) compared to both 2CLCs (33%) and MEFs (17%) (**Fig. 1B**). Additionally, the level of EdU incorporation in S-phase cells was significantly higher in mESCs than in either MEFs or 2CLCs, consistent with their high replicative activity^16^ (**Fig. 1C**).

**Figure 1.**
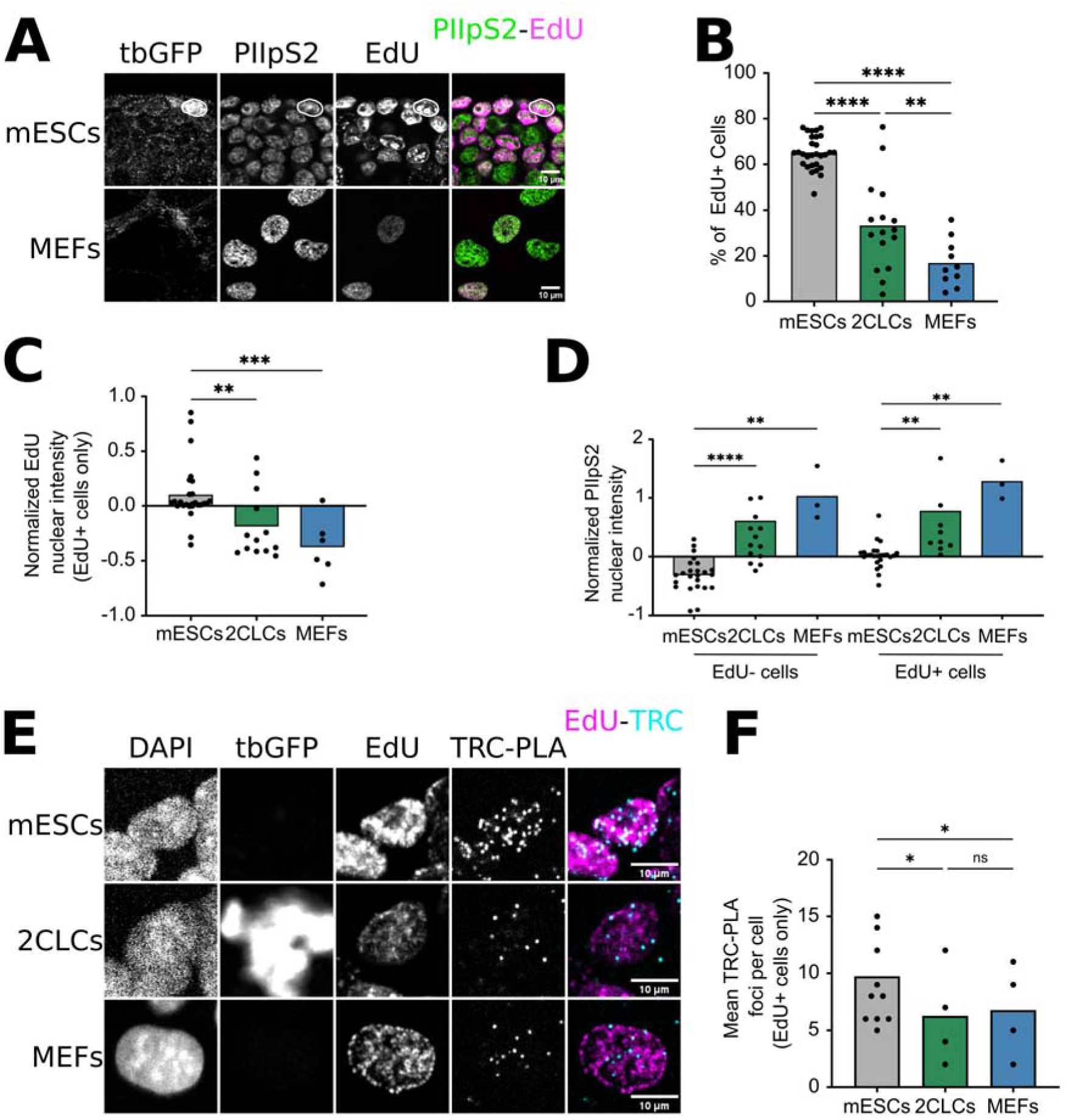
mESCs exhibit higher replication rates and higher TRC burden compared to MEFs and 2CLCs. (**A**) Representative immunofluorescence image of mESCs, 2CLCs (top), and MEFs (bottom). 2CLCs (circled) were identified by expression of tbGFP. Active transcription was assessed by chromatin-bound PIIpS2, and active DNA replication was measured by EdU incorporation. (**B**) Quantification of EdU-positive cells across cell types. Each dot represents one biological replicate (N = 10–50), calculated as the mean of ≥3 technical replicates, except for 2CLCs (200–10,000 analyzed mESCs or MEFs per technical replicate). For 2CLCs, each biological replicate corresponds to the mean of all 2CLCs within that biological replicate (≥50 2CLCs per replicate). Statistical significance was determined by one-way ANOVA followed by Bonferroni’s multiple comparisons test. (**C**) Quantification of EdU incorporation in EdU-positive cells across cell types. EdU incorporation was normalized replicate-wise through robust Z score transformation. Each dot represents one biological replicate (N = 6-27), calculated as the mean of ≥3 technical replicates, except for 2CLCs (200–10,000 analyzed mESCs or MEFs per technical replicate). For 2CLCs, each biological replicate corresponds to the mean of all 2CLCs within that biological replicate (≥30 2CLCs per replicate). Statistical significance was determined by one-way ANOVA followed by Bonferroni’s multiple comparisons test. (**D**) Quantification of chromatin-bound PIIpS2 across cell types and in EdU-negative cells (left) and EdU-positive cells (right). PIIpS2 was normalized replicate-wise through robust Z score transformation. Each dot represents one biological replicate (N = 3-24), calculated as the mean of ≥3 technical replicates, except for 2CLCs (200–10,000 analyzed mESCs or MEFs per technical replicate). For 2CLCs, each biological replicate corresponds to the mean of all 2CLCs within that biological replicate (≥30 2CLCs per replicate). Statistical significance was determined by one-way ANOVA followed by Bonferroni’s multiple comparisons test. (**E**) Representative TRC-PLA microscopy image of mESC (top), 2CLC (middle), and MEF (bottom). 2CLCs were identified by expression of tbGFP. S-phase cells were identified through a short EdU pulse and Click-it reaction. TRC-PLA foci were detected using an antibody against PCNA and an antibody against PIIpS2 in pre-extracted cells. (**F**) Quantification of the number of TRC-PLA foci per nuclei across cell types in EdU-positive cells. Each dot represents one biological replicate (N = 4-11), calculated as the mean of ≥3 technical replicates, except for 2CLCs (100-3,000 analyzed mESCs or MEFs per technical replicate). For 2CLCs, each biological replicate corresponds to the mean of all 2CLCs within that biological replicate (≥30 2CLCs per replicate). Statistical significance was determined by paired one-way ANOVA followed by Bonferroni’s multiple comparisons test.

Interestingly, mESCs displayed lower levels of chromatin-bound PIIpS2 than either 2CLCs or MEFs (**Fig. 1D**). This difference was also observed when using complementary transcription readouts such as EU incorporation and total non-phosphorylated RNAPII levels (**Fig. S1A, B**) and persisted across all cell cycle phases (**Fig. 1D**). This suggests that mESCs adopt a global transcriptional state characterized by reduced RNAPII occupancy when compared to MEF cells, rather than a previously described hyper-transcriptional state^17,18^. Altogether, these data suggest that there is a striking plasticity of transcription and replication activity between different cell types, with pluripotent cells adopting a unique ‘high-traffic’ genomic signature that balances maximal replication activity with a distinct, moderate transcriptional program.

To determine if the high replication activity of mESCs translates into increased physical collisions, we first quantified TRC levels using the well-established proximity ligation assay (TRC-PLA) between PIIpS2 and PCNA. Despite their lower global RNAPII occupancy (**Fig. 1D**), mESCs exhibited significantly higher TRC-PLA foci compared to 2CLCs or MEFs (**Fig. 1E, F**). Taken together, this suggests that the elevated TRC burden is not driven by higher global transcription, but rather by their high replication rates and potentially by the resulting poor spatial coordination between the two machineries.

To quantify this coordination globally, we developed an imaging-based framework to calculate the Manders’ correlation coefficient (MCC) between PIIpS2 and EdU signals (PIIpS2-EdU MCC), which quantifies global T-R coordination, here defined as the percentage of transcriptionally active regions of the nucleus overlapping with DNA synthesis (T-R overlap) (**Fig. 2A-B**, see also Methods and **Fig. S2**).

**Figure 2.**
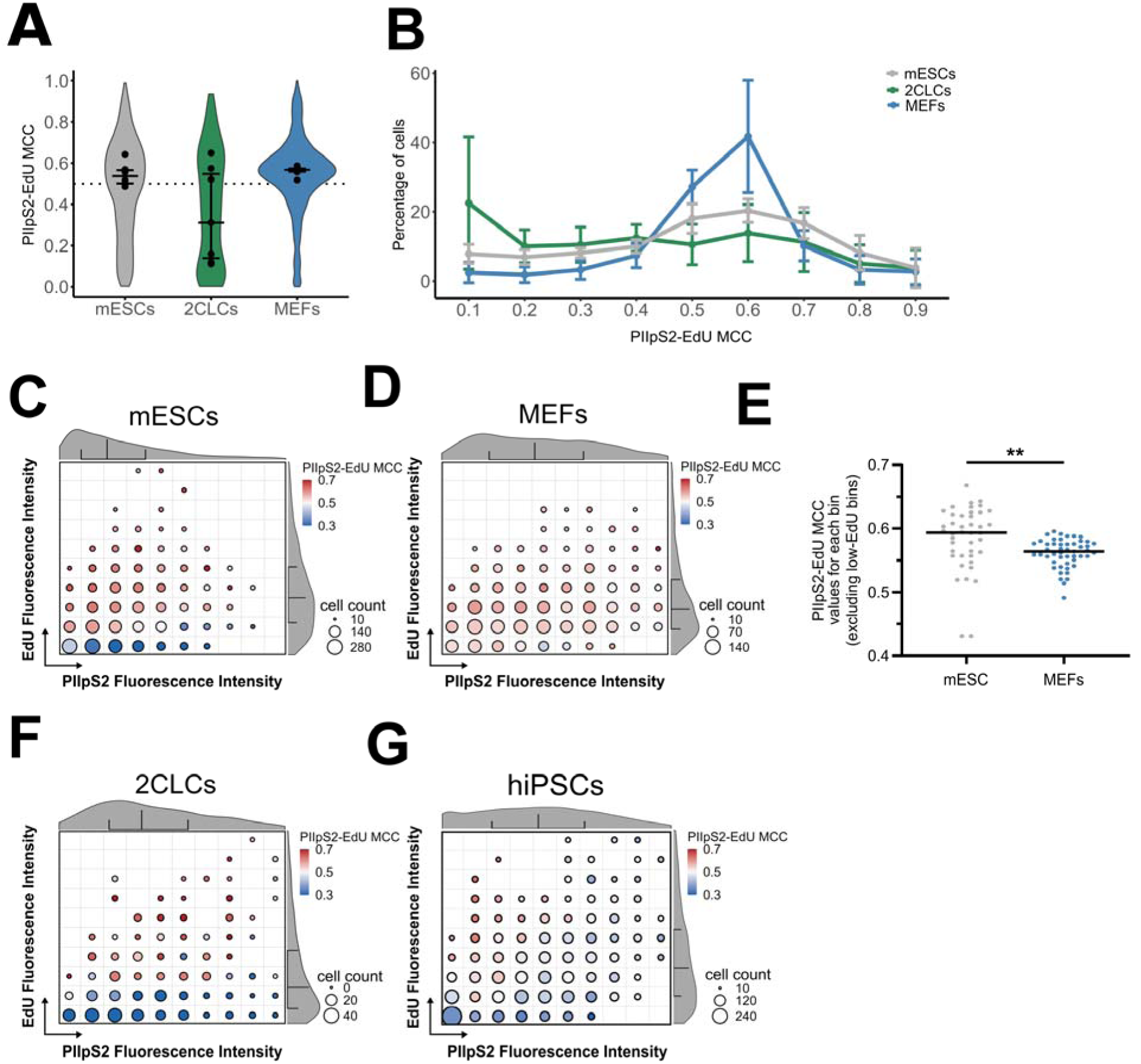
mESCs exhibit reduced transcription–replication coordination. **(A-B)** Quantification of transcription–replication coordination using PIIpS2-EdU MCC across cell types. Violin plots (**A**) and frequency distributions (**B**) show MCC distributions. Dots in the violin plots represent the mean value of biological replicates with the error bars representing the standard deviation amongst the replicates, the violin plot shows the distribution of all biological replicates pooled together. The frequency distributions show the mean and standard deviation of all biological replicates for each bin of MCC. Each cell type has been down sampled to the condition with the lowest number of cells for each independent experiment. N = 5 (7 for 2CLCs), comprising 464 2CLCs, 4097 mESCs, and 3178 MEFs. **(C)** Bubble plot of mESCs T-R overlap as quantified by the PIIpS2–EdU MCC. Values are stratified by PIIpS2 intensity (X-axis) and EdU intensity (Y-axis) using the same dataset as in (A) (see “Image analysis” in the methods for further information on the data processing). Bubble size indicates the number of cells per bin, while bubble color represents the median PIIpS2-EdU MCC for that bin. Parallel to the X-axis and the Y-axis are violin plots of the PIIpS2 and EdU signals, respectively, representing the whole population. **(D)** Bubble plot of MEFs T-R overlap as quantified by the PIIpS2–EdU MCC, as in (C). (**E**) Bin-wise comparison of MCC values from mESCs and MEFs (C and D). MCC values from all bins defined in panel (C and D) were extracted, excluding bins with low EdU signal (rows 1 and 2) to avoid regions with minimal replication activity. Statistical significance of the difference in MCC between cell types was assessed using a two-tailed paired t-test across all included bins. Each dot represents the MCC value of a single bin. **(F)** Bubble plot of 2CLCs T-R overlap as quantified by the PIIpS2–EdU MCC, as in (C). **(G)** Bubble plot of hiPSCs T-R overlap as quantified by the PIIpS2–EdU MCC, as in (C).

While MEFs maintain a narrow, homogeneous T-R coordination amongst the population (MCC 0.45-0.65), mESCs displayed a broader and more heterogeneous distribution, with an increased proportion of cells skewed towards higher MCC values (**Fig. 2A-B**). In contrast, 2CLCs exhibit a bimodal distribution: one population showing low MCC values, consistent with high segregation, while the other showed high MCC values, similar to mESCs with low segregation (**Fig. 2A-B**). Notably, when we stratified mESCs cells by their nuclear EdU pattern into early, mid or late S-phase cells^19^, we found that MCC values progressively decreases as cells transition toward late S-phase, yet mESCs maintain a characteristically higher overlap during early replication compared to 2CLCs, with significant cell-to-cell heterogeneity persisting across all cell cycle stages (**Fig. S1C, D**). To further disentangle the contributions of transcriptional and replicative activity to T-R coordination, we normalized and stratified cells in ten distinct bins from low to high transcription and replication activities (**Fig. 2C, D, F**). mESCs consistently exhibit higher MCC values than MEFs with matched transcription and replication activities (**Fig. 2E**). This implies that the poor segregation of transcription and replication in mESCs is a fundamental feature of these pluripotent cells. Consistently, we found that this replication-dependent “high-overlap” T-R profile is shared by human induced pluripotent stem cells (hiPSCs) (**Fig. 2G**), suggesting it is a conserved signature of pluripotency.

### High transcription-replication overlap is a developmentally regulated hallmark of pluripotency

We next asked whether this “high-conflict” environment is an inherent constraint related to pluripotency that is alleviated upon differentiation. To test this, we monitored TRC levels and T-R coordination as mESCs transitioned from naive ground-state pluripotency toward either primed pluripotency (Epiblast-like cells, EpiLCs) or early differentiation (-Lif) (**Fig. 3A, Fig. S3A**). Notably, high TRC levels, measured by the TRC-PLA assay, were maintained during the conversion to the primed state, suggesting that a high-conflict genomic environment is a shared hallmark of both naive and primed pluripotency. In contrast, TRC levels declined rapidly upon the onset of differentiation (**Fig. 3B**).

**Figure 3.**
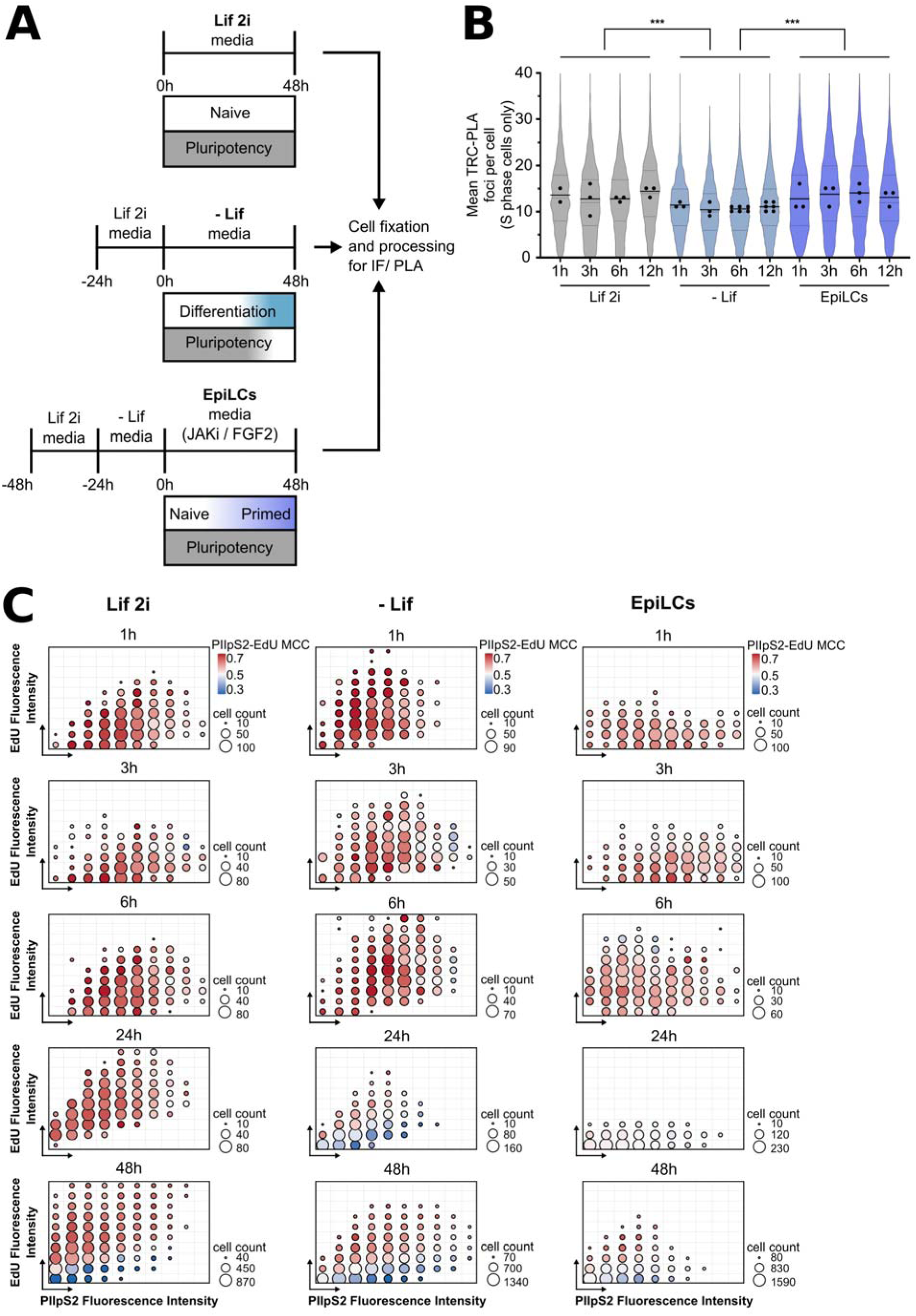
Transcription–replication conflicts and coordination dynamics during mESC cell fate transitions. (**A**) Experimental design for monitoring mESC fate changes: maintained naive state (left), pluripotency loss (middle), and naive-to-primed transition (right). (**B**) Quantification of TRC-PLA foci per nucleus in EdU-positive mESCs at early transition timepoints. Each dot represents one biological replicate (N = 2-6), calculated as the mean of ≥3 technical replicates (100–2,500 analyzed cells per technical replicate). Statistical significance was determined by one-way ANOVA followed by Bonferroni’s multiple comparisons test. (**C**) PIIpS2–EdU MCC bubble plots illustrating dynamic changes in coordination during transition. Cells are binned by PIIpS2 intensity (X-axis) and EdU intensity (Y-axis) (see “Image analysis” in the methods for further information on the data processing); bubble size reflects cell number, and bubble color encodes the median MCC per bin.

Quantitative time course analysis using the PIIpS2-EdU MCC readout revealed that the T-R coordination adapted according to a stepwise process. In the earlier timepoints, cells undergoing differentiation or priming display an increase in the amount of PIIpS2 on the chromatin (**Fig. 3C** (1 h vs 3 h and 6 h, changes along the X axis)), indicative of cells adjusting their transcription program as a response to the changed medium composition. This initial rewiring is followed, within one to two cell cycles, by a significant reduction in replication activity (**Fig. 3C** (1 h vs 24 h and 48 h, changes along the Y axis) and a corresponding improvement in T-R coordination (**Fig. 3C**, changes in MCC values**, Fig. S4B**).

Together, these findings establish that high genomic traffic and limited T-R coordination are intrinsic to pluripotency, creating a high-risk environment for TRCs that must be actively managed to sustain cell identity.

### mESCs show high expression of R-loop and TRC resolution regulators

Despite the high replication activity and a high proportion of cells with low T-R coordination, mESCs exhibit only a modest increase in TRC-PLA foci compared to 2CLCs and MEFs (**Fig. 1E, F**), suggesting they possess an enhanced capacity to resolve these conflicts. We hypothesized that the pluripotent state is supported by a specialized “resilient” replication program focused on resolving conflicts instead of avoiding them. To test this, we used publicly available RNA-seq datasets^20,21^ and performed differential gene expression analysis across naive mESCs, primed EpiLCs, differentiated MEFs, and totipotent-like 2CLCs, focusing on curated factors implicated in TRC resolution (123 genes, **Supplementary Table 1**)^22,23^, R-loop regulators (417 genes, **Supplementary Table 2**)^24^, DNA damage and repair (DDR) (234 genes, **Supplementary Table 3**) as well as chromatin and epigenetic regulators (752 genes, **Supplementary Table 4**).

Our analysis revealed a striking upregulation of known “conflict resolution” pathways in mESCs compared to differentiated MEFs (**Supplementary Table 5**), including the majority of TRC resolution genes (72 up, 10 down), R-loop regulatory factors (245 up, 48 down), and DDR-associated genes (123 up, 36 down) (**Fig. 4A**), supporting the notion that mESCs intrinsically express higher levels of these factors, potentially enabling more efficient TRC detection and resolution. Furthermore, mESCs showed a significant upregulation of chromatin and epigenetic regulators (386 up, 177 down) (**Fig. 4A**), likely reflecting an enhanced ability to modulate chromatin-bound processes at sites of interference.

**Figure 4.**
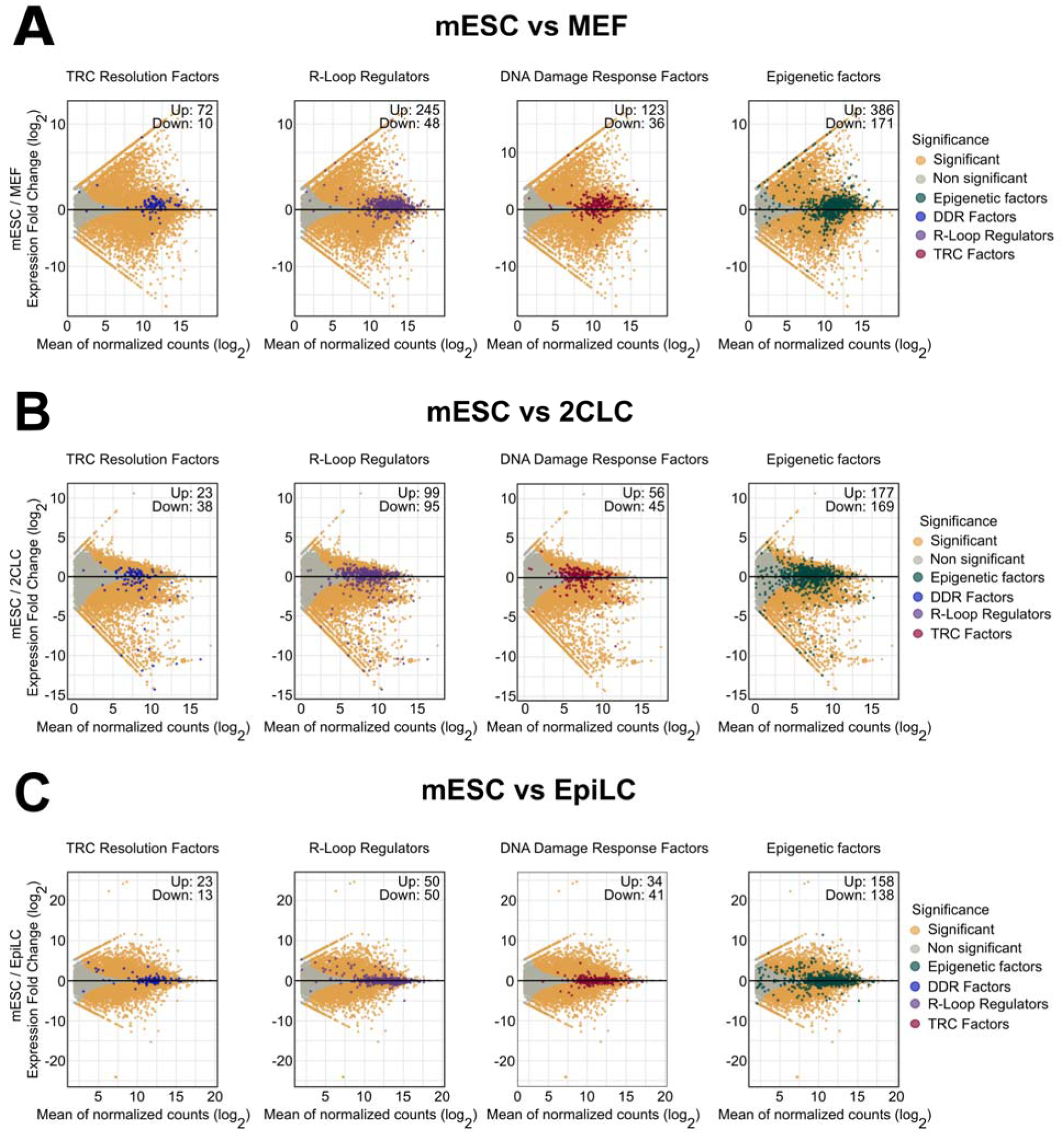
A Pluripotency-Associated ‘Resilient Replication’ Signature Enriches Conflict-Resolution Pathways. **(A-C)** MA plots of differentially expressed genes (DEGs) between mESCs and MEFs (**A**), between mESCs and 2CLCs (**B**), or between naive mESCs and primed mESCs (EpiLCs) (**C**) from publicly available datasets. Each panel highlights a selected gene category: TRC resolution factors (blue), R-loop regulators (purple), DNA damage response factors (red), and epigenetic regulators (green). Orange dots mark significant DEGs (adjusted P < 0.05), grey dots mark non-significantly changed genes, and the colored dots highlight members of the indicated functional categories. Numbers of significantly upregulated and downregulated genes in mESCs from each selected gene category are indicated at the top right of each panel.

In contrast, comparison of mESCs and 2CLCs (**Supplementary Table 6**) revealed no marked enrichment of differentially expressed genes within any of these functional categories, with relatively balanced numbers of up- and down-regulated genes (TRC: 23 up, 38 down; R-loop: 99 up, 95 down; DDR: 56 up, 45 down; chromatin regulators: 177 up, 169 down) (**Fig. 4B**). Similarly, differential gene expression analysis of naive mESCs and primed EpiLCs (**Supplementary Table 7**) did not show a biased expression of TRC resolution genes (23 up, 13 down) R-loop (50 up, 50 down), DDR (34 up, 41 down), or chromatin regulators (158 up, 138 down) (**Fig. 4C**) (**Supplementary Table 7**), suggesting that the high expression level of these “conflict resolution” pathways is maintained during the transition from naive to primed pluripotency or in totipotent-like 2CLCs.

These data suggest that mESCs are transcriptionally programmed to manage high genomic traffic. By constitutively expressing a robust battery of conflict-resolution factors at higher level, pluripotent cells can tolerate high levels of T-R overlap without incurring catastrophic genome instability. This established "resilient” replication mode raises the question of which specific factors are most critical for its maintenance.

### A functional screen identifies AQR as a key regulator of pluripotent identity maintenance

The inherent lack of T-R coordination in mESCs, coupled with their high expression of R-loop and TRC resolution factors (**Fig. 2, 4**), suggests that pluripotency relies on a “buffered” state of high genomic traffic. We hypothesized that this resilience must be maintained by specific factors, the loss of which could tip this balance toward genomic instability and subsequent erosion of cell identity. To identify these “gatekeepers”, we performed a targeted siRNA screen focusing on 32 candidates involved in R-loop processing, TRC-related chromatin remodelling, TRC prevention and resolution, RNAPII removal at TRC sites and replication restart (**Fig. S4A**). As a sensitive readout for the loss of pluripotent identity, we quantified the spontaneous transition of mESCs to 2CLCs by flow cytometry using the 2C::tbGFP reporter cells. As a positive control, we treated mESCs with 10μM retinoic acid (RA), which was previously shown to induce the expression of *Dux* and *Duxbl1* transcription factor and robustly increase 2CLCs ^25,26^ (**Fig. 5A**).

**Figure 5.**
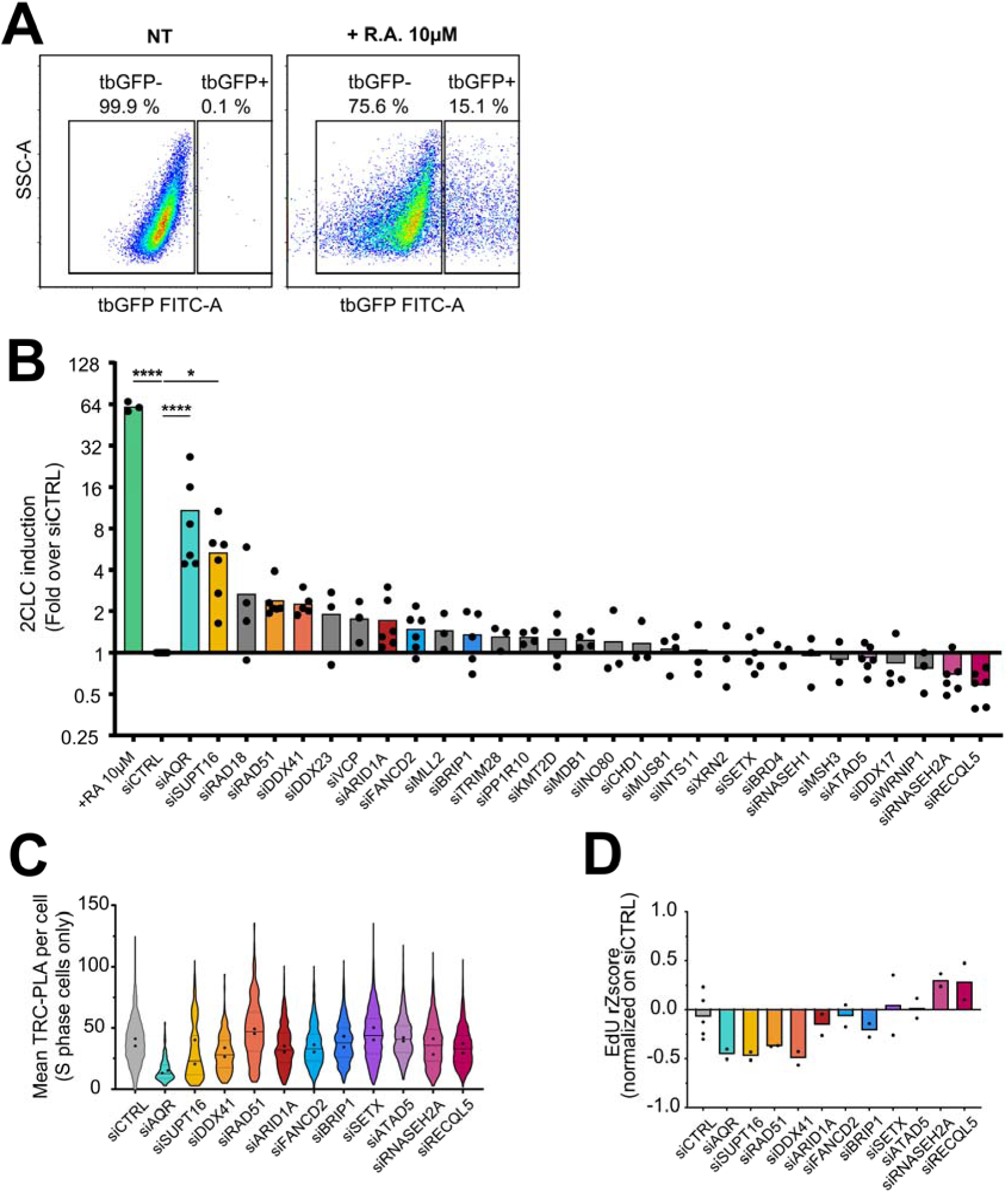
siRNA knockdown screen identifies siAQR as a regulator of 2CLC induction and TRC resolution. **(A)** FACS profiles (SSC-A vs tbGFP-A) of mESCs to identify and quantify the amount of 2CLCs (tbGFP+ cells) in mESC cultures. Non-treated (NT, left) show the typical mESC culture profile; mESCs treated with R.A. (right) induce 2CLC emergence. (**B**) FACS quantification of 2CLCs in MERVL::tbGFP-PEST reporter mESCs following siRNA KD. 2CLCs were identified by turboGFP expression. Data is presented as fold induction relative to siCTRL. Treatment with 10 µM retinoic acid (RA) for 48 h served as a positive control for 2CLC induction (N=3-6). Statistical significance was determined by paired one-way ANOVA followed by Bonferroni’s multiple comparisons test against siCTRL. (**C**) Quantification of TRC burden as the number of TRC-PLA foci per nucleus in EdU-positive cells following siRNA KD. Each dot represents one biological replicate (N = 2), calculated as the mean of ≥3 technical replicates (50–3,000 analyzed cells per technical replicate). (**D**) Quantification of EdU incorporation in EdU-positive cells following siRNA KD. EdU levels were normalized using robust Z-score transformation. Each dot represents the mean value of at least 3 technical replicates (N =2), calculated as the mean of ≥3 technical replicates (50–3,000 analyzed cells per technical replicate).

The majority of siRNAs led to a robust knockdown (KD) of their targets to 20-50% of the original mRNA levels as confirmed by RT-qPCR (**Fig. S4B**). Individual KD of most of the analyzed TRC factors had no or limited effect on reprogramming to 2CLCs (**Fig. 5B**). We postulate that one reason for this is the robust expression of compensatory R-loop/TRC resolution pathways that we identified in mESCs (**Fig. 4)**. However, depletion of a select few candidates significantly induced 2CLC emergence. Among these, the DEAD-box helicase AQR (*Aqr*) emerged as the most critical hit, with its depletion resulting in a ∼16-fold increase in the 2CLC population (**Fig. 5B**). While loss of the FACT complex subunit *Supt16* also prompted 2CLC emergence (**Fig. 5B**), AQR stood out for its unique potency and its independence from established DNA repair pathways.

We next investigated whether this identity loss was a direct consequence of elevated TRC levels. Surprisingly, AQR-depleted cells exhibited a reduction in TRC-PLA foci and a concomitant decline in global replication activity, as measured by EdU incorporation (**Fig. 5C-D**). While the decrease in TRC-PLA may result from reduced replicative activity rather than a specific decline in TRC frequency, these data suggest that the 2CLC induction might be a passive response to slowed replication forks, a phenomenon previously linked to totipotent-like reprogramming^2,27^. However, we found that further slowing of the fork with hydroxyurea (HU) in AQR-depleted cells exacerbated the emergence of 2CLCs (**Fig. S4C**). This supports a model in which AQR limits 2CLC emergence through a mechanism that cannot be explained solely by replication slowing and establish AQR as an essential safeguard of the pluripotent cell state.

### AQR maintains T-R coordination to sustain high replication rates in mESCs

While AQR is recognized as a conserved RNA helicase and component of the intron-binding spliceosomal complex in somatic cells^28–33^, little is known about AQR’s function in the unique regulatory environment of pluripotent stem cells. Given our screen results, we hypothesized that AQR is the primary factor responsible for managing the characteristic high genomic traffic of mESCs. Indeed, AQR depletion (siAQR) (**Fig. S5A, B**) resulted in a marked reduction in the proportion of EdU-positive cells and a significant decline in DNA synthesis rates among the remaining S-phase population (**Fig. 6A–C**). This replicative failure was accompanied by the accumulation of RS markers including FANCD2^34^ and γH2AX (**Fig. 6E-G**). Notably, these markers were specifically enriched in EdU-positive cells, indicating that DNA damage in mESCs is an active consequence of attempting to replicate the genome in the absence of AQR’s protective function.

**Figure 6.**
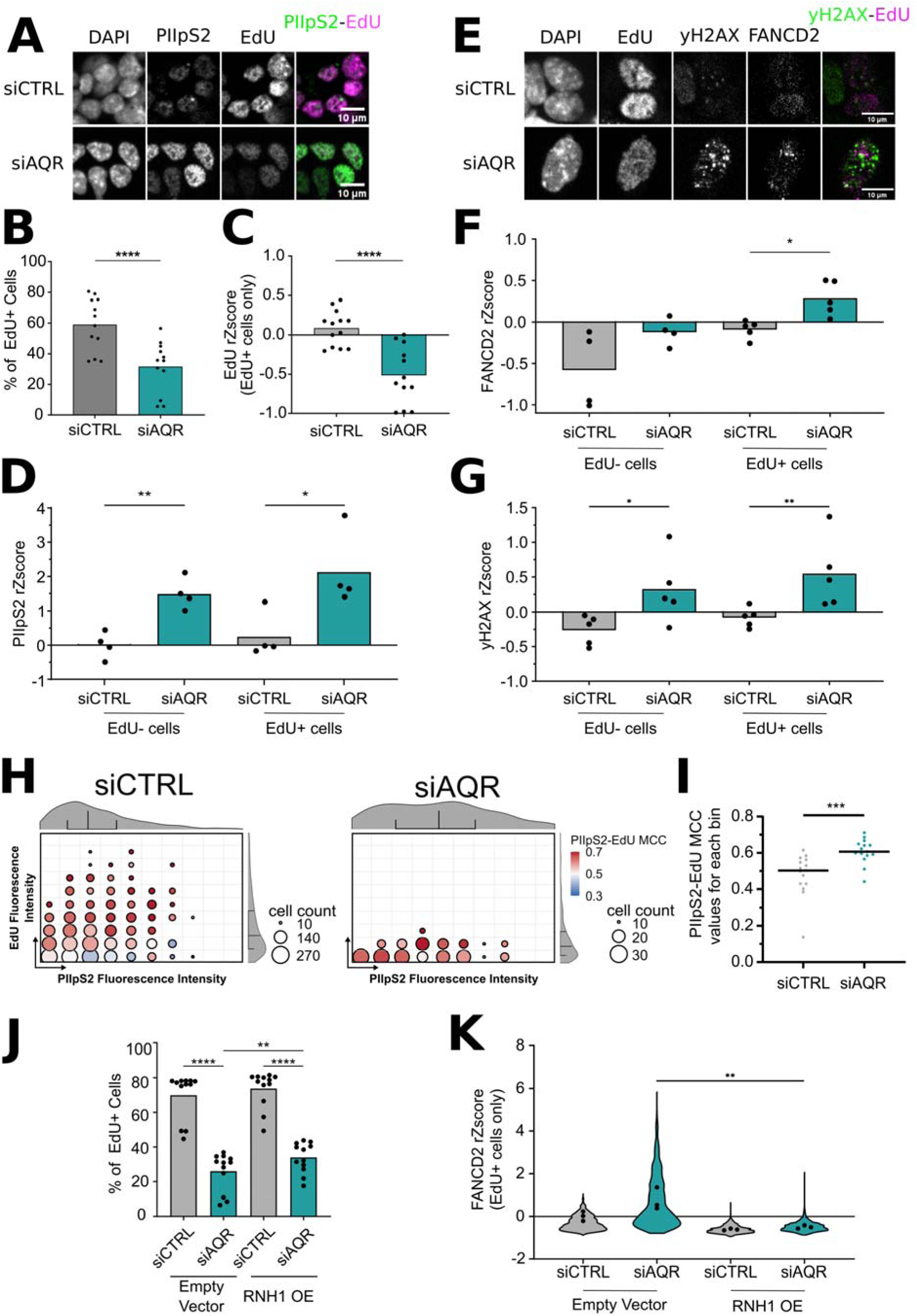
Aquarius depletion disrupts transcription–replication coordination and induces replication stress in mESCs. **(A**) Representative immunofluorescence images of mESCs transfected with siCTRL (top) or siAQR (bottom). Active transcription was visualized by chromatin-bound PIIpS2, and active DNA replication was detected by EdU incorporation. (**B**) Percentage of EdU-positive cells following siRNA transfection. Each dot represents the mean value of at least 3 technical replicates (N =12), calculated as the mean of ≥3 technical replicates (150–3,600 analyzed cells per technical replicate). Statistical significance determined by a two-tailed paired t-test. (**C**) EdU incorporation levels in EdU-positive cells normalized replicate-wise using robust Z score transformation. Each dot represents the mean value of at least 3 technical replicates (N =12), calculated as the mean of ≥3 technical replicates (150–3,600 analyzed cells per technical replicate). Statistical significance determined by a two-tailed paired t-test. (**D**) Chromatin-bound PIIpS2 signal in EdU-negative (left) and EdU-positive (right) cells across siRNA conditions. Values normalized replicate-wise by robust Z score transformation. Each dot represents the mean value of at least 3 technical replicates (N =4), calculated as the mean of ≥3 technical replicates (150–3,600 analyzed cells per technical replicate). Statistical significance was determined by paired one-way ANOVA followed by Bonferroni’s multiple comparisons test. (**E**) Representative immunofluorescence images of mESCs transfected with siCTRL (top) or siAQR (bottom), showing EdU incorporation (replication), γH2AX (DNA damage), and FANCD2 (replication stress). (**F**) Quantification of chromatin-bound FANCD2 in EdU-negative (left) and EdU-positive (right) cells across siRNA conditions. Normalized replicate-wise by robust Z score transformation. Each dot represents the mean value of at least 3 technical replicates (N =4-5), calculated as the mean of ≥3 technical replicates (150–3,600 analyzed cells per technical replicate). Statistical significance was determined by paired one-way ANOVA followed by Bonferroni’s multiple comparisons test. (**G**) Quantification of chromatin-bound γH2AX in EdU-negative (left) and EdU-positive (right) cells across siRNA conditions. Normalized replicate-wise by robust Z score transformation. Each dot represents the mean value of at least 3 technical replicates (N =5), calculated as the mean of ≥3 technical replicates (150–3,600 analyzed cells per technical replicate). Statistical significance was determined by paired one-way ANOVA followed by Bonferroni’s multiple comparisons test. (**H**) PIIpS2–EdU MCC bubble plots showing the effect of siAQR on transcription–replication coordination (N = 2; 50–1,000 analyzed cells per replicate). Cells are binned by PIIpS2 intensity (X-axis) and EdU intensity (Y-axis) (see “Image analysis” in the methods for further information on the data processing); bubble size indicates cell count per bin, and bubble color encodes median MCC values. (**I**) Bin-wise comparison of MCC values from (**H**) between siCTRL and siAQR treated cells. MCC values from all the bins present in the siAQR bubble plot and the corresponding bins in the siCTRL bubble plot bins defined in panel (**H**) were extracted. Statistical significance of the difference in MCC between cell types was assessed using a two-tailed paired t-test across all included bins. Each dot represents the MCC value of a single bin. (**J**) Partial rescue by hRNASEH1 OE on the fraction of EdU-positive cells following siAQR transfection. Each dot represents the mean value of a technical replicate of 2 biological replicates. Dot shape represents biological replicates (483–2,942 analyzed cells per technical replicate). Statistical significance was determined by paired two-way ANOVA followed by Bonferroni’s multiple comparisons test. (**K**) Rescue by hRNASEH1 OE on replication stress, quantified by the level of chromatin-bound FANCD2 in EdU-positive cells, normalized replicate-wise by robust Z score transformation. Violin plots represent the distribution of cells across replicates, and each dot represents a technical replicate (N =1), (160–2,942 analyzed cells per technical replicate). Statistical significance was determined by paired two-way ANOVA followed by Bonferroni’s multiple comparisons test.

Interestingly, AQR-depleted cells showed a striking accumulation of chromatin-bound PIIpS2 specifically during S-phase (**Fig. 6D**). This suggests that AQR is required to prevent transcriptional stalling that otherwise acts as a physical barrier to the replication machinery. To quantify this failure in spatial coordination, we utilized our PIIpS2-EdU MCC analysis (**Fig. 6H, Fig. S5C**). As expected from the quantification of EdU levels, PIIpS2-EdU MCC quantification revealed that AQR depletion results in a strong decrease of cells with high EdU incorporation levels (**Fig. 6C, H**, changes along the Y axis). Strikingly, we found that AQR loss profoundly impaired T-R coordination, shifting a large proportion of cells towards high MCC values (>0.7) (**Fig. 6H, I, Fig. S5C**), even in cells with low EdU levels, normally characterized by low MCC values (**Fig. 6H, I**). These results clearly demonstrate that AQR loss compromises T-R coordination in mESCs, leading to reduced replication activity and the onset of RS-associated DNA damage.

### Dual roles of AQR in splicing and R-loop homeostasis safeguards pluripotency

Given AQR’s dual roles as a spliceosomal component and an RNA:DNA helicase, we next sought to determine whether the erosion of pluripotent identity (2CLC transition) was a secondary consequence of global splicing failure, or a direct result of R-loop induced replication stress. First, we performed epistasis analysis by combining AQR knockdown with the splicing inhibitor Pladienolide B (PlaB). While PlaB treatment alone induced a 5-9-fold increase in 2CLCs, its combination with AQR depletion yielded a potent synergistic effect (25-28-fold increase) that exceeded either perturbation alone (**Fig. S5D**). Thus, while PlaB is known to induce R-loop accumulation at splice sites, these results indicate that AQR’s spliceosomal function alone cannot account for its role in suppressing 2CLC emergence.

As PlaB-mediated splicing inhibition has been shown to also induce R-loop accumulation at splicing sites^35^, we further refined our analysis by co-depleting AQR with splicing factors that act concurrently (*Dhx16*), or downstream (*Prpf8*) of AQR recruitment. Co-depletion with *Dhx16* showed additive effects, suggesting that the function of AQR in the suppression of 2CLC emergence is not solely mediated through its spliceosomal activity. In contrast, co-depletion with *Prpf8* precisely phenocopied the AQR single knockdown, suggesting that other splicing complexes downstream of AQR function can also impact 2CLC emergence (**Fig. S5E**). These results suggest that splicing failure and R-loop-associated replication stress each independently promote 2CLC entry, both of which AQR is involved in to maintain genomic stability and cell fate. To further support this, we examined the unspliced R-loop prone *Airn* locus to decouple AQR’s role in R-loop homeostasis from its function in mRNA splicing. In agreement, AQR depletion triggers a significant accumulation of RNaseH1-sensitive R-loops at the Transcription Start Site (TSS) (**Fig. S5F**), providing direct evidence that AQR suppresses R-loop formation through a distinct helicase activity independent of its spliceosomal context.

We then asked whether the replication stress and cell cycle defects associated with pluripotency exit upon AQR depletion were R-loop dependent. To address this, we overexpressed human RNAseH1 (hRNASEH1) to specifically degrade RNA:DNA hybrids in siAQR-treated cells (**Fig. S5G**). Exogenous hRNASEH1 significantly reduced chromatin-bound FANCD2 levels, a hallmark of R-loop induced replication stress, and partially mitigated the proliferation defects induced by AQR knockdown (**Fig. 6J-K**). However, hRNASEH1 expression failed to restore global replicative activity within S-phase cells (**Fig. S5H**), which likely reflects a combination of persistent splicing-dependent defects and secondary effects of hRNASEH1 overexpression, which itself elevated γH2AX levels in mESCs (**Fig. S6I**). Despite this, the specific rescue of FANCD2 (**Fig. 6K**) demonstrates that R-loop accumulation is a major driver of replication fork stalling in AQR-depleted mESCs. Collectively, these findings demonstrate that AQR’s essential role in pluripotency extends beyond splicing to include the resolution of R-loop-dependent TRCs that would otherwise threaten the pluripotent genome.

### AQR depletion specifically impairs R-loop and TRC resolution pathways underlying the “resilient” replication mode

To characterize the transcriptomic changes associated with AQR depletion, we performed differential gene expression analysis in control and AQR-depleted mESCs (**Supplementary Table 8**). Loss of AQR triggered profound transcriptional reprogramming, involving 2,641 up-regulated and 3,023 down-regulated genes. Of these, 1,401 up-regulated and 292 down-regulated genes exhibited a fold change greater than 2. (**Fig. 7A**). In parallel, analysis of alternative splicing revealed widespread splicing defects upon AQR depletion, with a marked accumulation of retained introns and skipped exons, consistent with AQR’s role in intron recognition and removal (**Fig. 7B**). Gene Ontology (GO) enrichment analysis of differentially expressed genes revealed that down-regulated genes were strongly enriched for terms associated with cellular metabolism, proliferation, and DNA replication (**Fig. 7C, left**), whereas up-regulated genes were enriched for processes related to transcriptional regulation and crucially, non-pluripotent developmental programs including cell fate commitment and lineage-specific morphogenesis such as cardiac atrium morphogenesis and skeletal muscle development (**Fig. 7C, right**). Together, these data indicate that AQR depletion induces transcriptional changes that extend beyond replication stress and DNA damage, and are suggestive of a destabilization of the naive pluripotency transcriptional program and a concomitant upregulation of lineage-associated gene expression.

**Figure 7.**
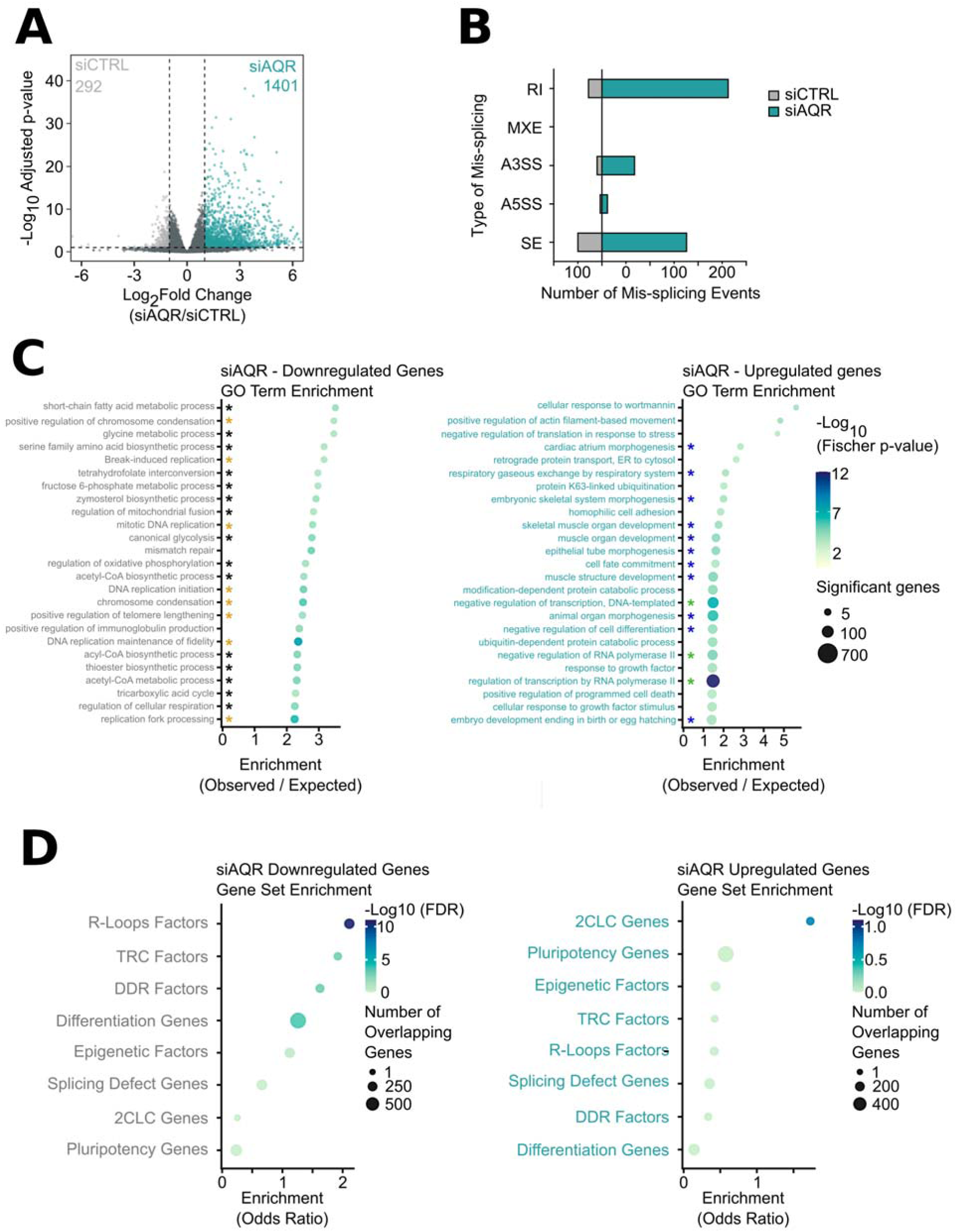
AQR depletion dysregulates the transcription program of mESCs. (**A**) Volcano plot of differential gene expression following siAQR knockdown. Log2 fold change (siAQR / siCTRL) is shown on the x-axis and –log10 p-value on the y-axis. Genes with log2 fold change ≥ 1 and p < 0.05 are colored aqua (1401 upregulated genes) or grey (292 downregulated genes), with dashed lines indicating fold change thresholds (±1) and significance (p = 0.05). Data are pooled three independent RNA-seq replicates per condition. (B) Differential splicing events between siCTRL and siAQR-treated cells. Alternative splicing effects were identified from bulk RNA-seq data using rMATS (skipped exons (SE), alternative 5′ splice sites (A5SS), alternative 3′ splice sites (A3SS), retained introns (RI), and mutually exclusive exons (MXE)). Significant splicing alterations between replicates were defined using an FDR threshold of <0.1. Numbers shown represent the total number of significant splicing defects in each category amongst the biological replicates (N=3). (**C**) GO biological process enrichment of differentially expressed genes upon AQR depletion. Dot plots show the top 25 enriched terms for downregulated (left) and upregulated (right) genes. Dot size indicates the number of significant genes per term, and color reflects statistical significance (−log₁₀ Fisher p-value). Enrichment is expressed as the observed/expected gene ratio. Terms were selected based on stringent statistical criteria (Fisher.classic < 1×10⁻³ and Fisher.elim < 0.05). Terms attributed to metabolism, replication, transcription, and to development and lineage specific aspects are highlighted with a black, yellow, green or a blue star, respectively. (**D**) Targeted Gene Set Overlap analysis of differentially expressed genes (DEGs). Dot plots show the enrichment of the differentially expressed genes (up- and downregulated genes upon AQR depletion) against multiple curated gene lists, with downregulated genes on the left and upregulated genes on the right. Dot size represents the number of overlapping genes between the query and target lists, color indicates statistical significance (−log₁₀ FDR), and the x-axis shows the odds ratio as a measure of enrichment. The curated gene lists tested are the lists of TRC resolution factors, R-loop regulators, DDR factors, and epigenetic regulators used in **Fig 2D**, **S3B**, and **S4C**. Additionally, DEGs were tested against the list of genes showing splicing defects upon AQR depletion and against genes linked to pluripotency maintenance, differentiation, and transition to 2CLCs.

We next asked whether this identity shift was coupled with a failure of the genome maintenance systems that characterize naive mESCs. Using targeted Gene Set Overlap analysis (GSO), we confirmed that genes upregulated upon AQR loss were significantly enriched for the mESC-to-2CLC transition^21^ (**Fig. 7D, right**). Strikingly, the most pronounced transcriptomic signature of AQR depletion was the coordinated downregulation of the “resilience” program. We observed a significant and systemic reduction in the expression of genes involved in TRC resolution, R-loop processing, and DNA repair pathways – the exact categories we previously found to be constitutively upregulated in naive mESCs (**Fig. 7D, left**). Importantly, we detected no enrichment of mis-spliced transcripts among these differentially expressed resolution factors (**Fig. 7D**), indicating that their downregulation is a regulated systemic response rather than a direct consequence of splicing failure.

Together, these results demonstrate that AQR depletion does not simply induce isolated splicing defects or gene dysregulation at individual loci. Rather, loss of AQR leads to widespread transcriptional changes that attenuates multiple genome-protective pathways, shifting cells from a resilient state toward one with diminished capacity for TRC and R-loop resolution. This systemic reduction in genome maintenance potential is likely the cause for increased replication stress and genomic instability of AQR-depleted mESCs.

### AQR prevents destabilization of the pluripotent state of naive and primed mESCs

We next wished to investigate the extent to which perturbing genomic resilience via AQR KD influences the pluripotent identity. We used siRNA-mediated depletion of AQR in naive mESCs (Lif/2i) and monitored the expression of 2CLC, naive and primed pluripotency and lineage-specific differentiation markers (ectoderm, mesoderm, endoderm) by RT-qPCR (**Fig. 8A**). AQR loss induced a heterogenous expression profile, characterized by the simultaneous activation of 2CLC signatures and a divergent response among core pluripotency factors: while Oct3/4 levels remained stable, we observed a significant decline in Sox2. Notably, we observed an aberrant activation of lineage-primed markers, most notably for the endoderm (*Gata4*, *Gata6*) (**Fig. 8A**). This expression profile suggests that AQR loss destabilizes the pluripotent state and creates permissive conditions for transition towards early differentiation programs^36^.

**Figure 8.**
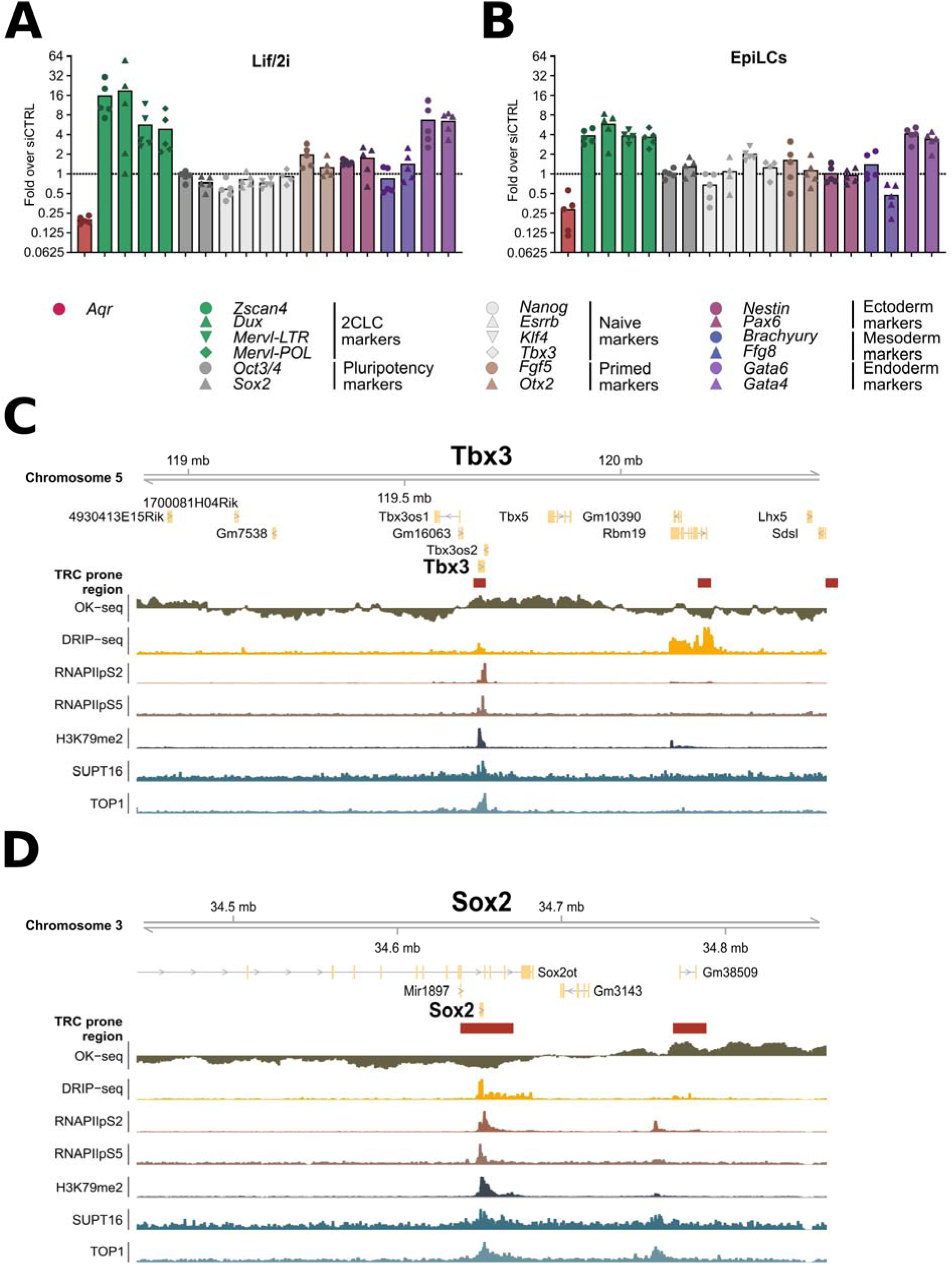
AQR depletion destabilizes cell fate transcriptional programs. **(A)** RT-qPCR analysis of cell fate–related genes upon siAQR KD in mESCs maintained in naive conditions. Cells were transfected with siCTRL or siAQR and cultured for 48 h in Lif/2i before RNA extraction. Relative expression was calculated by the ΔΔCt method, normalized to *Actinβ*, *Gapdh*, and *Rpl13a*, with siCTRL as the reference (N=5). (**B**) RT-qPCR analysis of cell fate–related genes upon siAQR KD during priming of mESCs to the EpiLC state. Transfection with siCTRL or siAQR was performed concomitantly with the switch to priming media. Expression values were calculated as in (**A**). (N=5) (**C-D**) Genomic snapshots of pluripotency genes *Tbx3* (**C**) and *Sox2* (**D**) showing their TRC related genomic context, with *Tbx3* showing a propensity for co-directional TRCs (**C**), and *Sox2* showing a propensity for head-on TRCs (**D**). Tracks shown from top to bottom include: predicted TRC-prone regions defined using a TRC prone region prediction framework developed by Aguilera and colleagues ^38^ OK-seq; DRIP-seq; RNAPII-pS2 ChIP-seq; RNAPII-pS5 ChIP-seq; H3K79me2 ChIP-seq; SUPT16 ChIP-seq; and TOP1 ChIP-seq.

The impact of AQR depletion was even more pronounced when performed during directed cell state transitions. When mESCs were primed towards EpiLCs, AQR deficiency led to a transcription profile characterized by a simultaneous up-regulation of competing markers (**Fig. 8B**). For example, *Sox2* and *Klf4* (naive) increased alongside 2CLC and endoderm markers, while *Nanog* significantly declined (**Fig. 8B**). In addition, the two mesoderm markers showed a heterogenous response with a mild induction of *Brachyury* but reduced *Fgf8* expression. This transcriptional heterogeneity suggests that AQR activity is required to prevent the co-expression of antagonistic transcription programs. Interestingly, although the chromatin remodeler FACT (*Supt16*) was a top hit in our initial screen, its co-depletion with AQR yielded only minor additive effects on marker gene expression, establishing AQR as the primary driver of this identity destabilization (**Fig. S6A-B**).

Because AQR triggers RS and activates the DNA damage response (**Fig. 6**), we next asked whether the cellular response to this stress is what drives the identity erosion. We focused on the ATR kinase, a master sensor of RS^37^. Strikingly, ATR inhibition (ATRi) exacerbated transcriptional heterogeneity in AQR-depleted cells and triggered the erratic expression of priming and lineage-specific markers. Indeed, ATR-inhibited cells selectively induced *Fgf5*, *Pax6*, and *Fgf8*, while failing to activate their counterparts *Otx2*, *Nestin*, and *Brachyury* **(Fig. S6C**). This implicates the ATR checkpoint in identity maintenance, especially when the “resilient” replication mode is compromised by AQR depletion.

Finally, we asked whether resolving R-loops can restore identity. Exogenous overexpression of hRNASEH1 in AQR-depleted cells effectively suppressed the aberrant upregulation of the key 2CLC marker *Dux*, the primed marker *Otx2* and the Endoderm lineage-specific differentiation markers *Gata6* (**Fig. S6D**). While hRNASEH1 did not fully restore the core pluripotency program, it specifically reversed several of the identity transitions induced by AQR loss, directly linking R-loop accumulation to the observed destabilized pluripotency program.

Collectively, our results demonstrate that AQR loss destabilizes mESC identity in an R-loop–dependent and partially ATR checkpoint–dependent manner, promoting 2CLC emergence and a heterogeneous erosion of pluripotency.

### Core pluripotency genes reside in conflict-prone genomic hotspots sensitive to AQR loss

We next asked whether the selective sensitivity of these core pluripotency genes is a consequence of their unique genomic context. To test this, we utilized a predictive framework integrating R-loop mapping (DRIP-seq) with replication fork directionality profiles (OK-seq)^38^ to identify regions of high TRC probability. Strikingly, several central regulators of the pluripotency network mapped precisely to these predicted TRC hotspots. Specifically, we identified *Tbx3* and *Esrrb* within R-loop–prone co-directional TRC regions (**Fig. 8C, Fig. S6E**), while *Sox2* and *Klf4,* the very factors that showed erratic expression upon AQR loss, lie within predicted head-on conflict zones (**Fig. 8D, Fig. S6F**). This localization in conflict prone regions was further reinforced by a convergence of conflict markers: they are significantly enriched for H3K79me2, a histone modification associated with TRC-prone chromatin^13^, and show high occupancy of the chromatin remodeler SUPT16 (FACT) and the topoisomerase TOP1 (**Fig. 8C, D, Fig. S6E, F**).

This unique genomic signature, including high R-loop propensity, intense chromatin remodeling and torsional stress and TRC-associated chromatin modifications robustly identify these loci as conflict-sensitive genomic regions. Collectively, these observations suggest that AQR depletion exposes critical pluripotency loci to TRCs, generating localized transcriptional instability that drives the overall observed decline in the pluripotent state.

### Single-cell profiling reveals that AQR loss increases the transcriptional plasticity of mESCs

To characterize the impact of AQR depletion on pluripotency maintenance at high resolution, we performed single cell RNA sequencing in mESCs maintained in their naive state (Lif/2i) or under priming conditions (EpiLCs) in siCTRL and siAQR treated cells. We obtained transcriptome profiles from 91,086 high-quality cells (34,783 Lif/2i siCTRL; 26,433 Lif/2i siAQR; 7,983 EpiLCs siCTRL; and 21,887 EpiLCs siAQR cells), detecting an average of 11,266 unique transcripts and 3,306 genes per cell. Unsupervised clustering identified 17 clusters of cell types, with clear separation of clusters between siCTRL and siAQR and Lif/2i (naive) versus EpiLCs (primed) cell states (**Fig. 9A**, **Fig. S7A, B, C**). We next annotated clusters based on established lineage and pluripotency markers (**Fig. 9B, C, Fig. S7D**), which allowed us to define the naive and primed pluripotent mESC populations, as well as an epithelial-like cluster which was equally represented across all conditions, suggesting it reflects a stable cell type in mESC cultures rather than an AQR-dependent transition (**Fig. 9C, D Fig. S7D**). Strikingly, AQR loss prompted the emergence of several new clusters under both Lif/2i and EpiLC conditions that were highly enriched in siAQR condition. These include *Zscan4*⁺ cells, corresponding to a transitional population between mESCs and 2CLCs; endoderm-like cells expressing *Gata4*, *Gata6*, *Sox17*, *Sox7*, and *Dnmt3a/b*; and formative-like cells, a state intermediate between naive and primed pluripotency characterized by mixed expression of markers from both states (**Fig. 9C, D, Fig. S7D**).

**Figure 9.**
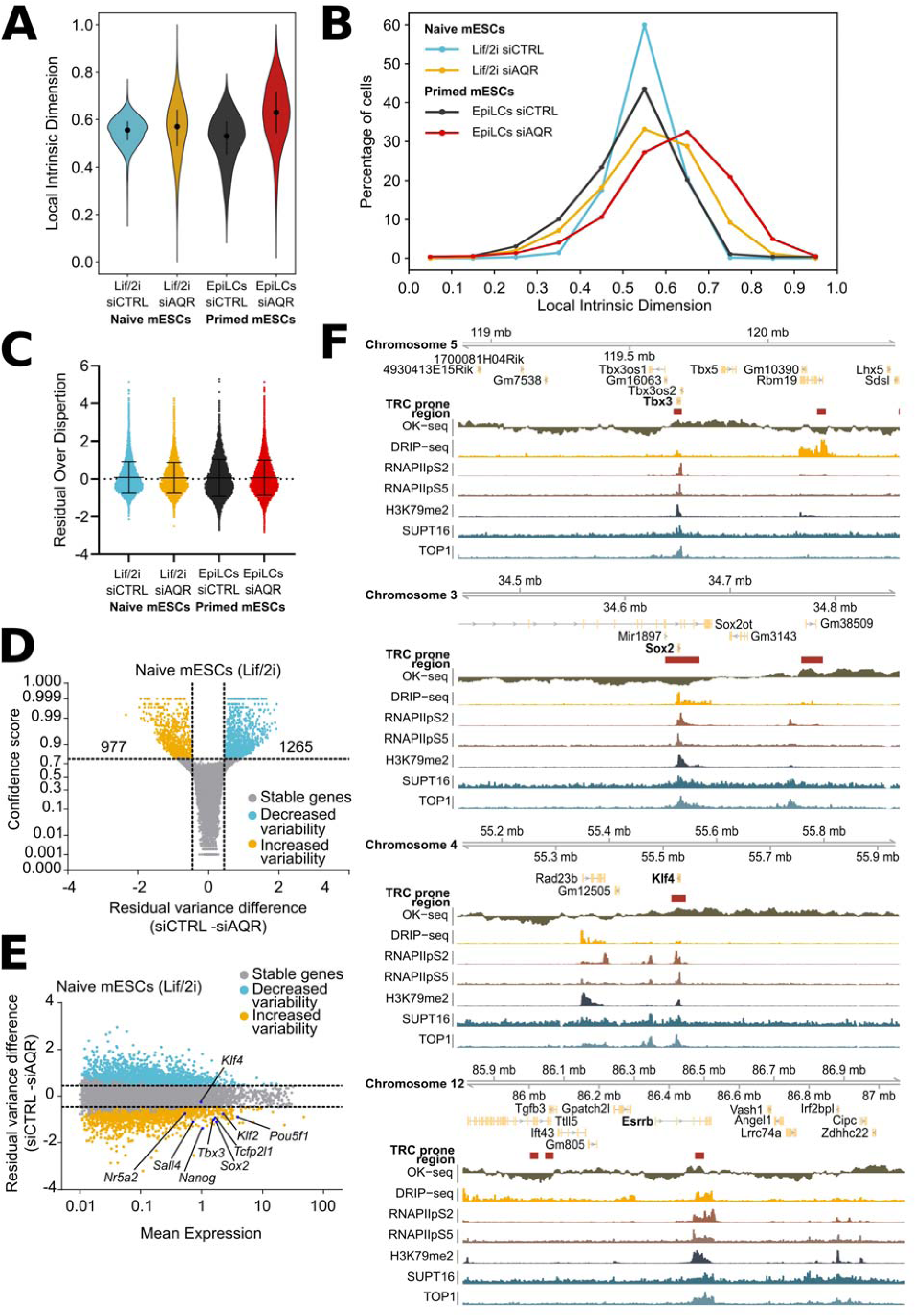
AQR depletion increases transcriptional plasticity and alters transcriptional variance in pluripotent mESCs. (A) UMAP visualization of single-cell transcriptomes showing the distribution of siCTRL and siAQR cells across naive (Lif/2i) and primed (EpiLC) conditions. (**B**) UMAP projection showing distinct cell identity clusters identified by Louvain clustering and annotated based on established pluripotency and lineage markers. (**C**) Characterization of the contribution of the different annotated cell lineage and pluripotency clusters to the different sample populations. (**D**) Dot plot displaying expression of different identity associated markers across the different annotated cell lineage and pluripotency clusters. Dot color reflects average expression levels, while dot size indicates the proportion of cells expressing each marker. (**E**) Violin plots with inter-quartile range showing the distribution of single-cell local intrinsic dimensionality, as measured by the IDEAS framework, in siCTRL and siAQR conditions within the naive mESC cluster (Lif/2i) and the primed mESC cluster (EpiLCs). (**F**) Frequency distributions of intrinsic dimensionality for the same cell populations shown in (**E**), highlighting the broadening of transcriptional state space upon AQR depletion. (**G**) Global transcriptional variance quantified using BASiCS, represented as the distribution of gene residual over-dispersion values in siCTRL and siAQR samples from naive (Lif/2i) and primed (EpiLC) mESCs. Error bars depict inter-quartile range. (**H**) Differential transcriptional variance in naive mESCs, plotted as confidence score versus residual variance difference (siAQR - siCTRL). Dashed lines indicate thresholds for statistical significance. Numbers denote the total genes exhibiting significantly decreased variability (blue) or increased variability (yellow) upon AQR depletion. (**I**) Differential transcriptional variance in naive mESCs represented as residual variance difference (siAQR - siCTRL) versus mean gene expression. Highlighted genes in blue correspond to key pluripotency regulators.

The appearance of these clusters suggests that AQR is required to maintain the pluripotent state; without it, the barriers preventing spontaneous drift into alternative lineages begin to collapse. To quantify this, we employed intrinsic dimensionality (ID) analysis via the IDEAS framework^39^, which estimates cellular potency by determining the minimum number of independent transcriptional programs required to define the variability within a cell population. While low ID values correspond to transcriptionally constrained, well-defined identities, higher ID values reflect a broadened transcriptional landscape and increased plasticity for lineage transitions.

Application of IDEAS revealed that AQR depletion significantly increases the ID of both naive mESCs (Lif/2i) and primed mESCs (EpiLCs), indicating a transition toward a less constrained, more plastic transcriptional state (**Fig. 9E**). Strikingly, we observed a simultaneous broadening of the single-cell ID distribution (**Fig. 9F**). This suggests that AQR loss induces increased transcriptional variance, where specific subpopulations exhibit enhanced potency, while others display higher constraints and lower potency. Together, these findings position AQR as a critical stabilizer of pluripotent identity, limiting the accessible transcriptional state space.

Interestingly, this increased plasticity was not driven by a single "differentiation" pathway. Differential gene expression (DEG) analysis revealed a broad, non-specific misregulation of transcripts (**Fig. S7E, F**), suggesting a global loss of regulatory constraint. Notably, AQR-depleted naive cells upregulate *Tcea3*, one of the three TFIIS isoforms involved in transcription restart and previously shown to affect the differentiation potential of mESCs^40^. In EpiLCs, AQR KD simultaneously induced primed-state markers (e.g. *Tdgf1*, *Fgf5*) and various neuronal/lineage-biased genes (*Ap3b2, Dpp6, Kctd16*), while repressing other lineage regulators such as *Gria3*, *Ptprt* and *Tbx4*. Together, the observed gene expression changes argue that loss of AQR affects expression of a large number of genes without clear functional specificity.

Transcriptional variance or noise (defined as stochastic fluctuations in gene expression) has been recently described to promote cell fate transitions^41^. To determine whether this may also contribute to the observed cell fate transitions upon AQR loss, we quantified the residual over dispersion as a metric of transcriptional variance using Bayesian Analysis of Single-Cell Sequencing (BASiCS)^42^ in siCTRL and siAQR treated cells **(Fig. 9G)**. To account for potential variations arising from shifts in cell-state composition, we restricted our analysis to the naive mESC cluster in Lif/2i and to the primed cluster in EpiLCs (**Supplementary Table 9, 10**).

Under these refined parameters, rather than a global increase in transcriptional variance, AQR depletion induced a redistribution of variability, with comparable numbers of genes gaining and losing transcriptional stability in both naive (977 more variable / 1265 less variable genes) and primed states (1733 more variable / 1617 less variable genes) (**Fig. 9H, Fig. S9A**). However, GO term analysis revealed that this redistribution was non-random in relation to the function of the affected genes: In naive mESCs, genes gaining variability were significantly enriched for genomic imprinting and developmental processes (**Fig. S8A**), whereas genes involved in metabolic pathways remained stable (**Fig. S8B**). Conversely, in primed mESCs, increased variance was concentrated in genes regulating transcription, metabolism, and cell morphology (**Fig. S9C, D**). Targeted gene-set analyses further revealed overlap between genes affected in naive and primed cells, particularly increased variability in 2CLC-related genes across both states (**Fig. S8C, D, Fig. S9E, F**). Notably, in primed mESCs, genes with increased variance significantly overlapped with those upregulated upon AQR knockdown (**Fig. S9E, F**). In naive mESCs, genes gaining variability were enriched for R-loop resolution factors (**Fig. S8C, D**), reinforcing the model that AQR activity is essential for maintaining R-loop–resilient transcriptional states.

To further investigate the link between pluripotency maintenance and transcriptional variance, we examined the behavior of an extended subset of genes associated with the core pluripotency network, several of which we found to be positioned in TRC-prone regions (**Fig. 8C**).

Intriguingly, most investigated factors, including *Pouf51* (*Oct3/4*), *Sox2*, *Tbx3*, *Nanog*, *Klf2*, *Tcfp2l1*, *Sall4*, and *Nr5a2*, showed increased variability upon AQR depletion in naive, but not in primed mESCs (**Fig.9I, Fig. S9B**). This amplification of transcriptional noise at TRC-prone identity genes directly links AQR-mediated conflict resolution to the stabilization of the pluripotency gene expression program. Together, these data demonstrate that loss of AQR-mediated protection at these high-traffic loci is sufficient to expand the accessible transcriptional landscape and facilitate stochastic transitions toward alternative cell fates.

## Discussion

The maintenance of pluripotency requires a delicate balance between rapid self-renewal, a dynamic transcription program that maximizes proliferative capacity and the preservation of genomic integrity. Traditionally, the field has viewed transcription-replication coordination as an important requirement for survival, where cells must strictly segregate these machineries to avoid genotoxic collisions. Our results challenge this paradigm as we show that naive pluripotent stem cells operate under a unique “resilient” replication mode, characterized by high genomic traffic and a constitutively upregulated "resolution signature" of R-loop and TRC factors to prevent catastrophic instability (**Fig. 1, 2, 4**). We propose that this ability to proliferate with reduced coordination may represent a developmental adaptation, enabling stem cells to accommodate the extensive transcriptional plasticity required for differentiation and reprogramming.

Using single-cell microscopy, we introduce T–R coordination as a quantitative and dynamic property that differs markedly between pluripotent, somatic, and totipotent-like states and adapts rapidly to transcriptional reprogramming (**Fig. 2**). This suggests that the degree of T-R coordination, alongside transcriptional output, replication timing, and chromatin organization, represents an underappreciated layer that could be used as a “fingerprint” to detect different cell identities, reflecting the balance between genome duplication and transcriptional regulation across developmental states. Therefore, we suggest that T–R coordination is not merely a by-product of transcriptional activity, but rather an actively tuned parameter of genome regulation that changes with cell identity (**Fig. 3**). This plasticity likely facilitates stable lineage transitions by mitigating the inherent risk of TRCs during periods of rapid transcriptional rewiring.

Our functional screen identified the RNA:DNA helicase AQR as the primary safeguard of this resilient state (**Fig. 5**). Depletion of AQR leads to a 16-fold increase in 2CLC emergence (**Fig. 5B**) and a systemic collapse of replication activity (**Fig. 6A-C**) and T-R coordination (**Fig. 6H**). AQR serves as the catalytic core of the pentameric Intron-Binding Spliceosomal Complex (IBC), operating in coordination with XAB2, ISY1, ZNF830, and PPIE to modulate the RNA:DNA interface during splicing. Notably, while the majority of spliceosomal helicases belong to the SF2 family, AQR is a member of the highly processive SF1-family^43^, a structural distinction consistent with a specialized or non-canonical function beyond traditional mRNA processing. This specialized role is further supported by the precise anchoring of the IBC at a fixed intronic position, 33-40 nucleotides upstream of the branch point^44^. This specific localization aligns with recent high-throughput mapping studies demonstrating that genomic RNA:DNA hybrids are preferentially enriched within the intronic regions of actively transcribed genes. Such strategic positioning provides the IBC with an ideal spatial orientation to sense and unwind intronic R-loops, potentially independent of its core spliceosomal duties. This model is further reinforced by the presence of XAB2 within the complex, a factor with established roles in transcription-coupled repair and R-loop metabolism^45,46^. Together, these features suggest that the IBC possesses a dual capacity: facilitating efficient splicing while simultaneously safeguarding the DNA template by purging genotoxic RNA:DNA hybrids from intronic loci. This non-canonical safeguard role is supported by our epistasis analysis using splicing inhibitors, which failed to phenocopy the AQR-deficient state (**Fig. S5D, E**), the direct observation of R-loop accumulation at the unspliced Airn locus (**Fig. S5F**), as well as the observation that exogenous hRNASEH1 expression suppresses DNA damage markers and partially restores proliferation of AQR-depleted cells (**Fig. Fig. 6J, K, S5G-I**).

A pivotal finding of this study is the link between the physical DNA template and transcriptional systems. We discovered that central pluripotency regulators, such as Sox2 and Klf4, are located within genomic hotspots prone to head-on TRCs (**Fig. 8C**). AQR depletion exposes these critical loci to replication interference, leading to localized transcriptional instability. The analysis of transcriptional entropy (ID) shows that AQR loss expands the accessible transcriptional state space, increasing "entropy" or plasticity (**Fig. 9E, F**). Interestingly, this does not manifest as a global increase in noise, but rather as a targeted redistribution of variability. Core identity genes, including Oct3/4, and Sox2, show significantly increased transcriptional noise specifically in naive mESCs upon AQR loss (**Fig. 9H, I**). This establishes a direct mechanistic connection: the collapse of R-loop resolution at conflict-prone loci triggers stochastic fluctuations in key transcription factors, which in turn facilitates the drift of the population toward alternative cell fates, such as the 2CLC or lineage-primed states (**Fig. 9C, D**).

Taken together, our study positions T–R coordination as a quantifiable property of cell state that integrates genome stability with developmental potency. Our findings challenge the “avoidance” model of genome maintenance, suggesting that stem cells have adapted a "high-risk, high-resolution" strategy. From an evolutionary perspective, the localization of key pluripotency genes within TRC-prone regions may offer a decisive advantage for developmental plasticity. By placing identity-defining loci in regions of high transcription-replication overlap, embryonic cells may utilize this inherent mechanical fragility as a regulatory mechanism for efficient cell-fate transitions. Under this model, the high-traffic environment acts as a poised state and as long as resolution factors like AQR are active, identity is maintained; however, any developmentally programmed attenuation of the resolution machinery would lead to immediate, stochastic fluctuations at core regulatory loci, providing the necessary "noise" to break symmetry and initiate lineage commitment. Along these lines, our findings raise significant questions for regenerative medicine and cancer biology. Given that somatic reprogramming needs to reinstate a “resilient” replication mode, could the high-traffic signature of hiPSCs make them uniquely susceptible to specific classes of replication-stress-inducing drugs? Future work should investigate whether this resilient mode is a universal feature of early embryogenesis *in vivo* and how it is bypassed during the first stages of lineage commitment.

## Materials and Methods

### Materials

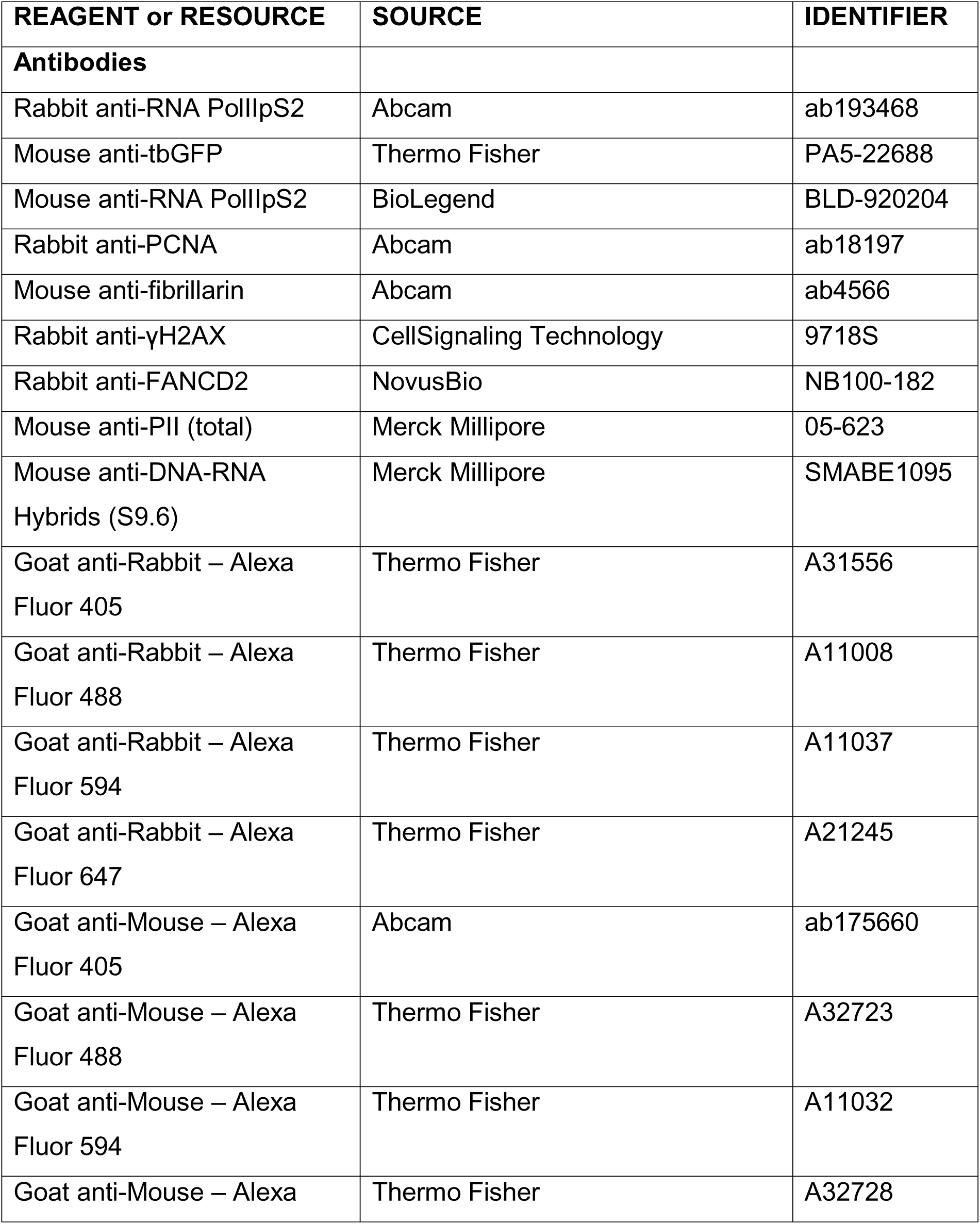
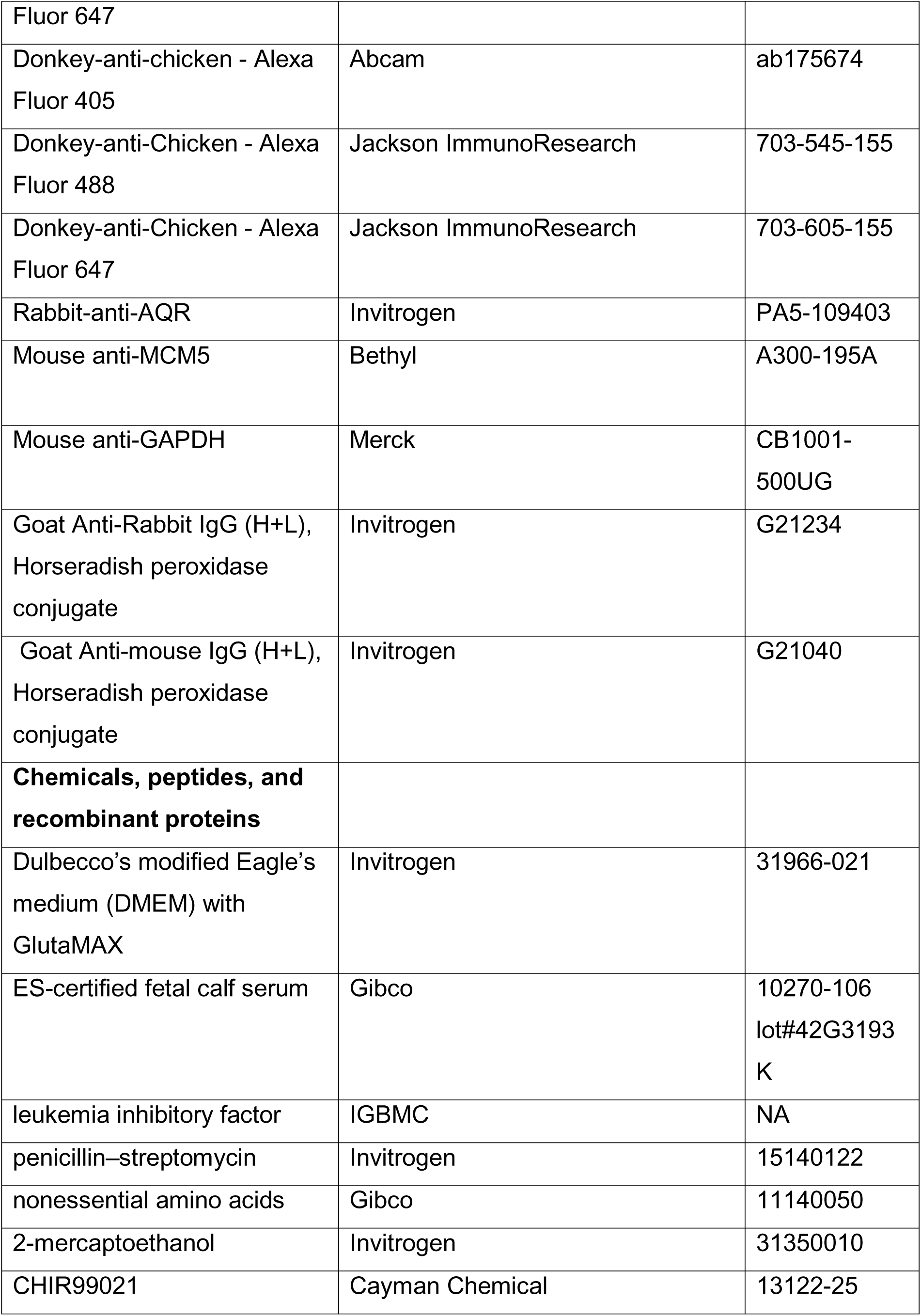
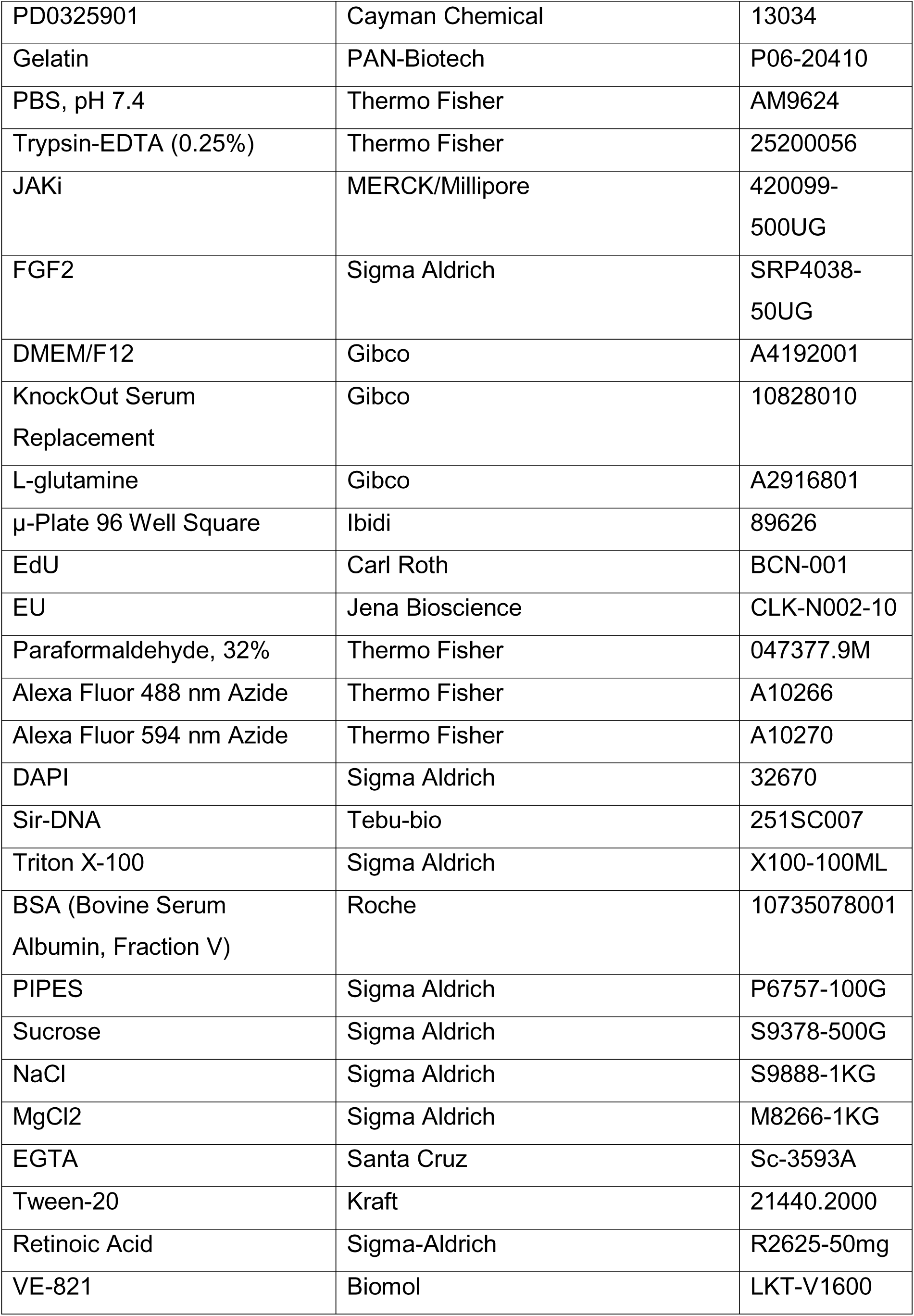
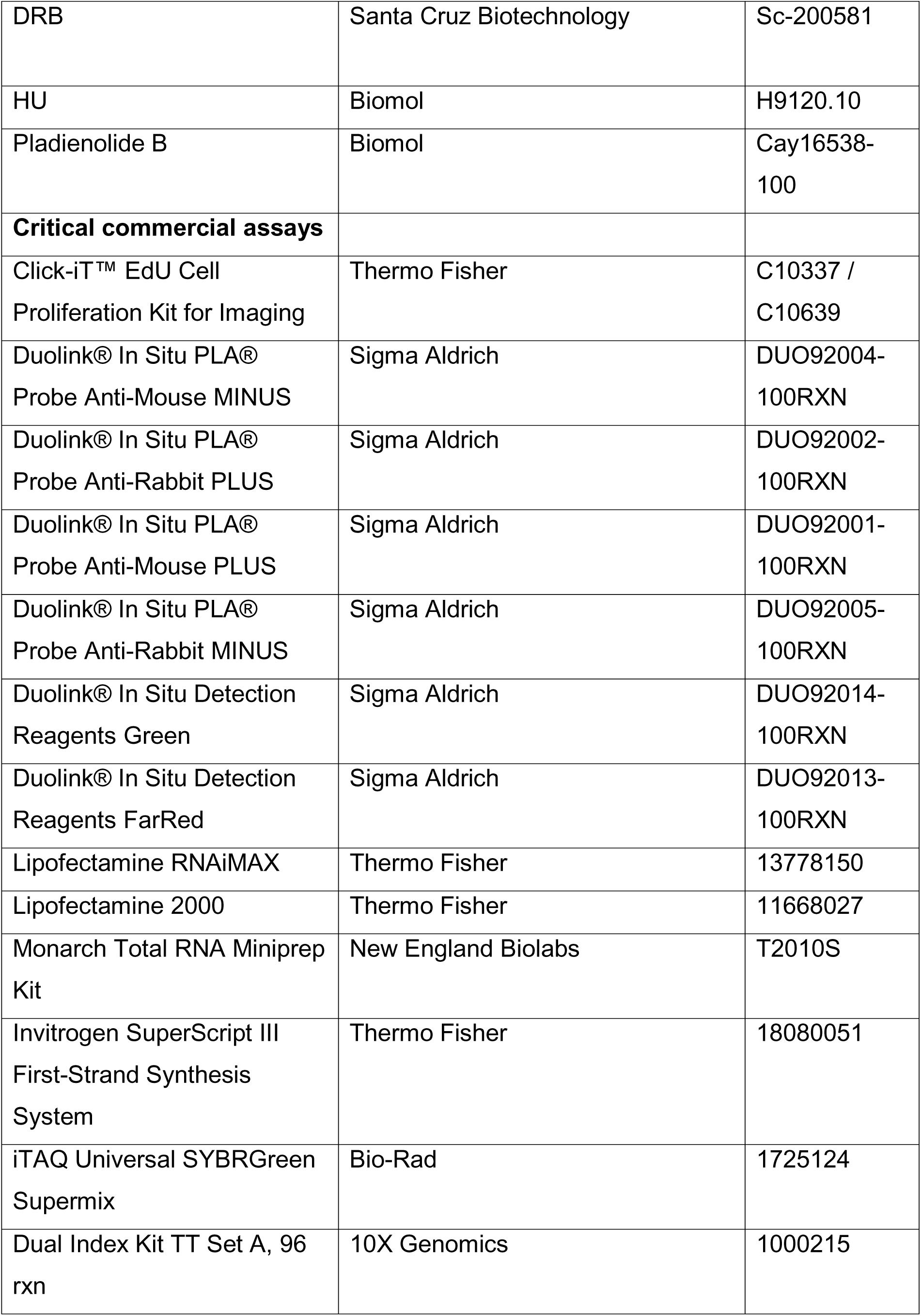
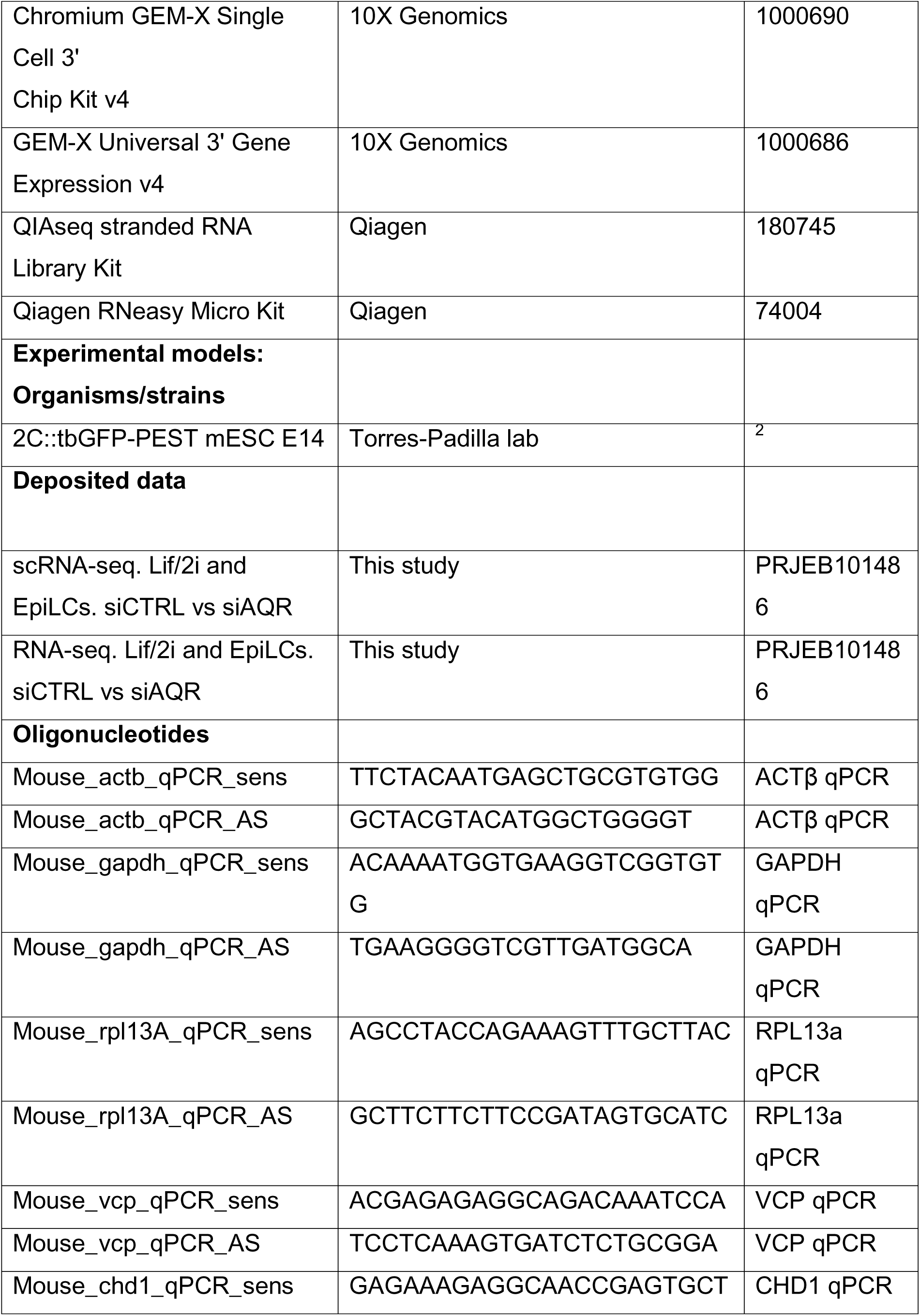
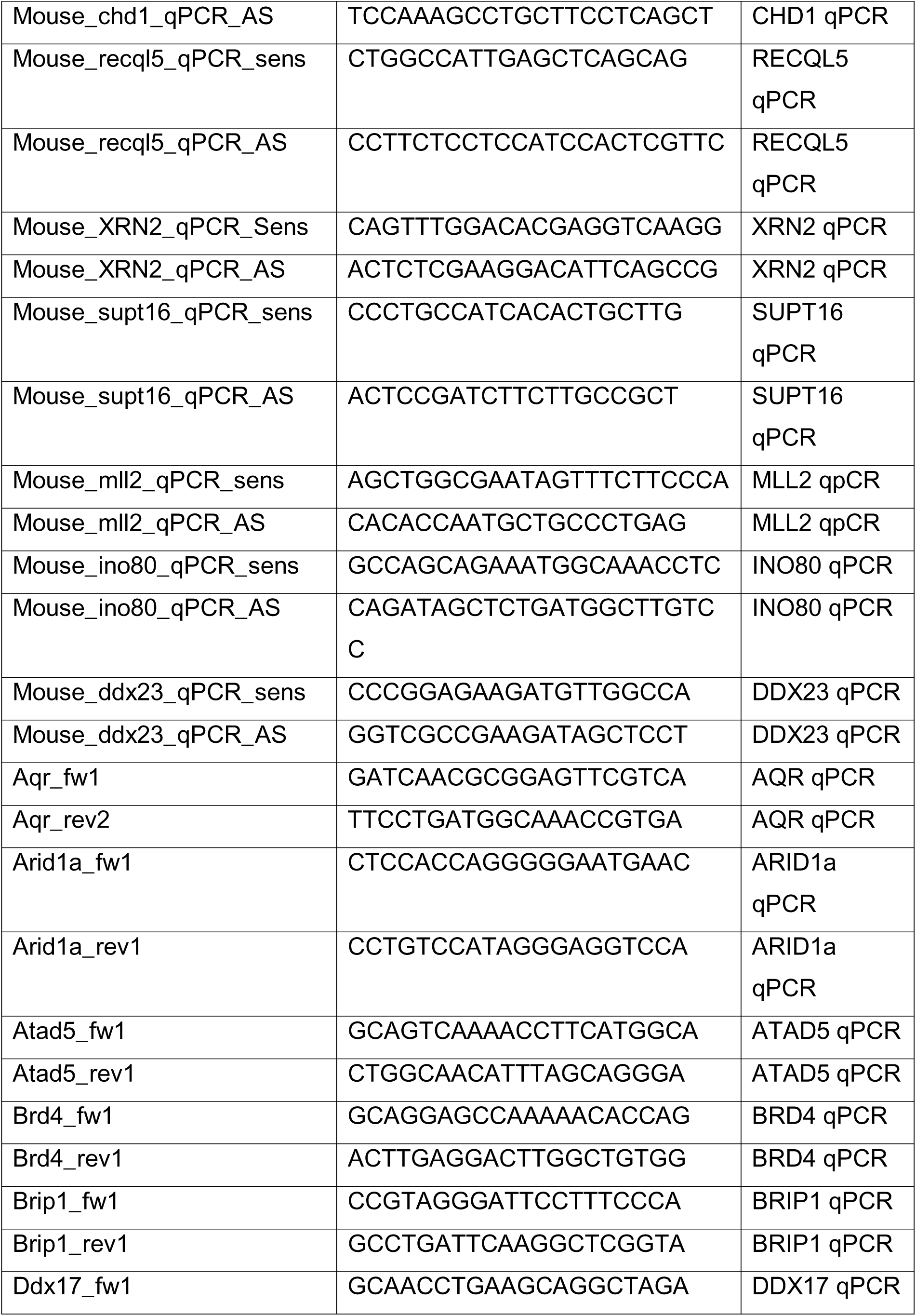
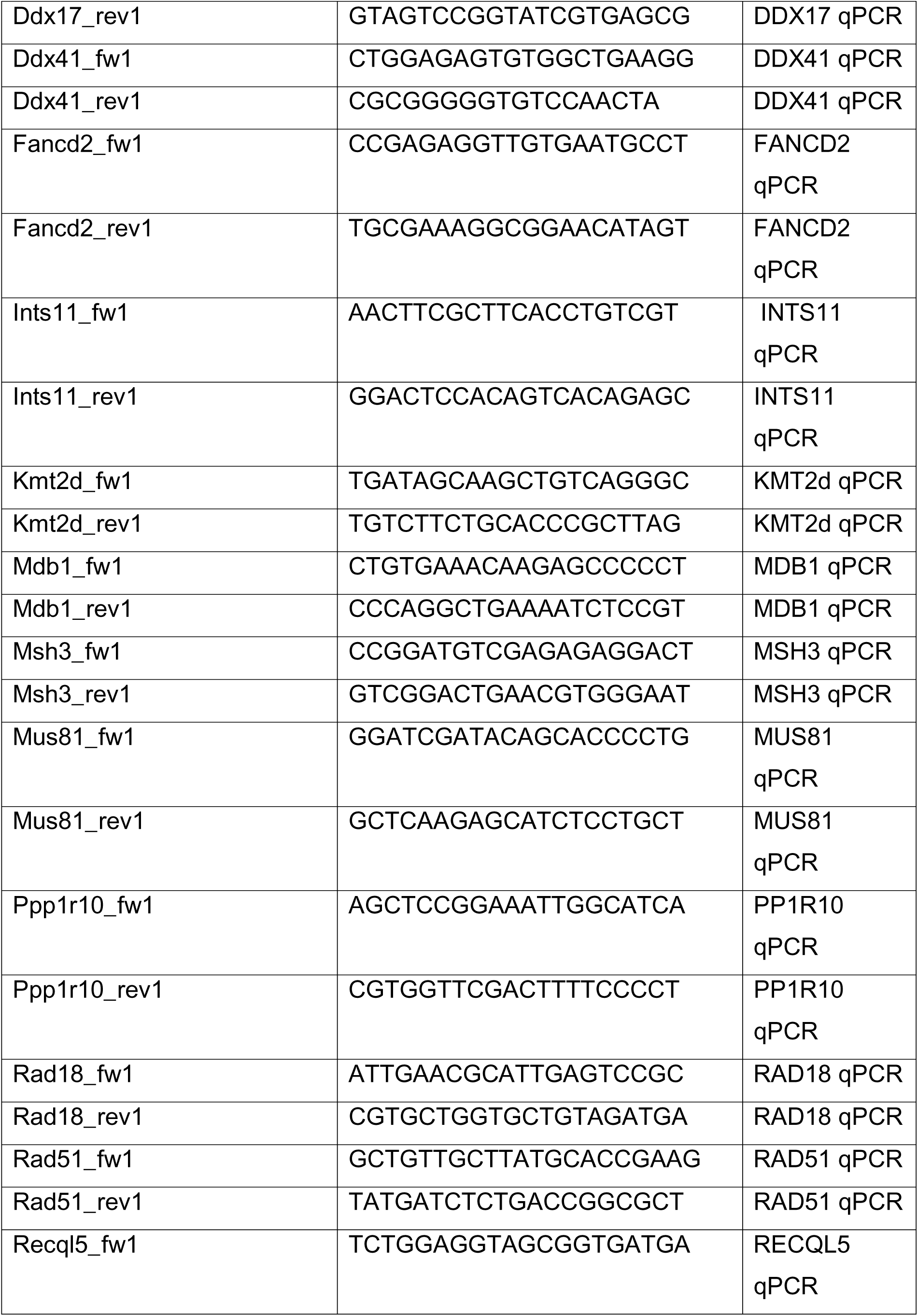
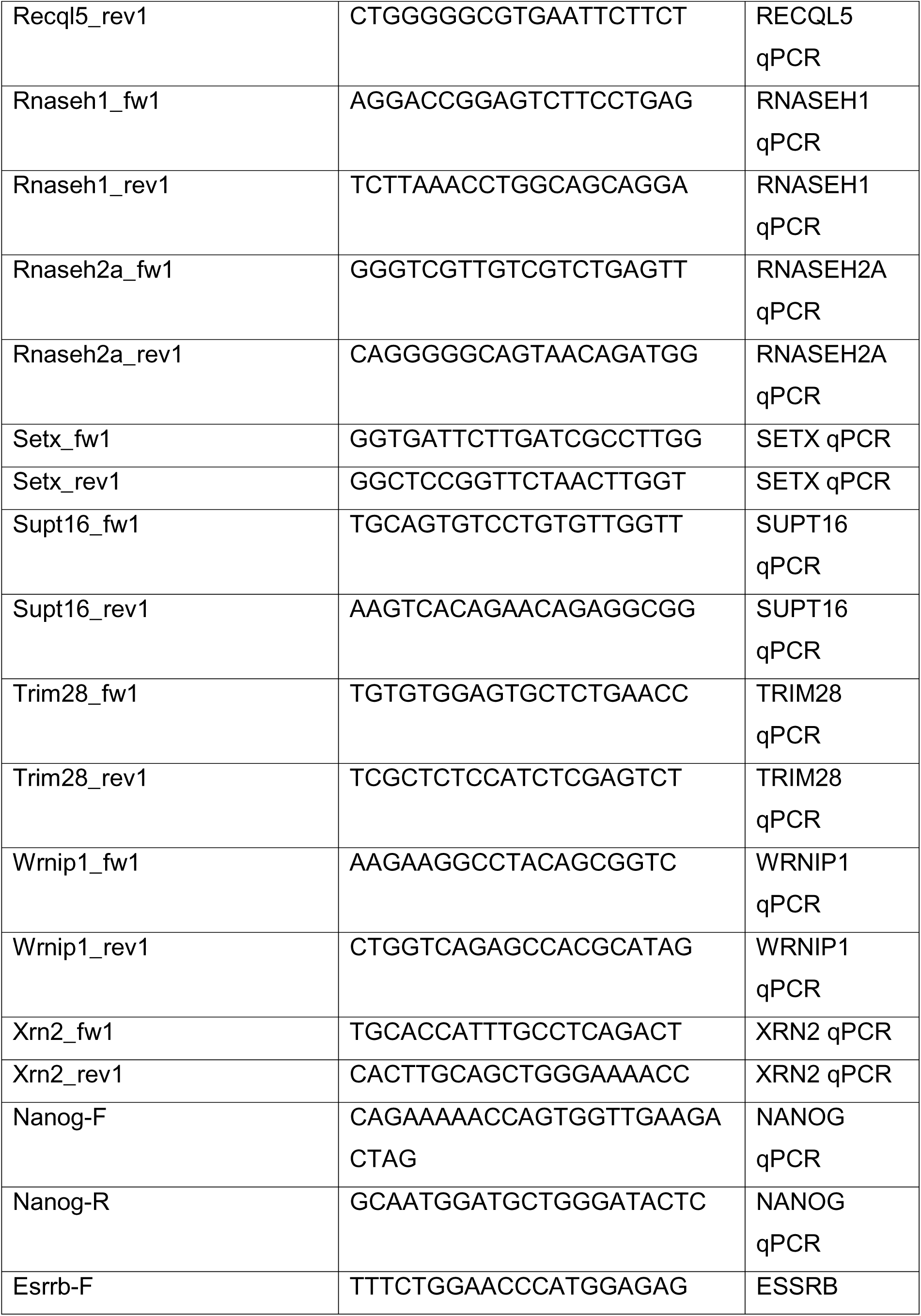
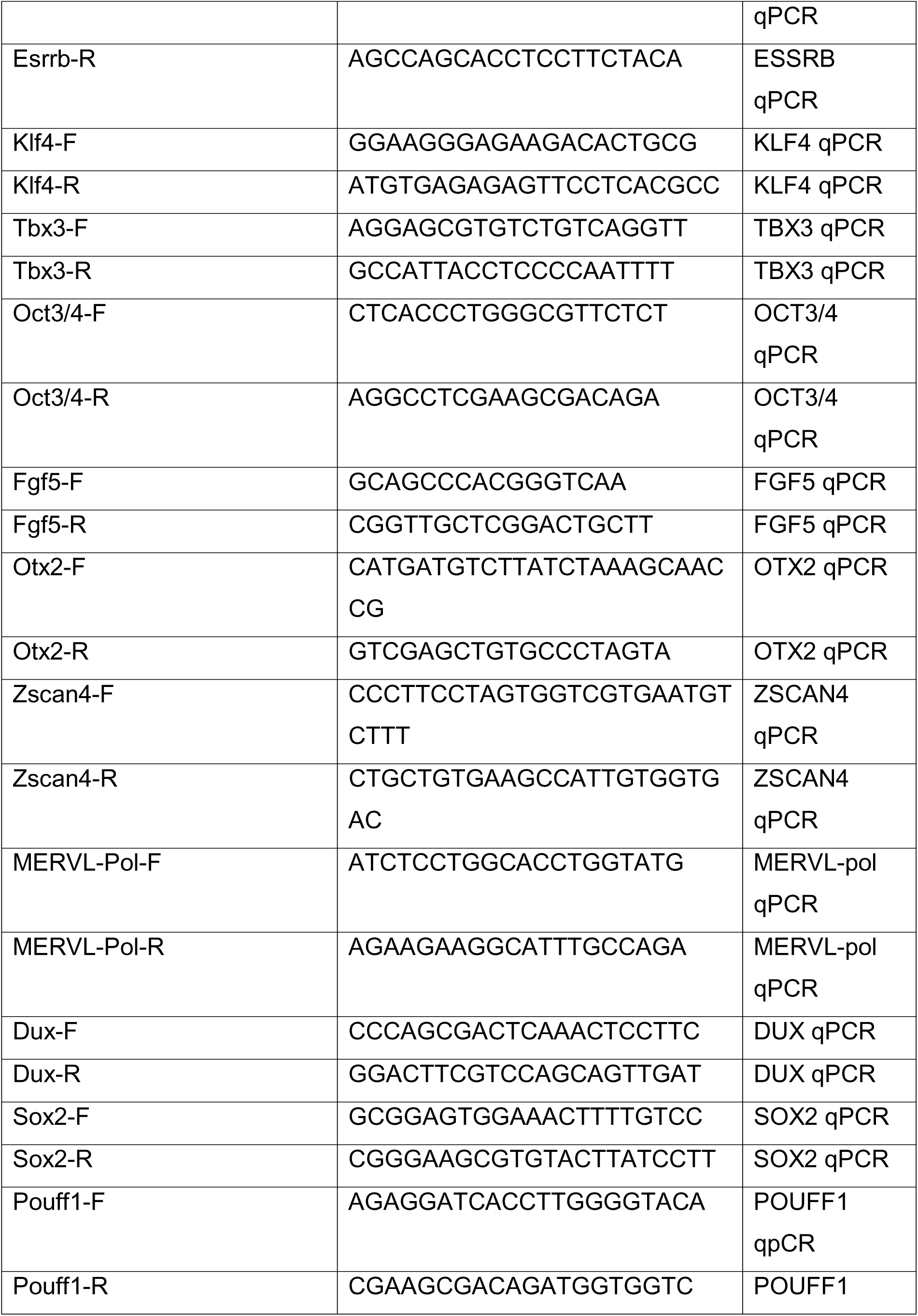
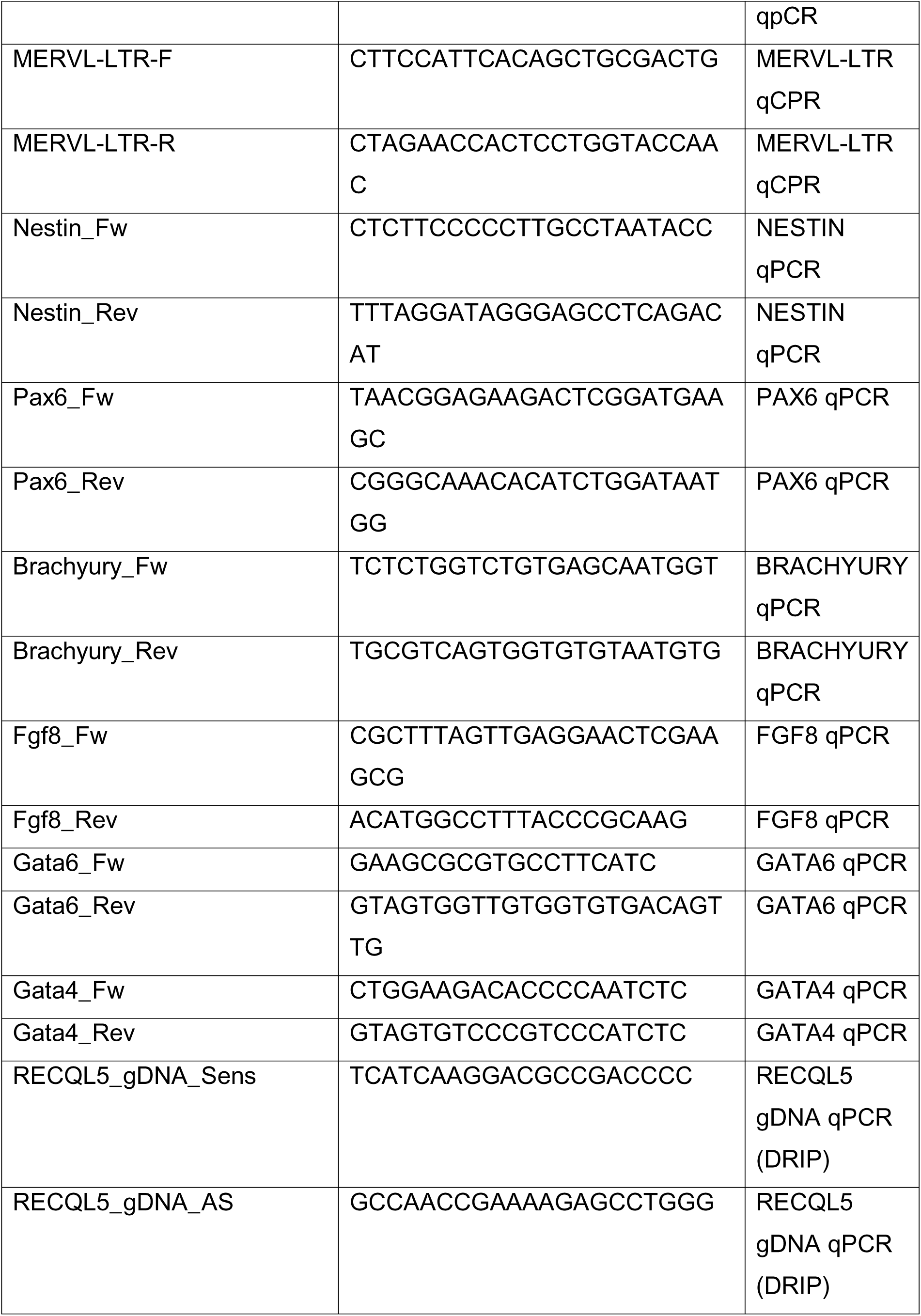
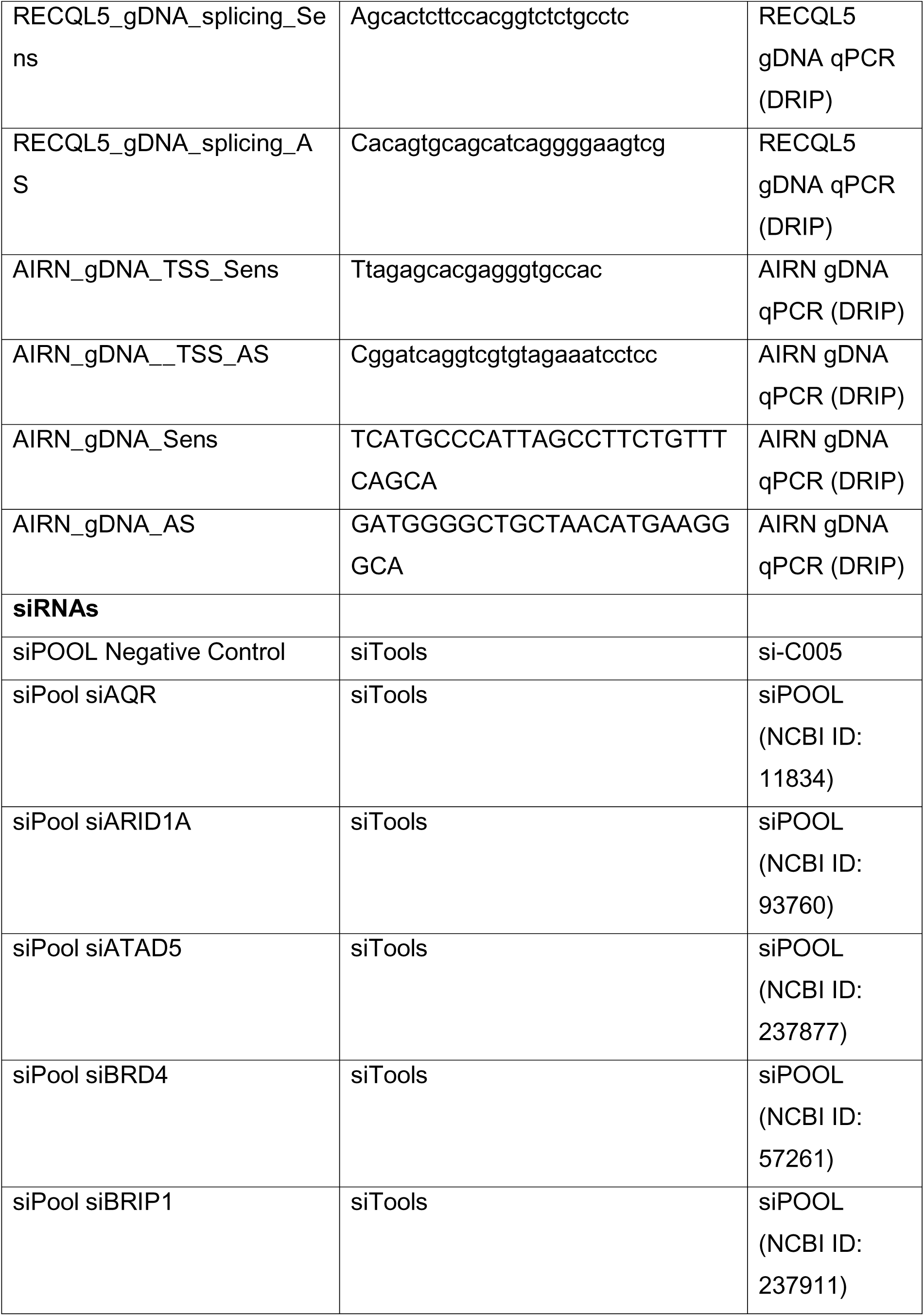
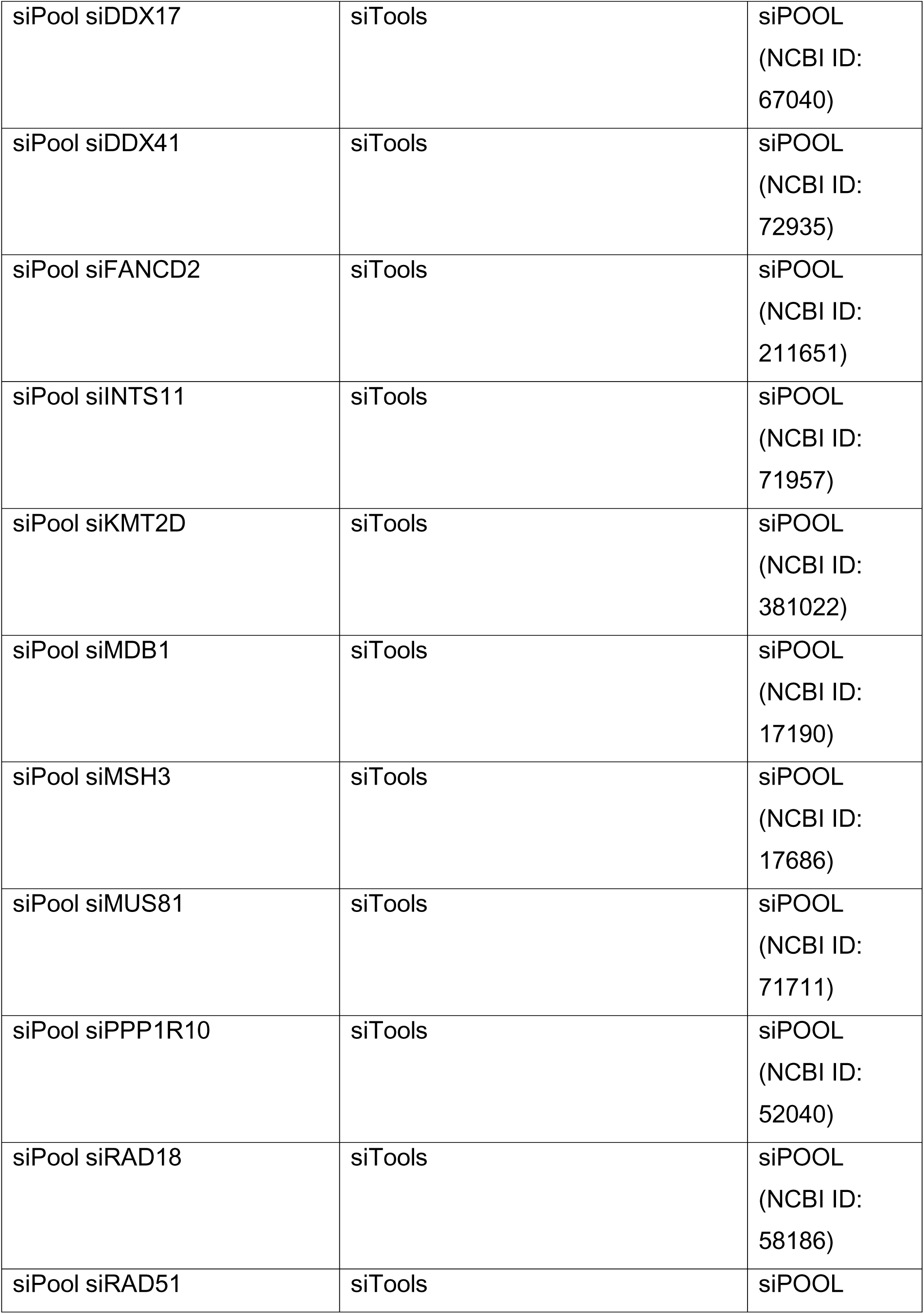
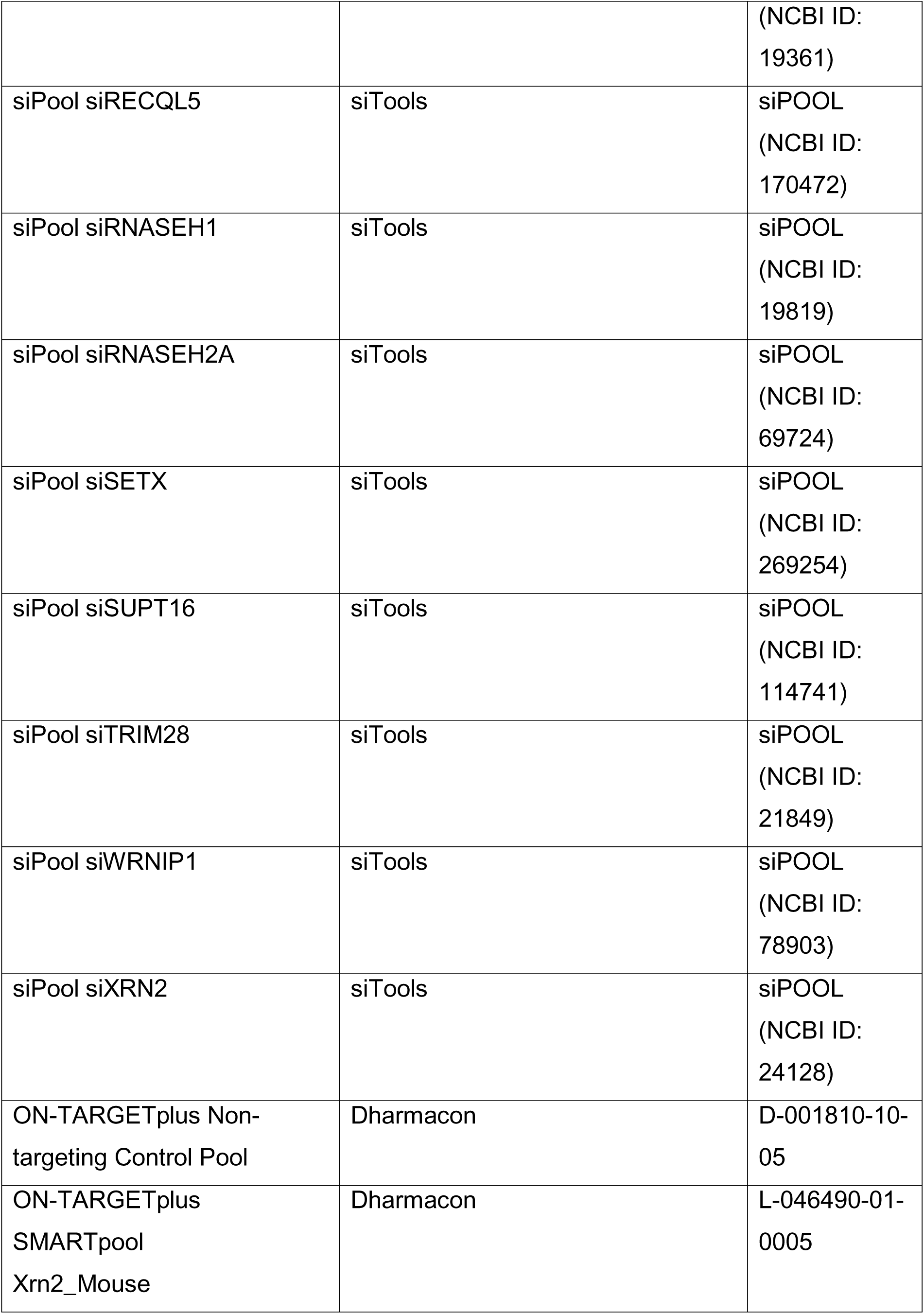
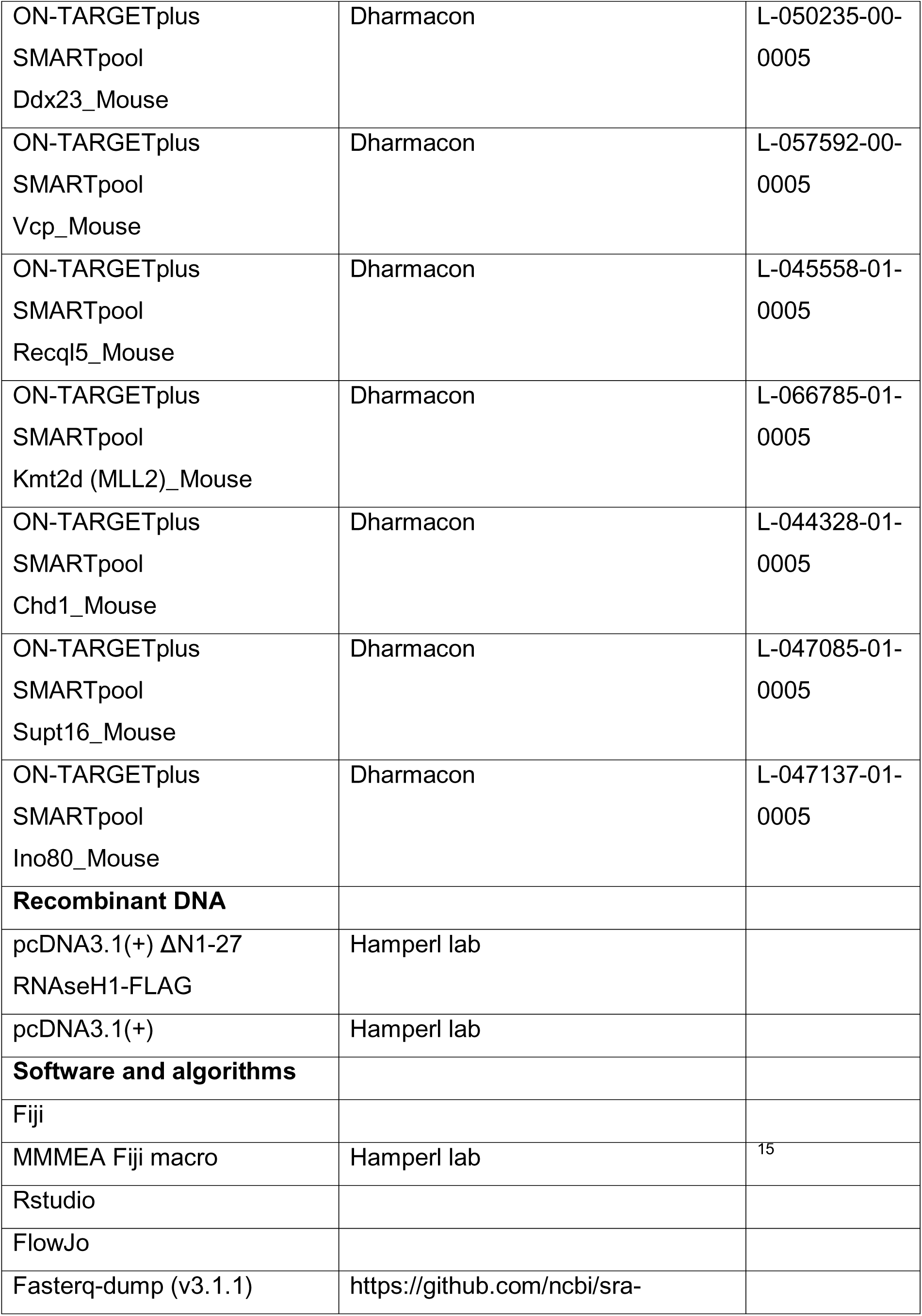
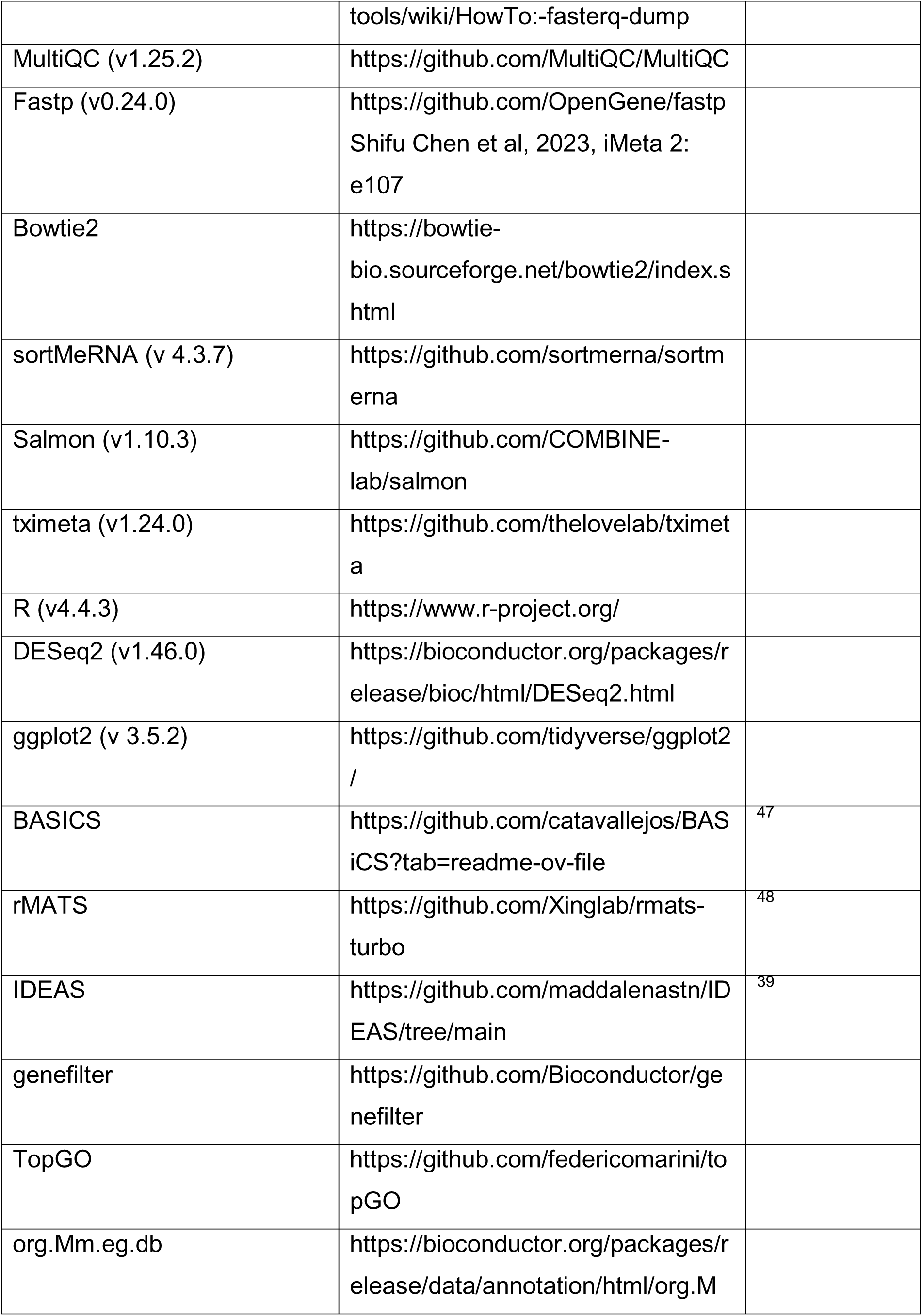
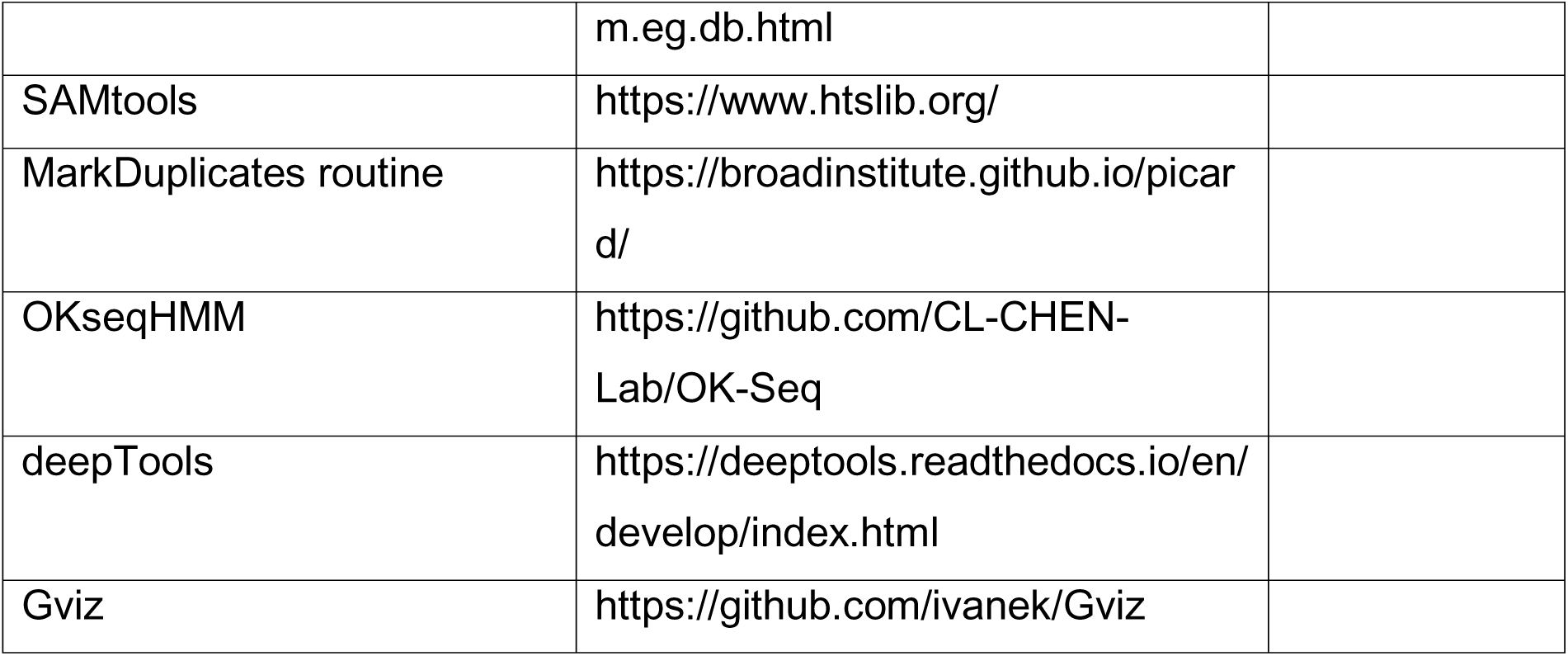

### Methods

#### Cell culture

mESC lines (E14) were cultured in 2i media (DMEM-GlutaMAX (Dulbecco’s modified Eagle’s medium, high glucose, GlutaMAX Supplemented) (DMEM) containing 15 % ES-certified fetal calf serum, 2× leukemia inhibitory factor (Lif), 100 units/mL penicillin–streptomycin, 0.1 mM nonessential amino acids, 0.1 mM 2-mercaptoethanol, 3 μM CHIR99021 (GSK3β inhibitor) and 1 μM PD0325901 (MEK inhibitor)) on gelatin-coated plates. Media was changed daily and passaged every 2-3 day. Primary MEFs were cultured in the same media but without Lif, CHIR99021, and PD0325901. To induce mEpiLCs, mESCs were treated as in ^49^. Briefly, cells were cultured in standard 2i media until mEpiLC induction. Then, the media was changed for -Lif media (DMEM-GlutaMAX (Dulbecco’s modified Eagle’s medium, high glucose, GlutaMAX Supplemented) (DMEM) containing 15% ES-certified fetal calf serum, 100 units/mL penicillin–streptomycin, 0.1 mM nonessential amino acids, 0.05 mM 2-mercaptoethanol) for 24 h and then again for EpiLC media for 48 h (DMEM/F12 supplemented with 20% knockout serum replacement, 2 mM L-glutamine, 1% penicillin/streptomycin, 0.1 mM 2-mercaptoethanol, 0.1 mM nonessential amino acids, 30 ng/ml FGF2, and 0.6 μM JAKi).

#### Drug treatments

Retinoic Acid (RA) was used as a 2CLC inducing control condition. RA treatment was performed by adding RA to a final concentration of 10 µM for 48 h prior to FACS analysis. The CDK9 inhibitor 5,6-Dichloro-1-β-D-ribofuranosylbenzimidazole (DRB) was used as a control condition to validate the functionality of the overlap analysis by Manders’ correlation coefficient (MCC). For DRB treatment, DRB was added at a final concentration of 100 µM for 1h prior to pre-extraction and fixation. EdU short 15 min pulse was performed by adding EdU in the DRB containing media. ATR inhibition was performed by adding the ATR inhibitor VE-821 at a final concentration of 10 µM for 4 h or 24 h. Hydroxyurea (HU) was used together with siAQR in FACS experiments. HU treatment was performed by doing siRNA transfection in media containing HU to a final concentration of 50 µM. Pladienolide B (PlaB) was also used together with siAQR in FACS experiments. PlaB pulse-chase was performed 24 h after siRNA transfection by adding PlaB for 4 h at a final concentration of 2.5 nM or 20 nM, washing it with fresh media, and letting cells grow for an additional 24 h.

#### siRNA transfection

24 h prior to siRNA transfection, 2i is removed from the media except when specified (i.e.: siRNA in naive maintained mESCs or siRNA during mESCs priming). Then, 200 000-500 000 mESCs are reverse transfected using Lipofectamine RNAiMAX according to the manufacturer’s protocol. Briefly, 7,5 µL of Lipofectamine in OptiMEM media (total volume:125 µL) is added to empty wells. In parallel, siRNA was diluted in Opti-MEM (total volume:125 µL), combined with the Lipofectamine mix, and incubated for 15 min. For siRNAs purchased from siTools, the final concentration of siRNA used varied between 30-60 nM. For siRNAs purchased from Dharmacon (Fig. S4E, F: siDDX23, siSUP16, siVCP, siMLL2, siINO80, siCHD1, siRECQL5, siXRN2), the final concentration of siRNA used was 20nM. Cells were then seeded directly onto the transfection mixture in 2 mL of medium lacking 2i (unless specified) and cultured for 48 h, with daily medium changes. For microscopy experiments, cells were trypsinized after 24 h of transfection and reseeded into 96well plates.

#### Plasmid transfection

Confluent mESCs cultured in 6well format were transfected using Lipofectamine 2000 according to the manufacturer’s protocol. Briefly, 10 µL of Lipofectamine 2000 was added to OptiMEM media (total volume:250 µL) In parallel, 4µg of plasmid DNA was diluted in Opti-MEM (total volume:250 µL), and then combined with the Lipofectamine mix, and incubated for 15 min. The transfection was applied on the cells and cultured for 48h, with daily medium change. For Plasmid and siRNA experiments, cells were transfected with plasmid 12h prior to siRNA transfection.

#### FACS

ESCs were harvested with 0.25% trypsin, resuspended in ESC medium, and maintained on ice. Dead cells, debris, aggregates, and doublets were excluded based on forward- and side-scatter profiles. 2CLC were identified by the fluorescence of the tbGFP reporter. Sorting was carried out on a FACS Melody (BD Biosciences), data analysis was performed using the FlowJo software.

#### Immunofluorescence (IF)

Cells were seeded in gelatine-treated ibidi 96well plate at 20-50 000 cells in 200 µL of media per wells and were cultured for 12-24 h. If needed, the media was replaced by fresh media containing 10 µM EdU for 15 min to perform a short EdU pulse, followed by 2 washes with PBS. To perform an EU pulse, the media was replaced by fresh media containing 500 µM EU for 30 min, followed by 2 washes with PBS. Unless stated otherwise, a pre-extraction step was performed before fixation. Media/pre-extraction buffer was replaced by CSK buffer (10 mM PIPES, 300 mM sucrose, 100 mM NaCl, 3 mM MgCl2, and 1 mM EGTA) + 0.5 % Triton X-100 for 15 min at room temperature, followed by 2x wash with PBS. Then, cells were fixed using a 4% PFA solution in PBS for 20 min at room temperature, washed twice using PBS, permeabilized for 1 h at room temperature using CSK buffer + 0.5 % Triton X-100, and then washed again 2x with PBS. If an EdU pulse was performed, a Click-it reaction was performed at this point (see below).

The wells were then blocked with a 3% BSA PBS solution for 1h at room temperature or overnight at 4°C. Primary antibodies incubations were performed overnight at 4°C in 3% BSA PBS, and then three 5 min washes in PBS + 0.1 % Tween-20 were performed. Secondary antibodies and DNA staining (DAPI (1:2000) or sirDNA (1:5000)) incubations were performed at room temperature for 1 h in 3% BSA PBS and then washed 3 times with PBS + 0.1 % Tween-20. Wells were kept in PBS until imaged.

The primary antibodies used and their dilution are: rabbit anti-RNA PolIIpS2 (1:500) for; mouse anti-tbGFP (1:500); mouse anti-RNA PolIIpS2 (1:500) (used for double IF against PIIpS2); mouse anti-fibrillarin (1:200); Rabbit anti-γH2AX (1:500); Rabbit anti-FANCD2 (1:500).

All AlexaFluor conjugated secondary antibodies against mouse and rabbit were used at a 1:1000 while the IF secondary antibodies against chicken was used at a 1:500.

#### Nascent DNA labelling - EdU fluorophore conjugation

To label nascent DNA replication, a short EdU pulse is performed as stated previously. Then, following cell fixation, EdU is then conjugated with an AlexaFluor fluorophore. To do so, a Click-it^TM^ reaction was performed using the manufacturer’s protocol. Alternatively, an equivalent chemical conjugation reaction was performed with custom reagents. Briefly, cells were incubated for 30 min at room temperature with 70µL of labelling reaction cocktail (100 mM Tris-HCl pH 8.5, 1 mM CuSO_4_, 100 mM Ascorbic acid, 13 µg/mL Alexa Fluor Azide (488nm or 594nm)) followed by 3x PBS washes.

#### Proximity Ligation Assay (PLA)

Proximity Ligation Assays were performed as described in ^15^ and as per the manufacturer’s protocol. Briefly, cells were incubated overnight at 4°C with a mouse anti-RNA PIIpS2 and a rabbit anti-PCNA (1:2000 in 3% BSA PBS) primary antibodies in a to detect TRCs. Then, cells were washed twice with PBS and incubated with the PLUS and MINUS probes (diluted at 0.5 X in the Duolink Antibody Diluent) for 1h at 37°C in a humidity chamber. Cells were then washed twice with Wash Buffer A, followed by a 30 min ligation reaction at 37°C in a humidity chamber, 2 washes with Wash Buffer A, a 100 min amplification reaction at 37°C in a humidity chamber, 2 washes with Wash Buffer B, and finally cells were left in 200 µL of PBS until imaging. If PLA was combined with IF and/or DNA staining, they were performed after the PLA reaction.

#### Imaging

Following IF, EdU labelling, and/or PLA, microscopy images were acquired in a semi-automatized high-throughput fashion on a Nikon T2 inverted microscope with an Andor Dragonfly spinning disk, combined with a Plan Apochromat 40× air objective with a numerical aperture of 0.95, an iXon Life 888 EMCCD camera, and an additional 2.0× magnification lens, resulting in an effective image pixel size of 162.5 × 162.5 nm. 67–400 z-stack (11 steps, 1 µm per step) images were acquired per well.

#### Image analysis

Image analysis was performed using a previously published ImageJ macro^15^, followed by custom R scripts for downstream analysis. Briefly, the ImageJ macro extracts a single focal plane from Z-stacks by identifying the plane with the highest variation in DNA-staining pixel intensity^50^, corresponding to the middle of each nucleus. Background subtraction is applied, and nuclei are segmented using the Stardist neural network plugin ^51^ trained on DAPI-stained mESC nuclei. Nuclei that are out-of-focus are identified by comparing DNA staining in adjacent Z-planes and removed. Shape, size, and fluorescence parameters are then measured, and if necessary, the ImageJ “Find Maxima” tool is used to count foci. Downstream processing in R includes filtering out nuclei with abnormal size or DNA intensity and classifying nuclei as EdU+ or EdU- by generating a EdU mean fluorescence frequency distribution and identifying the intersection point between low- and high-EdU mean fluorescence intensity. 2CLCs were manually identified from the mESC population using scatter plots of tbGFP mean fluorescence intensity versus its coefficient of variation, analogous to FACS-based gating. Signal intensities are reported as robust Z-scores, calculated relative to the median and standard deviation of the reference population (mESCs for cell type comparisons, or non-treated/siCTRL for treatment comparisons). For comparisons of EdU+ and EdU- populations, only EdU+ cells are used to calculate the median and SD.

#### T-R overlap analysis

PIIpS2 and EdU fluorescence channels were first thresholded in ImageJ to define positive signal pixels, with sub-threshold values set to zero. Channel overlap was then quantified using the EZColocalization plugin^52^, operating on thresholded images with negative values excluded. Because the Pearson correlation coefficient (PCC) and related correlation-based metrics primarily capture linear relationships between pixel intensities, they were considered unsuitable for quantifying physical overlap between PIIpS2 and EdU signals (**Fig S2A, B**). Instead, we employed Manders’ colocalization coefficient (MCC), which directly measures the fraction of one signal overlapping with the other and is independent of intensity correlation, providing a more appropriate metric for assessing transcription–replication spatial overlap. In addition, although the reciprocal measurement of the overlap of EdU with PIIpS2 (EdU-PIIpS2 MCC) showed the same trends of PIIpS2-EdU MCC, we found that this readout was more sensitive to outlier cells with very few EdU-positive pixels (such as late S-phase cells) and decided to keep the more robust PIIpS2-EdU MCC as the default measured parameter (**Fig S2A, B**).

To validate this readout, we confirmed the expected anticorrelation between nuclear PIIpS2 and nucleolar Fibrillarin signals (MCC 0.2-0.4) and strong positive correlation between two simultaneous PIIpS2 stainings (MCC 0.8-1.0). PIIpS2-EdU MCC values as the readout for T-R coordination exhibited a broad distribution that significantly decreased following transcriptional inhibition with 5,6-Dichloro-1-β-D-ribofuranosylbenzimidazole (DRB) (**Fig. S2D, E)**. These results demonstrate that our readout effectively captures dynamic changes of T-R coordination.

To generate bubble plots and heatmaps for MCC evaluations, outliers (top and bottom 2.5%) were first excluded. To control for sample-size bias, cell numbers within each experiment were downsampled to match the smallest population. Signal intensities were then rescaled per experiment to a 0–100 range to allow comparison between experiments and binned into 10 groups along each axis, generating a two-dimensional grid of signal intensity bins. For each bin, the median MCC (or PCC) was calculated. Bubble size reflects the number of nuclei per bin, while bubble color represents the median MCC (or PCC), displayed on a blue–white–red gradient from low to high values.

#### Western Blot

To prepare whole-cell extracts, 1 × 10^6^ cells were lysed in 100μl RIPA buffer (50 mM Tris-HCl, pH 7.4, 150 mM NaCl, 0.5% deoxycholate, 0.1% sodium dodecyl sulfate, 1% NP-40, 1× protease inhibitor cocktail). Whole-cell extracts were sonicated alternating 2.5 sec on, 2.5 sec off for 10 min (Bioruptor UCD-200) before separation by electrophoresis and transfer onto polyvinylidene difluoride membranes. Membranes were blocked in 5% skimmed milk dissolved in 0.1% Tween/PBS overnight at 4°C. Membranes were incubated with primary antibodies dissolved in 3% BSA/PBS overnight at 4°C followed by washing thrice with 0.1% Tween/PBS. HRP-linked secondary antibodies were then added for 1 h at 25°C and washed thrice prior to signal detection using chemiluminescence. Primary antibodies: AQR (1:200), MCM5 (1:10,000), FLAG (1:500), GAPDH (1:10,000).

#### DNA:RNA immunoprecipitation (DRIP)

DRIP was performed as described ^53^ with minor changes. Briefly, DNA was extracted with phenol/chloroform, precipitated with EtOH/sodium acetate, washed with 70% EtOH, and resuspended in 120 μl TE buffer. DNA was sonicated with 10% duty factor, 200 cycles/burst, 140 peak incident power for 4 min (Covaris). For RNase H-treated samples, 4 μg of DNA was treated with 10U *E. coli* RNaseH (NEB #M0297S) overnight at 37°C. DNA was purified by phenol/chloroform, EtOH/sodium acetate precipitation as described above. In total, 4 μg of DNA was bound with 7 μg of S9.6 antibody in 1× binding buffer (10 mM NaPO4 pH 7, 140 mM NaCl, 0.05% Triton X-100) overnight at 4°C. Protein A/G agarose beads were added for 2 h. Bound beads were washed 3 times in binding buffer, and elution was performed in elution buffer (50 mM Tris pH 8, 10 mM EDTA, 0.5% SDS, Proteinase K) for 45 min at 55°C. DNA was purified as described. Quantitative PCR of immunoprecipitated DNA fragments was performed on a Roche LightCycler 480 Instrument II using SYBR-Green master mix (Biorad).

#### RNA extraction and RT-qPCR

Total RNA was extracted using the Monarch Total RNA Miniprep Kit (NEB) according to the manufacturer’s instructions. RNA concentration and purity were determined by spectrophotometry. For cDNA synthesis, 2 µg– 5 µg of total RNA was reverse-transcribed using the SuperScript III First-Strand Synthesis System (Invitrogen) with random hexamers. Quantitative PCR was performed using iTaq Universal SYBR Green Supermix (Bio-Rad) on a LightCycler PRO (Roche) with a standard 3-step qPCR program (60°C annealing; 45 cycles). Relative gene expression levels were calculated using the ΔΔCt method, with ACTINβ, GAPDH, and RPL13a as reference genes, and either mESCs, Lif 2i, non-treated, or siCTRL samples as the calibrator condition.

#### Bulk RNA-seq

##### Data acquisition of publicly available datasets and analysis

Publicly available bulk RNA-seq datasets were obtained from the NCBI Sequence Read Archive (SRA). Data were downloaded from three BioProjects: PRJNA483504^54^, PRJNA669873^21^, and PRJNA299625^20^, which together included samples from mouse embryonic stem cells, 2C-like cells, epiblast-like cells, and fibroblasts. Raw sequencing data (FASTQ files) were retrieved with the fasterq-dump (v 3.1.1) software from SRA Toolkit ^55^, individual run accessions for each dataset are listed in Supplementary table I.

Read quality was assessed using MultiQC (v1.25.2)^56^. Adapter trimming and quality filtering were performed using fastp (v0.24.0)^57^ with default parameters. To minimize ribosomal RNA contamination, reads mapping to rRNA were removed with sortMeRNA (v 4.3.7)^58^ using default databases for the corresponding version.

Transcript-level abundances were estimated with Salmon (v1.10.3) by pseudo-mapping to the Mus musculus GRCm38 transcriptome from Ensembl release 102^59^ as reference. Salmon outputs were summarized to the gene level using tximeta (v1.24.0) in R (v4.4.3)^60^.

Gene-level count matrices were analysed using DESeq2 (v1.46.0) in R. Normalization was performed using DESeq2’s default median-of-ratios method. Differential expression was assessed using the Wald test, and p-values were adjusted using the Benjamini-Hochberg false discovery rate (FDR) procedure. Genes with adjusted p-value < 0.05 and |log_2_ fold-change| > 0 were considered significantly differentially expressed. MA-plots were generated with ggplot2 (v 3.5.2)^61^ to visualize global patterns of differential expression.

##### TRC prone region prediction

We employed the described approach in^38^ for the estimation of significant changes in the abundance of factors at R-loop enriched TRC site in respect of the surrounding region. We first retrieved OK-seq public data from the NCBI Sequence Read Archive (SRA) database under accession code SRR7535256^62^, and public DRIP-seq with accession number SRR2075686^63^ to generate the potential TRC sites in silico. ChIP-seq datasets for factors of interest were retrieved as listed in the Data availability section.

The OK-seq and ChIP-seq reads were evaluated for quality control and processed by Fastp (v0.23.4)^64^ and then aligned to Mus musculus GRCm38 genome from Ensembl release 101 as reference using Bowtie2 (v2.5.3)^65^, with parameters “--end-to-end --very-sensitive --no-unal -q”. Uniquely mapped reads with mapping quality ≥ 12 were retained using SAMtools (v1.20)^66^. Duplicate reads were discarded by Picard tool’s MarkDuplicates routine (v3.1.1)^67^. OKseqHMM (v2.0.0)^68^ was used to calculate replication fork directionality (RFD) scores from the processed OK-seq data, regions within > |0.17| were retained and annotated for all gene bodies located inside them. Then, genes were overlapped with DRIP-seq data to define TRC sites. Based on the identified regions, the windows were re-defined by centering the site using the closest beginning and end positions of the overlap and then extending +/- 10Kb. For each TRC site, control regions upstream and downstream were generated with 1Mb distance and 20Kb length. ChIP coverage tracks were normalized by Reads Per Kilobase per Million mapped reads (RPKM), using bamCoverage routine from deepTools (v3.5.5)^69^], with arguments “--binSize 100 --normalizeUsing RPKM”. Replicates were merged using bigwigAverage from deepTools to produce an aggregate signal by ChIP experiment. Genome tracks were displayed using Gviz R package (v1.50.0)^70^.

##### RNA-seq library preparation

RNA-seq libraries were prepared from total RNA with rRNA depletion using the QIAseq stranded RNA Library Kit (Qiagen, 180745), according to manufacturer’s instructions. Total RNA was extracted using Qiagen RNeasy Micro Kit (Qiagen, 74004) according to manufacturer’s protocol. In brief, 1 ug total RNA was used as starting material, fragmentation time (95°C, 10 min) was optimized to generate 150-250bps fragments, and the libraries were amplified with 12 cycles of PCR. cDNA libraries were sequenced for 76 cycles in paired-end mode with dual 8 bp indexes, with 20M reads/sample., with 20M reads/sample.

##### Identification and quantification of alternative splicing defects

Alternative splicing defects were identified using Replicate Multivariate Analysis of Transcript Splicing (rMATS), which detects differential splicing events from RNA-seq data by comparing exon–junction usage between experimental conditions^48^. For each condition, rMATS was run in paired mode using aligned RNA-seq reads to quantify five classes of splicing events: skipped exons (SE), alternative 5′ splice sites (A5SS), alternative 3′ splice sites (A3SS), retained introns (RI), and mutually exclusive exons (MXE). Event-level output files corresponding to junction count-based statistics (JC) were parsed programmatically, and for each event the associated gene symbol and false discovery rate (FDR) were extracted.

Splicing defects were defined as events passing an FDR threshold of <0.1, applied independently for each splicing class. For each condition, significant splicing events were grouped by event type and reduced to non-redundant gene-level sets representing genes affected by at least one statistically significant splicing alteration of a given class.

##### Differential gene expression analysis

Differential expression results were obtained from DESeq2 (v1.50.0), and genes with adjusted p-value < 0.1 were selected and stratified by direction of regulation based on the Wald statistic sign. For each gene subset, enrichment was tested against a locally defined background derived from the expression space of the dataset.

To reduce annotation bias and local expression density artifacts, a custom background selection strategy was implemented: for each foreground gene, the 10 nearest neighbors in expression space (based on DESeq2 baseMean values and Manhattan distance) were identified using the genefilter package^71^, and the union of these neighbors was used to construct a context-matched background set excluding the foreground genes themselves.

Gene expression values were obtained from normalized RNA-seq counts averaged across three biological replicates per condition. For each gene, differential expression between siCTRL and siAQR was evaluated independently in Lif2i and EpiLC conditions. Log2 fold changes were calculated as log2(siAQR / siCTRL) using the mean expression values of each group. Statistical significance was estimated using two-sided Student’s t-tests performed on replicate-level expression values (n = 3 per group).

##### Gene ontology analysis

Biological process enrichment was assessed separately for upregulated and downregulated genes obtained from the differential expression analysis performed in our bulk RNA-seq data using the topGO Bioconductor package (v2.62.0)^72^, Ensembl gene identifiers, and the org.Mm.eg.db annotation package^73^. Both the “classic” and “elim” algorithms were applied using Fisher’s exact test, and results were ranked by classic p-values. Enriched GO terms were filtered based on stringent statistical criteria (Fisher.classic < 1×10⁻³ and Fisher.elim < 0.05), ranked by the ratio of observed to expected genes, and the top 25 terms were retained for visualization. Enrichment strength was expressed as the observed/expected ratio, statistical significance as −log₁₀(Fisher p-value), and term size as the number of significant genes contributing to each category.

##### Targeted Gene Set Overlap (GSO)

To systematically compare a query gene set against multiple curated gene lists, we implemented a targeted overlap analysis. A single gene list was selected and tested for enrichment against each remaining list individually. The background universe was defined as the union of all genes present across the imported gene sets, thereby restricting statistical testing to the experimentally relevant gene space. For each pairwise comparison, we computed the size of each gene set, the number of overlapping genes, and the Jaccard index as a measure of relative similarity. Statistical enrichment was assessed using Fisher’s exact test (one-sided, testing for overrepresentation), based on a 2×2 contingency table constructed from the query set, target set, and background universe. For each comparison, odds ratios and associated p-values were extracted, and multiple hypothesis testing correction was applied using the Benjamini–Hochberg procedure to control the false discovery rate (FDR).

##### Single-cell RNA-seq (10x Genomics) and sequencing

Mouse embryonic stem cells were harvested at ∼70–80% confluence, resuspended in 0.04% BSA/PBS and dissociated to single cells by pipetting. Cell suspensions were filtered through 40 µm strainers, washed twice, and assessed for viability by trypan blue (> 90%). Cells were resuspended at 800–1,400 cells/µL in 0.04% BSA/PBS and kept on ice until loading. Single-cell 3′ RNA-seq libraries were prepared using the Chromium Controller (10x Genomics) and Single Cell 3′ Gene Expression kitv4, following the manufacturer’s protocol (CG000000731). Briefly, cells were loaded per channel to target recovery of ∼20,000 cells. Gel bead-in-emulsions (GEMs) were generated, followed by reverse transcription, GEM breakage, and cDNA cleanup. cDNA was amplified for 11 cycles, enzymatically fragmented, end-repaired, A-tailed, adaptor-ligated, and sample-indexed by PCR (8–12 cycles). Library quality and size distribution were confirmed by Bioanalyzer/Fragment Analyzer, and concentrations were determined by Qubit/RT-qPCR. Pooled libraries were paired end sequenced at the Helmholtz Munich Sequencing Core on an Illumina NovaSeq X using a 10B flow cell.

##### Analysis of single-cell RNA-seq data

Reads from each single-cell gene expression library were demultiplexed, aligned to the genome reference GRCm39 (pre-built reference “2024-A”) and collapsed into gene-barcode matrices containing Unique Molecular Identifier (UMI) counts using cellranger count pipeline from 10x Genomics’ Cell Ranger Software Suite (v9.0.1)^74^, indicating an expectation of 20,000 cells per sample. Filtered feature-barcode matrices were imported into R statistical environment (v 4.4.3) for analysis based on Seurat (v5.3.0)^75^. Seurat objects were created for each sample, genes expressed in at least 1 cell and cells expressing at least one gene from each dataset were kept for analysis.

Putative doublets were identified and removed using the DoubletFinder (v2.0.6)^76^ package. Each Seurat object was first normalized and subjected to PCA prior to doublet estimation for the purpose of computing doublet simulations. Expected doublet rate was defined to 8% based on cell recovery expectancy, defined by 10xGenomics protocol for Chromium GEM-X Single Cell 3’ Reagent Kits v4. Artificial doublets were generated and classified using paramSweep() and doubletFinder() functions, following the developers’ recommendations. Cells predicted as doublets were excluded from subsequent analyses.

Further, potential remaining multiplets as well as low-quality or stressed cells were removed by computing quality control (QC) feature cutoffs, we established cutoffs above the 10th percentile and below the 99th percentile for the total UMI counts per cell, above 10th percentile for the number of detected genes per cell, and below 10% of mitochondrial transcript content. Metrics were tailored according to the predefined cell populations to account for cell type variability in the distribution of each of their raw QC features. The resulting high-quality singlet cells were retained for downstream analyses.

Following QC, normalization and variance stabilization were performed using SCTransform^77^, which models UMI counts using Pearson residuals from a negative binomial regression. This method effectively removes technical noise and accounts for sequencing depth while preserving biological variability. Mitochondrial percentage was regressed out to correct for cell quality-related variation. Highly variable genes identified by SCTransform were used for all downstream dimensional reduction and clustering analyses.

We identified cell clusters representing subpopulations of each predefined cell population having cells with similar transcriptomic profiles by computing Euclidean distances across genes. First, we performed principal component analysis (PCA) using RunPCA() on the scaled SCT assay, and then applying the function FindNeighbors() which will construct a KNN graph based on the euclidean distance in PCA space, and refine the edge weights based on the shared overlap in their local neighborhoods, this then informs FindClusters() (provided by 30 principal components (PCs)), which uses unsupervised modularity optimization with the Louvain algorithm to group multiple data points into clusters based on their similarities. Uniform Manifold Approximation and Projection (UMAP) embeddings were calculated with the function RunUMAP() for visualization of clustering. Clusters were annotated based on marker genes identified using FindAllMarkers() and reference known cell-type markers from literature.

##### Transcriptional variance analysis with BASiCS

To assess transcriptional variability between conditions, we employed BASiCS (Bayesian Analysis of Single-Cell Sequencing data) (v2.18.0)^78^, which implements an integrated Bayesian hierarchical model that performs built-in normalization and quantifies both technical and biological variability.

For each condition (Lif/2i siCTRL, Lif/2i siAQR, EpiLC siCTRL, EpiLC siAQR) genes were filtered out correspondently if they were expressed in < 1% of the cells from the respective condition. We ran the BASiCS_MCMC() sampler using the following parameters: N=20,000, Thin=10, Burn=10,000. We employed the regression-based implementation (Regression=TRUE) to derive residual measures of over-dispersion that are not confounded by mean expression levels.

Differential expression analysis between siCTRL and siAQR from correspondent cell types was performed using the BASiCS_TestDE() function with the following customized thresholds: EFDR_M=0.1, EFDR_R=0.1. Genes were classified as differentially expressed if they exceeded the posterior probability threshold (determined by EFDR calibration) for changes in mean expression. Additionally, we identified genes with differential variability based on changes in [biological over-dispersion (delta) / residual over-dispersion (epsilon)].

##### Transcriptional Entropy and Stemness inference with IDEAS

To quantify cellular differentiation potential and stemness, we calculated transcriptional entropy using IDEAS (Intrinsic Dimensionality Estimation Analysis of single-cell RNA sequencing data)^39^. IDEAS computed single-cell intrinsic dimensionality (ID) from the scRNA-seq raw counts matrix, providing a data-driven, unbiased measure of transcriptional complexity that correlates with developmental potential. The approach is based on the principle that pluripotency or stem-like cells exhibit higher transcriptional promiscuity and diversity (higher entropy), while differentiated cells display more restricted and specialized gene expression profiles (lower entropy). Local intrinsic dimensionality was estimated for each cell using the compute_ID() function with the parameters: method=”local_2nn”, n_neighbors=200, id_score=True, capturing the complexity of the local transcriptional landscape.

## Acknowledgments

We thank current and former members of the Hamperl laboratory. The SH laboratory was supported by the Helmholtz association, by the Deutsche Forschungsgemeinschaft (project IDs 552129721 and 213249687 (CRC1064)) and by ERC Starting Grant no. 852798 – ConflictResolution. Work in the Scialdone lab is funded by the Helmholtz Association. The funders had no role in study design, data collection and analysis, decision to publish, or preparation of the manuscript. We would like to thank Nadine Übelmesser for providing us with human induced pluripotent stem cells. We would like to thank Dr. Giacomo Masserdotti and Tatiana Ebert-Simon for their expertise, help, and support with scRNA-seq library preparation.

## Author contributions

Conceptualization: SH, METP, AS

Methodology: ML, EMG, AB, FRB, ACM, AE, AS

Investigation: ML, EMG, AB, FRB, TS, TS, ACM

Visualization: ML, EMG

Funding acquisition: SH, AS, METP

Project administration: SH, AS, METP

Supervision: SH, AS, METP

Writing – original draft: ML, SH

Writing – review & editing: ML, EMG, AB, CSKL, IT, MW, AS, FRB, ACM, TS, TS, XK, AE, METP, SH

## Data and materials availability

bulk RNA-Seq and scRNA-Seq data have been deposited at the European Nucleotide Archive under access number PRJEB101486. All other data are available in the main text or the supplementary materials.

## Supplementary Tables

Supplementary Table 1. List of TRC factors

Supplementary Table 2. List of R-loops regulators

Supplementary Table 3. List of DNA Damage Response factors

Supplementary Table 4. List of Epigenetics factors

Supplementary Table 5. mESC 2CLC DEA (from publicly available datasets)

Supplementary Table 6. mESC vs MEF DEA (from publicly available datasets)

Supplementary Table 7. mESC EpiLC DEA (from publicly available datasets)

Supplementary Table 8. Bulk RNA-seq results (TPMs)

Supplementary Table 9. Differential Transcriptional Variance analysis from BASiCS (naive mESCs (Lif/2i))

Supplementary Table 10. Differential Transcriptional Variance analysis from BASiCS (primed mESCs (EpiLCs))

---

**Figure S1.**
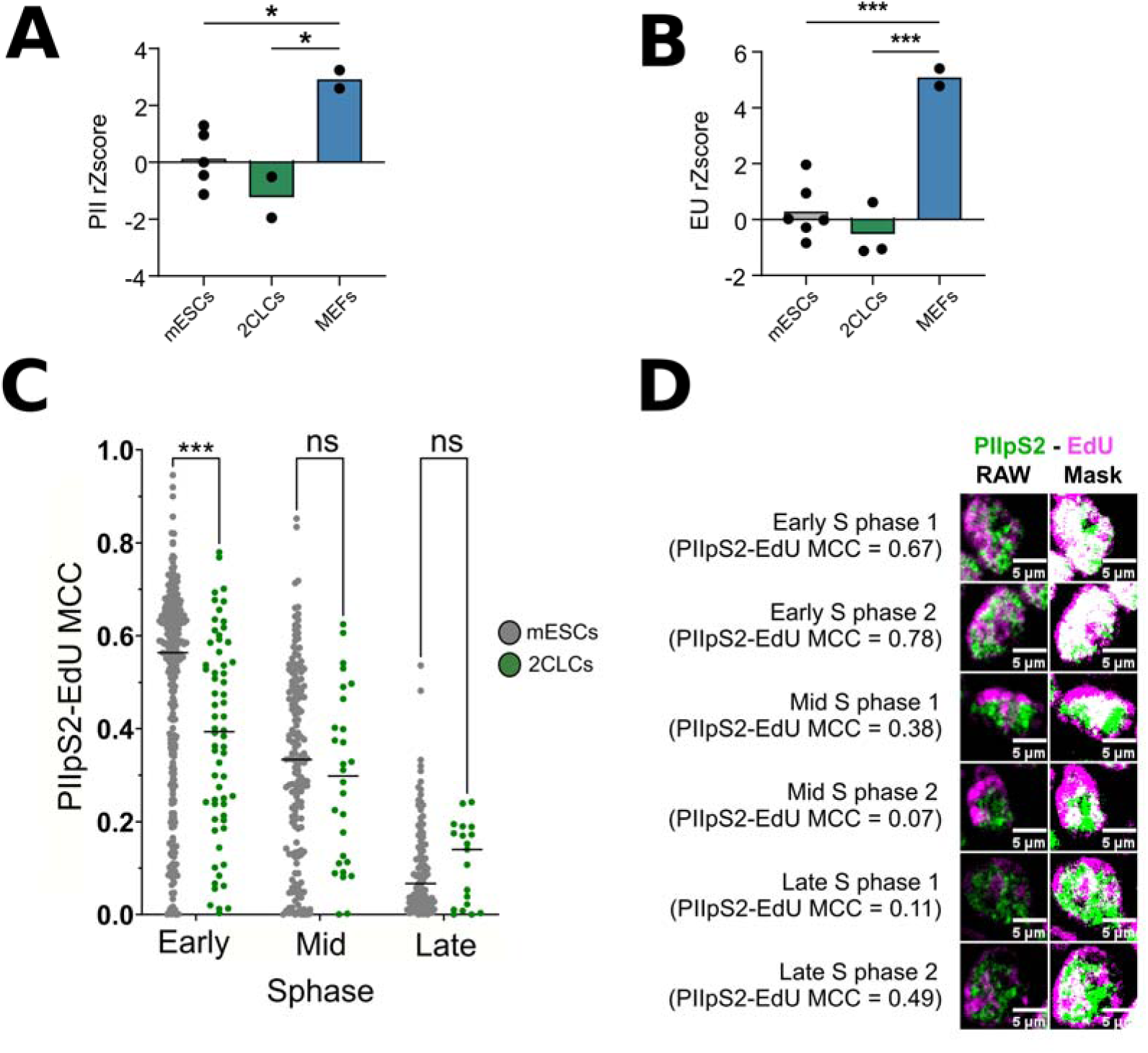
Technical and Biological Controls for T-R coordination quantification. (**A**) Quantification of total PII across cell types. PII signal was normalized replicate-wise through robust Z score transformation. Each dot represents one biological replicate (N = 2-5), calculated as the mean of ≥3 technical replicates, except for 2CLCs (1300–5,300 analyzed mESCs or MEFs per technical replicate). For 2CLCs, each biological replicate corresponds to the mean of all 2CLCs within that biological replicate (≥20 2CLCs per replicate). Statistical significance was determined by one-way ANOVA followed by Bonferroni’s multiple comparisons test. (**B**) Quantification of EU incorporation across cell types. EU signal was normalized replicate-wise through robust Z score transformation. Each dot represents one biological replicate (N = 2-6), calculated as the mean of ≥3 technical replicates, except for 2CLCs (1300–5,300 analyzed mESCs or MEFs per technical replicate). For 2CLCs, each biological replicate corresponds to the mean of all 2CLCs within that biological replicate (≥20 2CLCs per replicate). Statistical significance was determined by one-way ANOVA followed by Bonferroni’s multiple comparisons test. (**C**) Quantification of transcription–replication coordination in mESCs (grey) and 2CLCs (green) across S-phase progression zones, using PIIpS2–EdU MCC. S-phase zones were manually defined by EdU patterns^19^. Statistical significance was determined by one-way ANOVA followed by Bonferroni’s multiple comparisons test. (**D**) Representative images of mESCs in different S-phase zones. Left panels: raw PIIpS2 and EdU signals. Right panels: binarized masks of PIIpS2 and EdU signals to emphasize the overlap (white) quantified by the MCC. Two examples per zone are shown.

**Figure S2.**
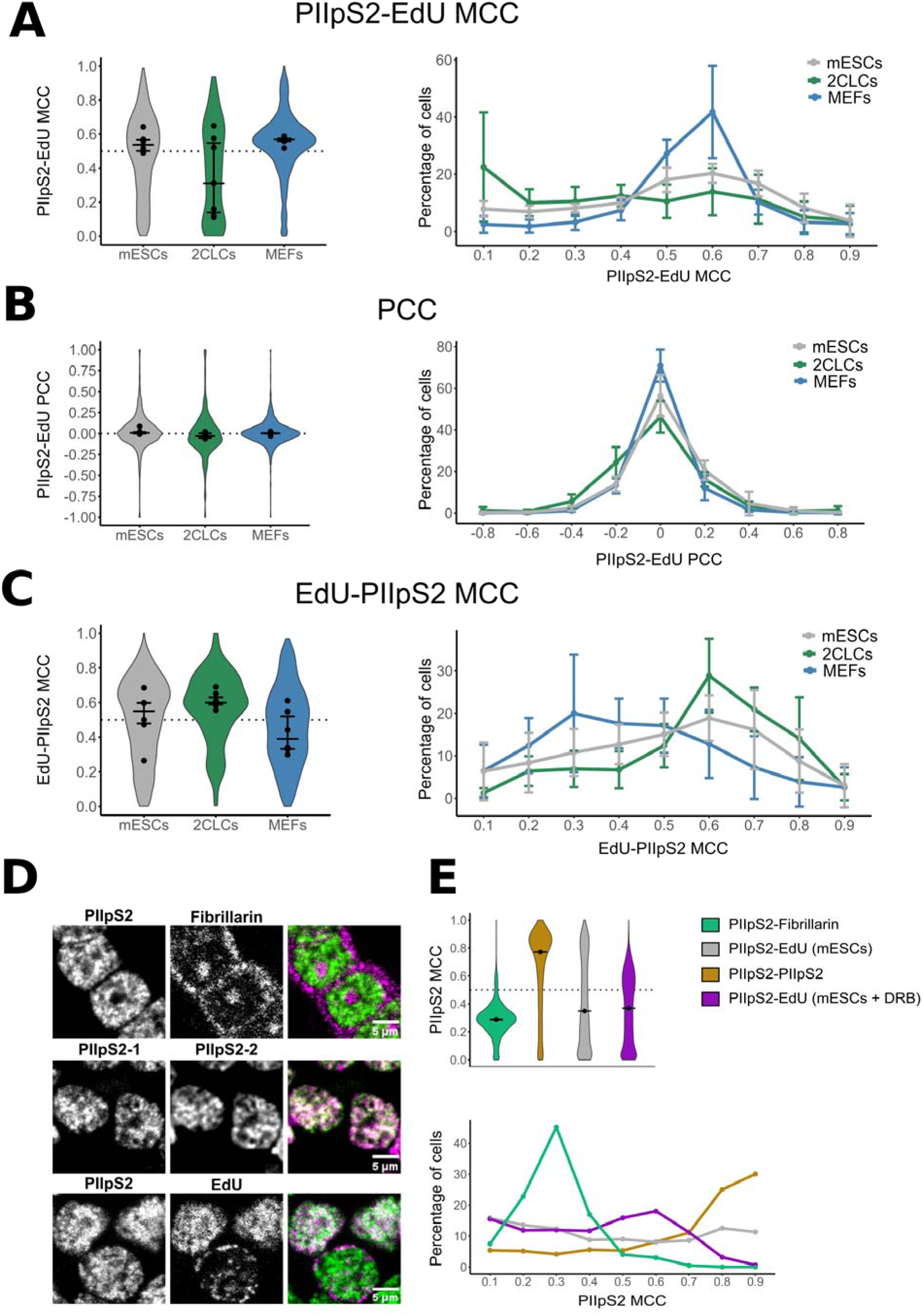
Comparison of different coefficient for the quantification of transcription–replication coordination. Quantification of transcription–replication coordination using (**A**) PIIpS2-EdU MCC, (**B**) PCC or (**C**) EdU-PIIpS2 MCC,across cell types using the same datasets used in Fig. 2A. Violin plots (left) and frequency distributions (right) show MCC distributions. Dots in the violin plots represent the mean value of biological replicates, the violin plot shows the distribution of all biological replicates pooled together. The frequency distributions show the mean and standard deviation of all biological replicates for each bin of MCC. Each cell type has been down sampled to the condition with the lowest number of cells for each independent experiment. N = 5 (7 for 2CLCs), comprising 464 2CLCs, 4097 mESCs, and 3178 MEFs. (**D**) Representative immunofluorescence images for MCC controls. Anticorrelation with corresponding low MCC values was measured using PIIpS2 and the nucleolar marker Fibrillarin (top); Colocalization with corresponding high MCC values was measured using two antibodies recognizing PIIpS2 (middle). PIIpS2–EdU MCC in mESCs shows heterogeneous distribution (bottom). The right panels show both channels, with the overlap appearing as white. (E) Quantification of MCC for the conditions in (**C**), including mESCs treated with 100 µM DRB for 1 h. Violin plots (left) and frequency distributions (right) show MCC distributions.

**Figure S3.**
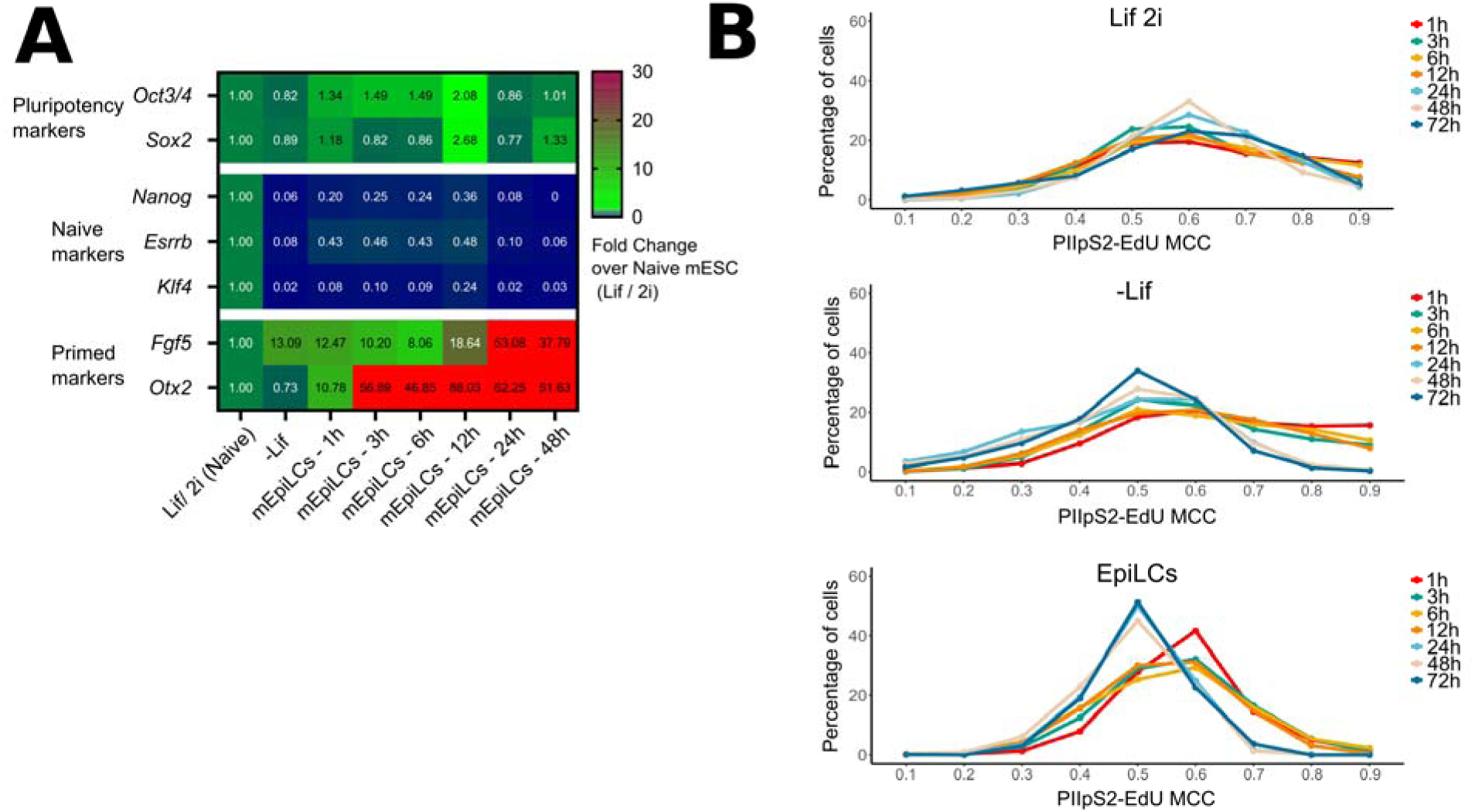
Complementary analyses of transcription–replication coordination and naive-to-primed transition. (**A**) RT–qPCR validation of the naive-to-primed differentiation protocol. Cells were processed as in Fig. 3A (right). RNA was extracted at the indicated timepoints, and relative expression levels were quantified using the ΔΔCt method, normalized to *Actinβ*, *Gapdh*, and *Rpl13a*, with naive cells (Lif 2i) as reference. Priming efficiency was assessed by sustained expression of pluripotency genes, downregulation of naive markers, and induction of primed markers. (**B**) Distribution of PIIpS2–EdU MCC values measuring transcription–replication coordination during cell fate transition time courses (as in Fig. 3A).

**Figure S4.**
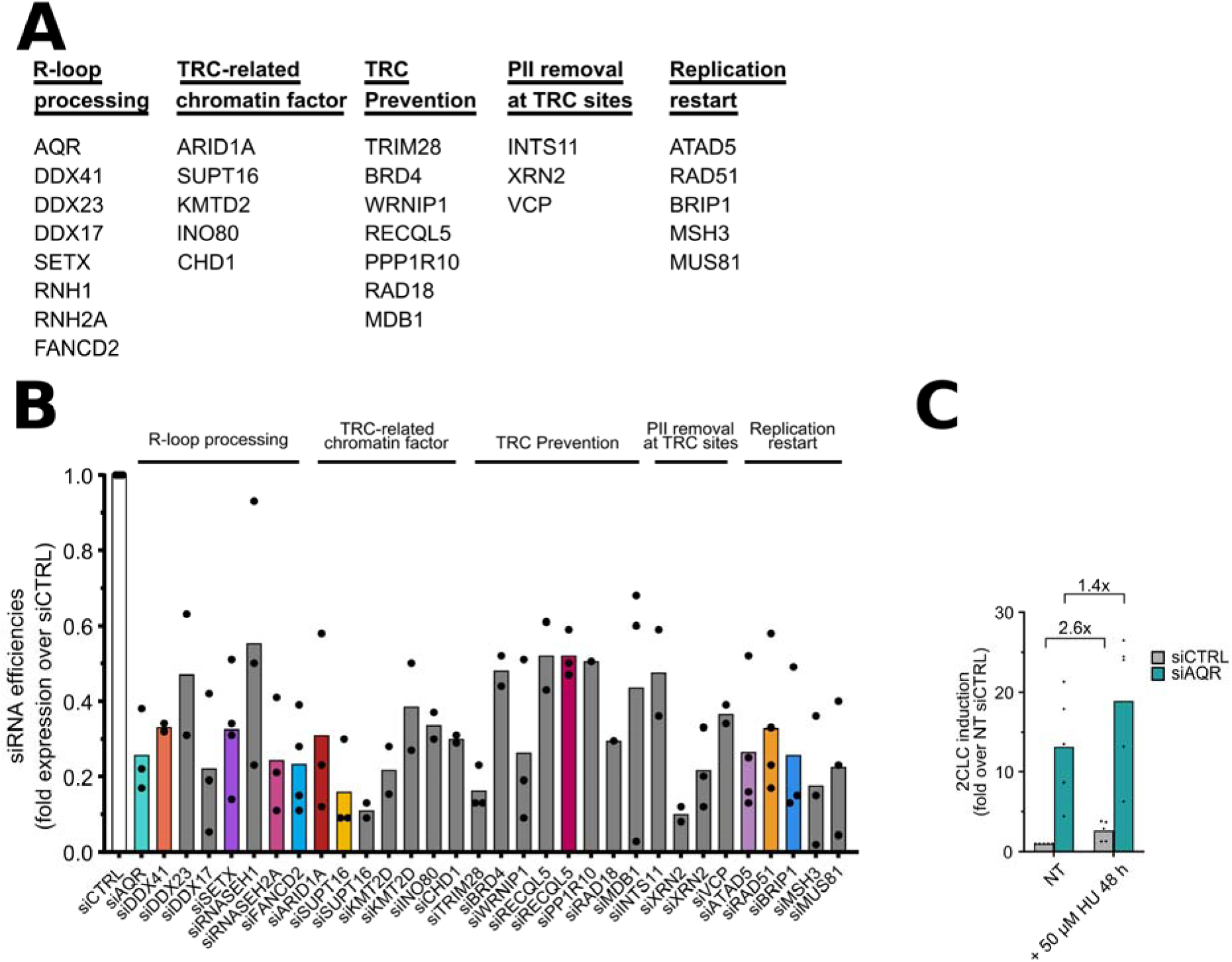
qPCR validation of siRNA KDs and additional 2CLC emergence characterizations. (**A**) List of all factors used for the siRNA screen shown in Fig. 4, grouped by functional category. (**B**) RT–qPCR validation of KD efficiency for the siRNA subsets shown in Fig. 3. Cells were transfected with siRNAs, cultured for 48 h, and RNA extracted. Relative expression was calculated using the ΔΔCt method, normalized to *Actinβ*, *Gapdh*, and *Rpl13a*, with siCTRL as reference. (**C**) Evaluation of the added impact of hydroxyurea (HU) treatment during AQR KD on 2CLC emergence, as measured by FACS. HU was added at a 50 µM concentration for the 48 h duration of the siRNA KD (N=4). Statistical significance was determined by paired two-way ANOVA followed by Bonferroni’s multiple comparisons test.

**Figure S5.**
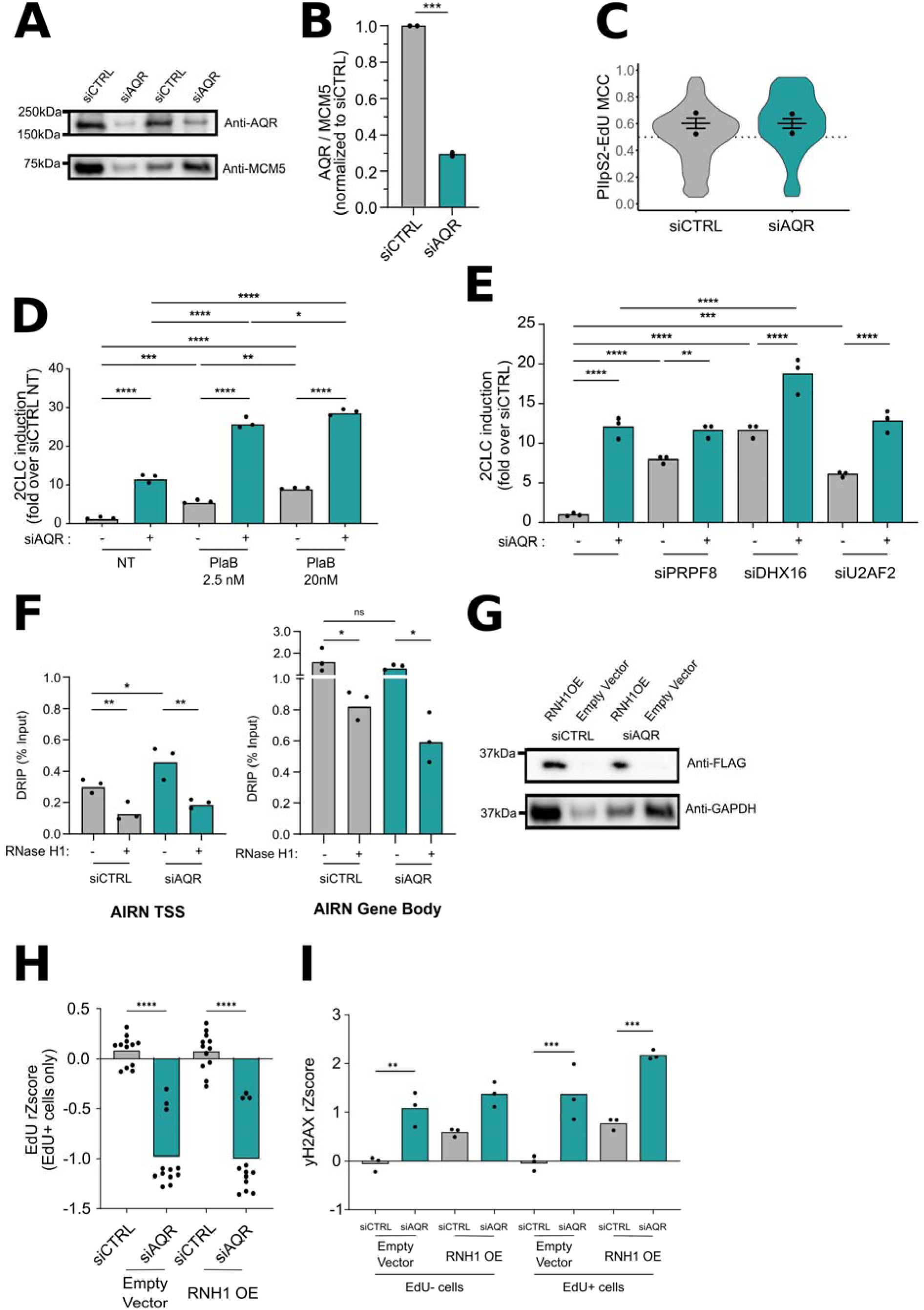
Complementary analyses of AQR KD phenotype. (**A**) Representative Western blot showing depletion of AQR protein following siRNA-mediated KD. AQR protein levels are shown in the upper panel (marked by the **\***), with MCM5 as loading control in the lower panel. (**B**) Quantification of AQR protein KD relative to siCTRL. Band intensities were normalized to MCM5 and expressed as fold change over siCTRL (N=2). Statistical significance determined by a two-tailed paired t-test. (**C**) PIIpS2–EdU MCC violin plots showing the effect of siAQR on transcription–replication coordination. Dots in the violin plots represents the mean value of biological replicates, the violin plot shows the distribution of all biological replicates pooled together (N = 2; 50–1,000 analyzed cells per replicate). (**D**) Evaluation of the additive effect of a 4h pulse of Pladienolide B (PlaB) (2.5nM or 20 nM) during AQR KD on 2CLC emergence, as measured by FACS (N=3). Statistical significance determined by a two-way ANOVA followed by followed by a Bonferroni’s multiple comparisons test. (**E**) Evaluation of the additive effect of additional knockdown of splicing factors with AQR KD on 2CLC emergence, as measured by FACS (N=3). Statistical significance determined by a two-way ANOVA followed by followed by a Bonferroni’s multiple comparisons test. (**F**) DRIP-qPCR analysis quantifying R-loops accumulation upon AQR KD at the R-loop prone *Airn* gene at the TSS (left) and in the gene body (right). *E. coli* RNaseH1 treatments were added to verify the specificity of the immunoprecipitation (N=3). Statistical significance determined by a one-way ANOVA followed by followed by a Bonferroni’s multiple comparisons test. (**G**) Representative Western blot showing overexpression of hRNASEH1-FLAG. hRNASEH1-FLAG protein levels were detected using an antibody against FLAG and are shown in the upper panel (marked by the **\***), with GAPDH as loading control in the lower panel. (**H**) Effect of hRNASEH1 OE on EdU incorporation in EdU-positive cells in siCTRL and siAQR transfected cells, normalized replicate-wise by robust Z score transformation. Each dot represents the mean value of a technical replicate of 2 biological replicates. Dot shape represents biological replicates (160–2,942 analyzed cells per technical replicate). Statistical significance was determined by two-way ANOVA followed by Bonferroni’s multiple comparisons test. (**I**) Effect of hRNASEH1 OE on DNA damage in EdU-negative and EdU-positive cells in siCTRL and siAQR transfected cells. DNA damage quantified by yH2AX levels, normalized by robust Z-score transformation. Each dot represents a technical replicate (N =1) (160–2,942 analyzed cells per technical replicate). Statistical significance determined by a two-way ANOVA followed by followed by a Bonferroni’s multiple comparisons test performed on the technical replicates.

**Figure S6.**
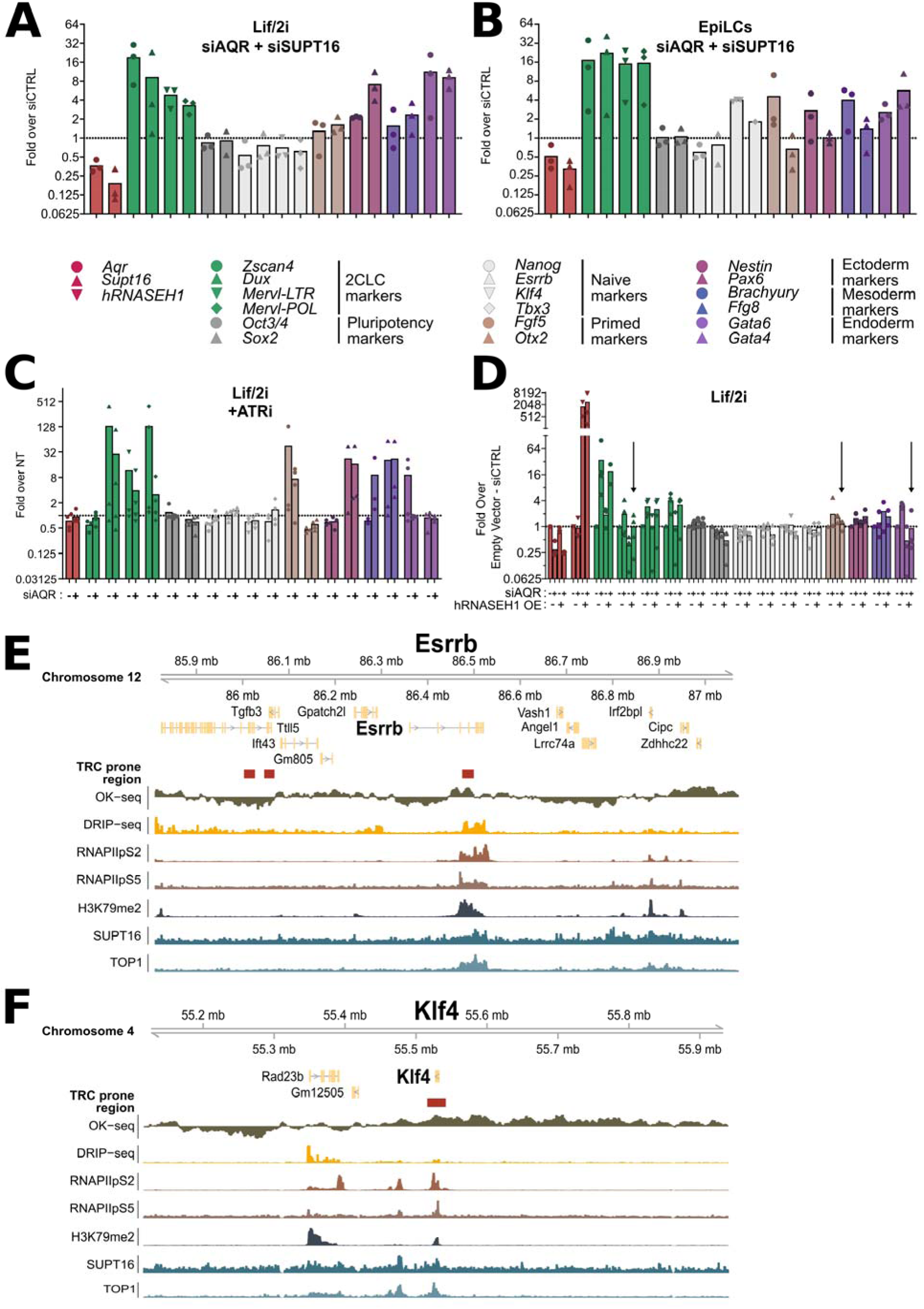
Complementary analyses of AQR KD effects on pluripotency maintenance. (**A-B**) RT–qPCR analysis of cell fate–related genes following siRNA-mediated co-depletion of AQR and SUPT16 in mESCs maintained under naive conditions (Lif/2i) (**A**) or during priming of mESCs to the EpiLC state (**B**). Cells were transfected with siCTRL or a combination of siAQR and siSUPT16 and cultured for 48 h before RNA extraction. For EpiLCs, the siRNA transfection was performed concomitantly with the switch to priming media. Relative gene expression was calculated by the ΔΔCt method, normalized to *Actinβ*, *Gapdh*, and *Rpl13a*, with siCTRL as reference (N=3). (**C**) RT–qPCR analysis of cell fate–related genes upon ATR inhibition (VE-821, 10 µM, 4 h) during AQR KD. mESCs were transfected with siCTRL or siAQR and cultured for 48 h in Lif/2i. ATRi was applied during the final 4 h before RNA extraction. Relative expression was calculated as in (**A**) (N=3). (**D**) RT–qPCR analysis of cell fate–related genes upon overexpression of human RNASEH1 during AQR knockdown. Relative expression was calculated as in (A) (N = 3–4). Arrows indicate genes for which hRNASEH1 OE antagonizes the effect of siAQR. (**E-F**) Genomic snapshots of pluripotency genes Esrrb (**E**) and Klf4 (**F**) showing their TRC related genomic context, with Essrb showing a propensity for co-directional TRCs (**E**), and Klf4 showing a propensity for head-on TRCs (**F**). Tracks shown from top to bottom include: predicted TRC-prone regions defined using a TRC prone region prediction framework developed by Aguilera and colleagues ^38^ OK-seq; DRIP-seq; RNAPII-pS2 ChIP-seq; RNAPII-pS5 ChIP-seq; H3K79me2 ChIP-seq; SUPT16 ChIP-seq; and TOP1 ChIP-seq.

**Figure S7.**
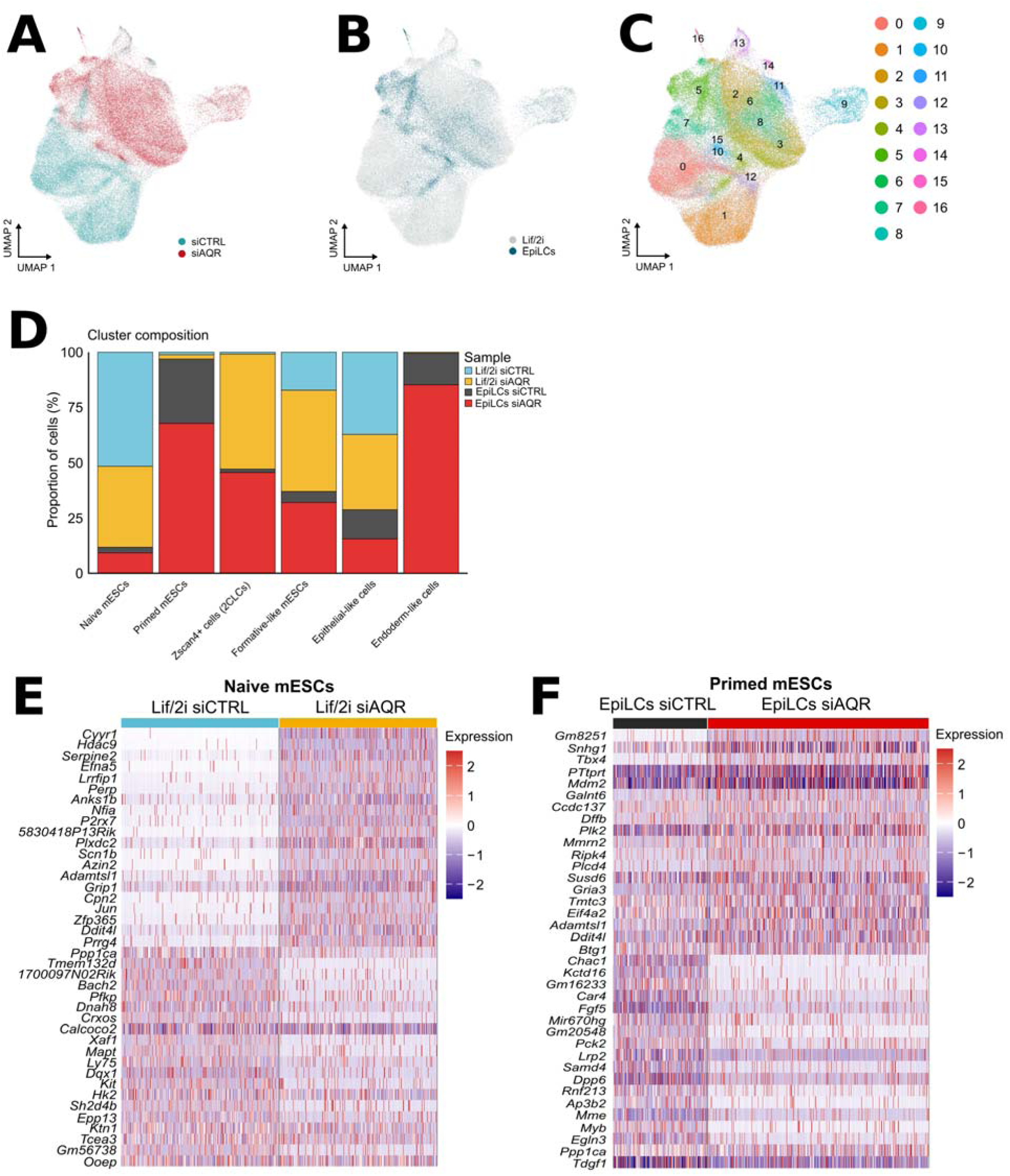
Differential expression in AQR-depleted naive and primed mESCs. **(A)** UMAP visualization of single-cell transcriptomes showing the distribution of siCTRL and siAQR cells (**B**) UMAP visualization of single-cell transcriptomes showing the distribution of naive (Lif/2i) and primed (EpiLC) conditions. (**C**) UMAP visualization of single-cell transcriptomes showing the distribution of 17 cell clusters identified by Unsupervised Louvain clustering. (**D**) Characterization of the contribution of the different samples to the annotated cell lineage and pluripotency clusters. (**E**) Differential gene expression between siCTRL and siAQR within the naive mESCs cluster under Lif/2i conditions. **(F)** Differential gene expression between siCTRL and siAQR within the primed mESCs cluster under EpiLCs conditions.

**Figure S8.**
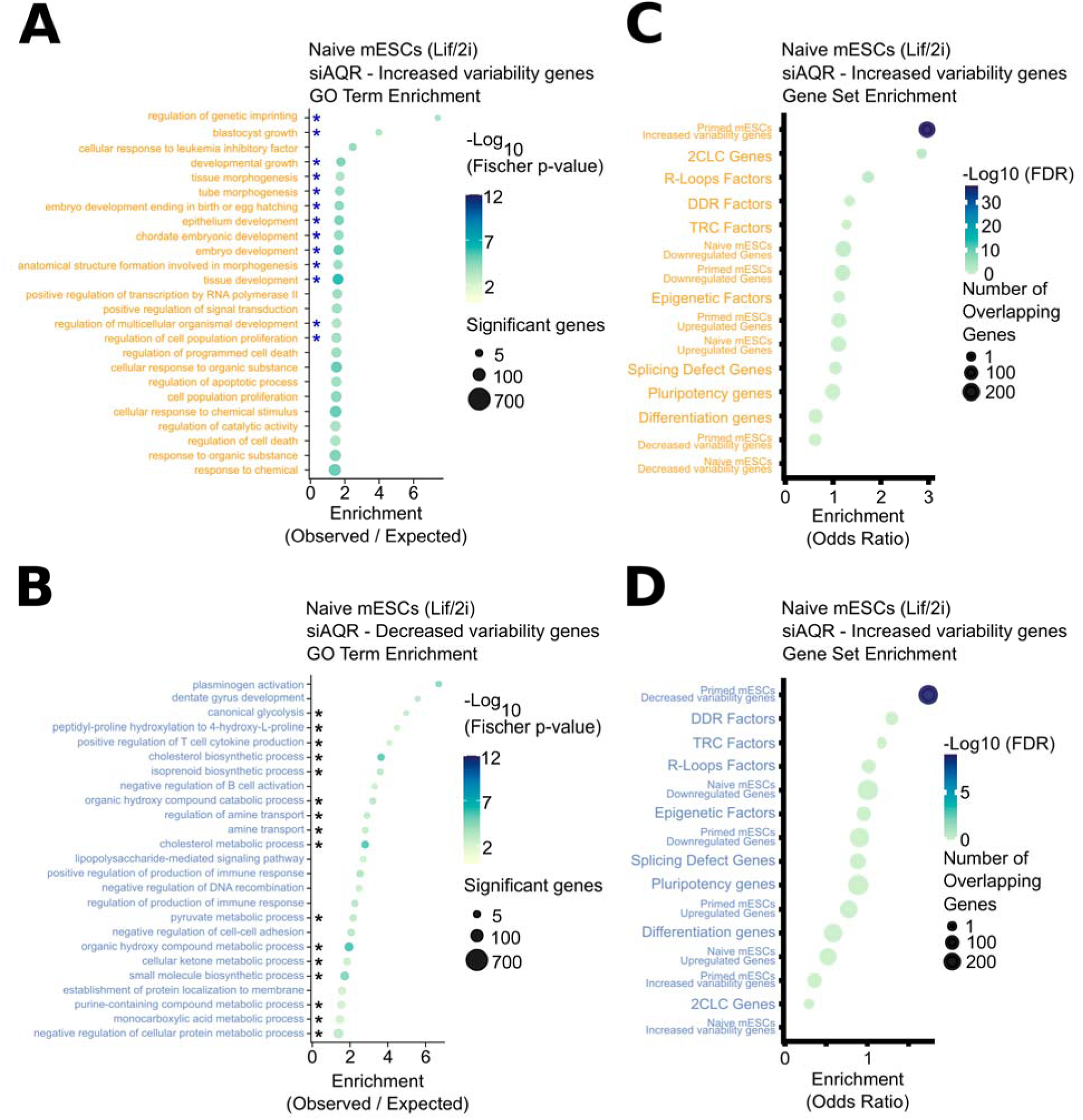
Functional enrichment analysis of genes with altered transcriptional variability upon AQR depletion in naive mESCs. **(A-B)** GO biological process enrichment of genes showing altered variability upon AQR depletion in naive mESCs (Lif/2i). Dot plots show the top 25 enriched terms for increased variability genes (**A**) and decreased variability genes (**B**). Dot size indicates the number of significant genes per term, and color reflects statistical significance (−log₁₀ Fisher p-value). Enrichment is expressed as the observed/expected gene ratio. Terms were selected based on stringent statistical criteria (Fisher.classic < 1×10⁻³ and Fisher.elim < 0.05). Terms attributed to genomic imprinting and developmental processes are highlighted with a blue star and terms attributed to metabolic pathways are highlighted with a black star. (**C-D**) Targeted Gene Set Overlap analysis of genes showing altered variability upon AQR depletion in naive mESCs (Lif/2i). Dot plots show the enrichment of the increased variability genes (**C**) and of decreased variability genes (**D**) against multiple curated gene lists. Dot size represents the number of overlapping genes between the query and target lists, color indicates statistical significance (−log₁₀ FDR), and the x-axis shows the odds ratio as a measure of enrichment. The curated gene lists tested are the lists of TRC resolution factors, R-loop regulators, DDR factors, and epigenetic regulators used in Fig. 2D, S3B, and **S4C**. Additionally, DEGs were tested against the list of genes showing splicing defects upon AQR depletion and against genes linked to pluripotency maintenance, differentiation, and transition to 2CLCs.

**Figure S9.**
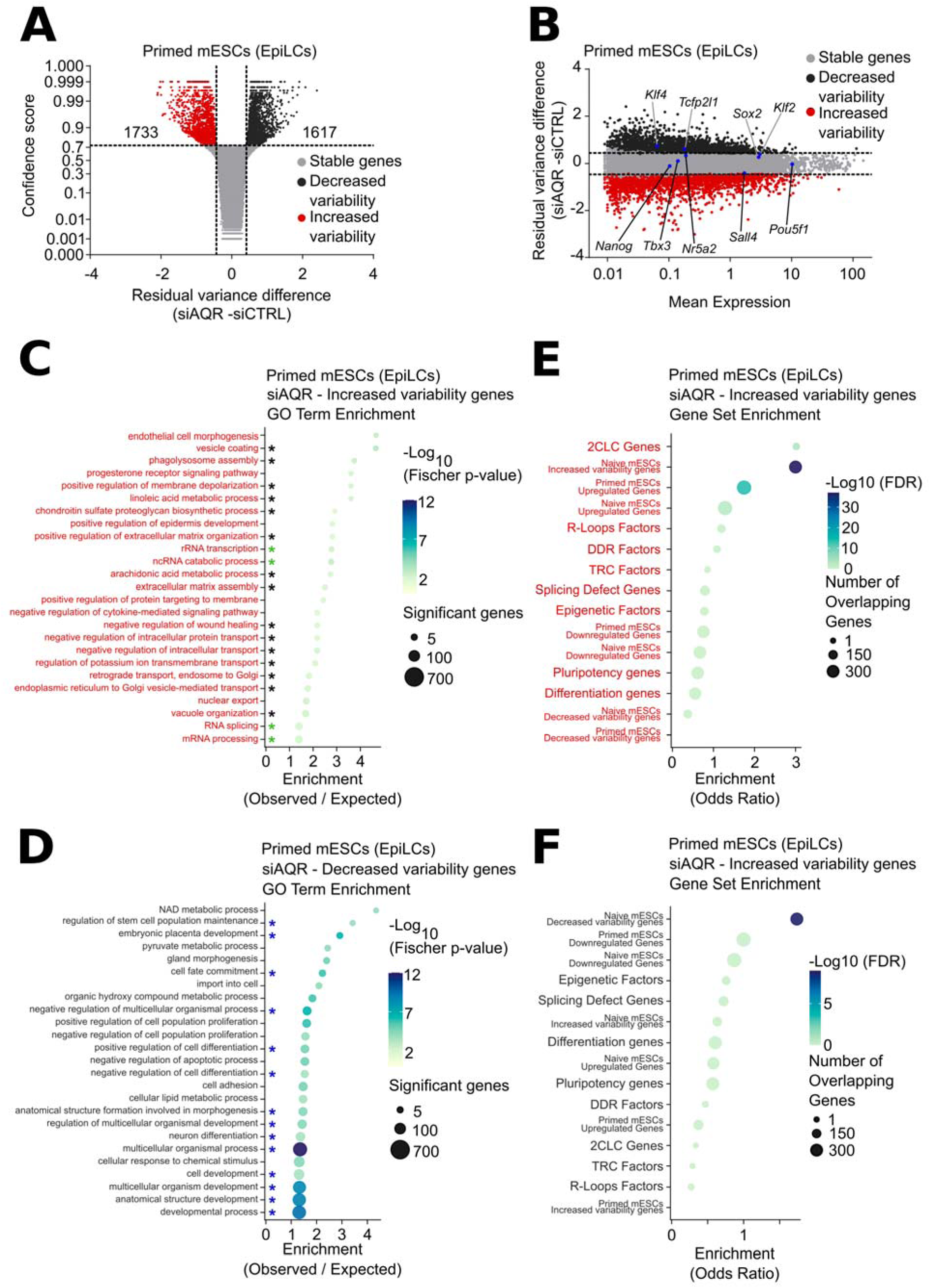
Functional enrichment analysis of genes with altered transcriptional variability upon AQR depletion in primed mESCs. **(A)** Differential transcriptional variance in primed mESCs, plotted as confidence score versus residual variance difference (siAQR - siCTRL). Dashed lines indicate thresholds for statistical significance. Numbers denote the total genes exhibiting significantly decreased variability (black) or increased variability (red) upon AQR depletion. (**B**) Differential transcriptional variance in primed mESCs represented as residual variance difference (siAQR - siCTRL) versus mean gene expression. Highlighted genes in blue correspond to key pluripotency regulators. (**C-D**) GO biological process enrichment of genes showing altered variability upon AQR depletion in primed mESCs (Epi/LCs). Dot plots show the top 25 enriched terms for increased variability genes (**C**) and decreased variability genes (**D**). Dot size indicates the number of significant genes per term, and color reflects statistical significance (−log₁₀ Fisher p-value). Enrichment is expressed as the observed/expected gene ratio. Terms were selected based on stringent statistical criteria (Fisher.classic < 1×10⁻³ and Fisher.elim < 0.05). Terms attributed to developmental processes are highlighted with a blue star, terms attributed to metabolism and cell morphology are highlighted with a black star, and terms attributed to transcription regulation are highlighted with a green star. (**E-F**) Targeted Gene Set Overlap analysis of genes showing altered variability upon AQR depletion in primed mESCs (EpiLCs). Dot plots show the enrichment of the increased variability genes (**E**) and of decreased variability genes (**F**) against multiple curated gene lists. Dot size represents the number of overlapping genes between the query and target lists, color indicates statistical significance (−log₁₀ FDR), and the x-axis shows the odds ratio as a measure of enrichment. The curated gene lists tested are the lists of TRC resolution factors, R-loop regulators, DDR factors, and epigenetic regulators used in Fig. 2D, S3B, and **S4C**. Additionally, DEGs were tested against the list of genes showing splicing defects upon AQR depletion and against genes linked to pluripotency maintenance, differentiation, and transition to 2CLCs.

